# Inherent instability of simple DNA repeats shapes an evolutionarily stable distribution of repeat lengths

**DOI:** 10.1101/2025.01.09.631797

**Authors:** Ryan J. McGinty, Daniel J. Balick, Sergei M. Mirkin, Shamil R. Sunyaev

## Abstract

Using the Telomere-to-Telomere reference, we assembled the distribution of simple repeat lengths present in the human genome. Analyzing over two hundred mammalian genomes, we found remarkable consistency in the shape of the distribution across evolutionary epochs. All observed genomes harbor an excess of long repeats, which are potentially prone to developing into repeat expansion disorders. We measured mutation rates for repeat length instability, quantitatively modeled the per-generation action of mutations, and observed the corresponding long-term behavior shaping the repeat length distribution. We found that short repetitive sequences appear to be a straightforward consequence of random substitution. Evolving largely independently, longer repeats (above roughly 10 nt) emerge and persist in a rapidly mutating dynamic balance between expansion, contraction and interruption. These mutational processes, collectively, are sufficient to explain the abundance of long repeats, without invoking natural selection. Our analysis constrains properties of molecular mechanisms responsible for maintaining genome fidelity that underlie repeat instability.

## Introduction

Over 2.5% of human genomic DNA consists of simple DNA repeats^1^. Also known as short tandem repeats (STRs) or microsatellites, simple repeats consist of direct tandem repetitions of short sequence motifs, e.g., mononucleotides, dinucleotides, trinucleotides and so forth. In a randomized DNA sequence, the probability of encountering a simple repeat is exponentially decreased with increasing tract length. Yet this relationship fails to predict the enormous overrepresentation of long simple repeats in most genomic sequences, including in humans^2,3,4^. The origin of this overrepresentation remains to be elucidated.

This overrepresentation is even more striking in light of the existence of repeat expansion disorders, a growing list of severe human diseases caused by disruption of gene function due to long STRs^5,6^. Decades of study have demonstrated that repeat tract lengths vary between and within individuals^7^, owing to frequent expansion and contraction mutations. The rate of these mutations increases with the length of a repeat, a phenomenon known as repeat length instability^8^. Length instability is commonly ascribed to DNA strand slippage during replication and/or DNA repair, although a variety of other molecular mechanisms can also contribute8. Instability rates differ between various repeat motifs, being particularly pronounced for motifs that form non-B DNA secondary structures^9^. Importantly, when repeat length exceeds a threshold of approximately 75-90 nt, carriers frequently transmit a substantially longer repeat to the next generation. Known as ‘genetic anticipation’, this effect continues to compound in subsequent generations, which leads to more severe presentation and/or earlier age of onset^6^. Recently developed techniques such as ExpansionHunter^10^ and long-read sequencing have accelerated the discovery of pathogenic repeats; in particular, the growing number of repeat expansion disorders mapped to introns and other non-coding regions sheds light on repeat disease biology beyond coding regions. Repeat expansions are also observed in various cancers^11,12,13^ and serve as hotspots for genomic rearrangements^14^.

While numerous studies focus on the instability of disease-length repeats, comparatively less is known about shorter repeats, including the so-called ‘long-normal’ alleles that sit immediately below the disease-length threshold. Carriers of long-normal repeat alleles are healthy, but risk transmitting a disease-length allele due to the higher rate of repeat expansion; additionally, some ‘long-normal’ alleles contain protective interruptions that, if lost, result in reversion to disease length^6^. Complementing our understanding of long disease-causing repeats, a recent finding identified an autosomal dominant thyroid disorder linked to a (TTTG)_4_ repeat, with a recurrent deletion to (TTTG)_3_ in affected individuals^15^. Additionally, instability of A_8_ and C_8_ repeats in the coding sequences of mismatch repair genes MSH3 and MSH6, respectively, promotes tumor adaptability via frequent frameshifts and subsequent reversions^16^. The latter examples suggest that relatively short repeats, which comprise a much larger portion of the genome, also have biomedical relevance.

In light of the rapidly growing list of repeat-associated diseases, it is surprising to find repeats harbored in abundance in the genome. Interest in this discrepancy goes back at least three decades^2^ and has led to speculation that natural selection preserves longer repeat lengths despite the risk of disease^17^. The best-supported examples of functionality are specific to telomeric and centromeric repeats^9,17,18^, though some recent studies have suggested that simple repeats play a role in gene regulation^17^. However, before assuming the overabundance of repeats is evidence of functionality, a more basic explanation should be considered: the excess of repeats in the genome is solely a consequence of mutational processes. Several studies, largely pre-dating the human genome era, considered this premise, but were limited by the availability of sufficiently long genome sequences, lacked robust direct measurements of repeat instability, and/or considered oversimplified mutational models^4,19,20,21,22,23,24,25,26,27,28,29,30,31,32,33,34^. Indeed, all such studies of simple repeats have been limited by long-standing technical challenges to sequencing repetitive regions^35,36,37,38^. Technological developments led to the release of the human Telomere-to-telomere genome (T2T-CHM13), which more than doubled the number of mapped simple repeats compared to the previous reference genome GRCH38^1^. This warranted a fresh look at the distribution of repeat lengths and whether mutational processes, in the absence of selection, can explain their abundance.

In this study, we measured genome-wide distributions of repeat lengths across mammals, observing that the distribution, including the prevalence of long repeats, is remarkably stable over evolutionary timescales. We modeled the effects of repeat length instability on evolution of the distribution, finding that the observed repeat length distribution can emerge and be maintained solely due to the interplay between distinct mutational processes. After incorporating empirical estimates and inference of repeat length instability rates, the most parsimonious explanation for the abundance and stability of long repeats does not require invoking selection; rather, extreme mutation rates cause long repeats to emerge as independently evolving elements. We discuss how inherent constraints of DNA replication and repair machineries could lead to persistent repeat length polymorphism.

## Results

### Features of the repeat length distribution and evolutionary stability

Using T2T-CHM13, we first assembled a genome-wide Distribution of Repeat tract Lengths (henceforth, DRL) for each simple tandem repeat motif, pooling over bioinformatically indistinguishable permutations (see **Fig. S1a, Methods**). Each distribution was assembled by counting contiguous, uninterrupted repetitions of a specified motif, allowing for straightforward bioinformatic assembly of the DRL (see **Methods**). Each DRL showed a marked excess of repeats longer than ∼10 nt., relative to a randomly shuffled control. This was apparent for nearly all motifs (**Fig. S1b**) but with motif-specific variation in the shape of the extended tail of long tract lengths. **Fig. 1a** plots the DRLs after pooling motifs of the same unit length (e.g., mononucleotide repeats, dinucleotide repeats, etc. up to hexamer repeats), each with a clear tail of long repeats. We found that short read sequencing was sufficient to reconstruct the well-populated length classes of nearly all DRLs, lacking estimates only for the very longest repeats (**Fig. S1c**). We were therefore able to estimate distributions from genome sequences of over 300 mammals from the Zoonomia project^39^ and compare them to humans. Due to differences in total assembly length, direct comparison was performed on the normalized DRL (see **Fig. S2, Methods**). There was surprisingly little variation in the shape of the DRLs between primates; DRL shapes were qualitatively similar but more variable in mammalian DRLs, consistent with the longer divergence time. This comparison is shown in **Fig. 1b** for mono-A/mono-T repeats, which are the most prevalent in the human genome and are the primary focus of our subsequent analyses (normalized DRLs for additional motifs shown in **Fig. S3**). Consistency of the shape of the DRLs across the primate lineage suggests that both the repeat tract length distributions and, as a corollary, maintenance of the underlying mechanisms, were largely stable for at least 70 million years. This highly conserved DRL evolution directly suggests the emergence of a steady state equilibrium.

**Fig. 1.**
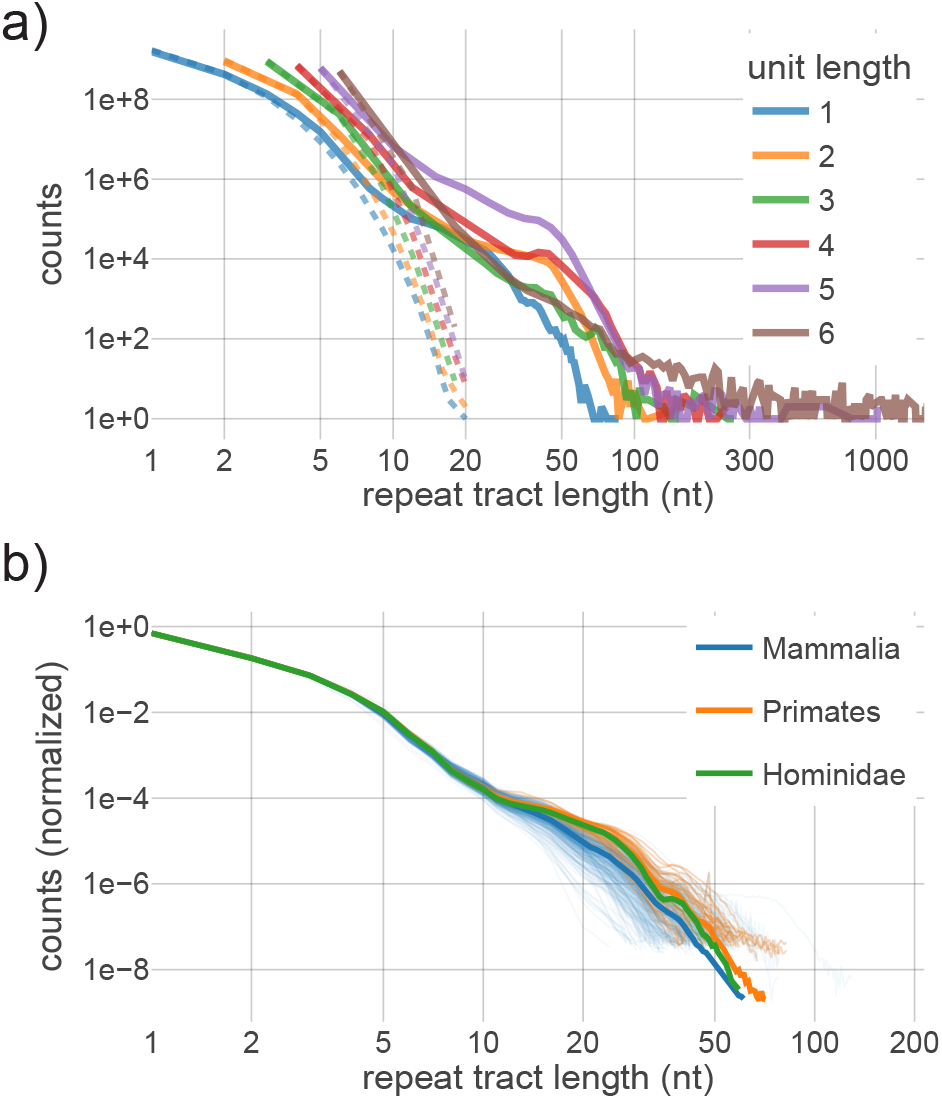
Repeat tract length distributions (DRLs) by motif length and across phylogenies. **a)** Counts of repeats in human T2T genome pooled by motif unit length (e.g., unit length 1 pools DRLs for A/T and C/G). Dashed lines represent counts in a randomly-shuffled human genome sequence. Canonical centromeric and telomeric motifs are excluded from unit lengths 5 and 6, respectively, due to qualitative differences in the DRLs. **b)** Normalized DRLs of mononucleotide-A repeats in mammals, primates and hominids. (See **Fig. S2** for other motifs.) Counts are necessarily normalized to account for different genome lengths (see **Methods**). Solid line indicates median values per length bin. Median calculations within phylogenies are inclusive (e.g., primates are included as a subset of mammals). Thin transparent lines show individual species. Similarity within phylogenies suggests long-term stability of the DRLs.

The empirical DRLs extend to lengths that, at disease loci, would be subject to genetic anticipation, risking progression to repeat expansion disorders in subsequent generations^5^; despite the associated disease risk, this tail of long repeats appears to be a generic and evolutionarily conserved feature of repeat length distributions. One proposed explanation is that longer repeats confer a selective advantage due to some repeat length-specific biological function^17^. As an alternative, we propose that long repeats emerge and are maintained by the complex interplay between distinct mutational forces. Though these hypotheses are not mutually exclusive, we sought to understand the extent to which mutagenesis, alone can maintain the shape of the distribution, without introducing natural selection.

### Mutational transitions in repeat tract length

As described above, literature suggests that repeat instability emerges as a very rapid increase in the rate of length changes as tract length increases, with the longest repeats mutating nearly every generation. In light of such high mutation rates, the observation that the DRL evolves in steady state over long timescales is somewhat surprising, suggesting that the maintenance of the distribution results from a dynamic balance between the ensemble of mutational processes that alter repeat length.

To better understand the genome-wide distribution, we therefore require a comprehensive understanding of all involved mutational processes (e.g., nucleotide substitutions, insertion, deletion; see **Fig. 2** for schematic of mutational processes) and how they differ by repeat length. Estimating repeat tract lengths from sequencing data is a notorious bioinformatic challenge, particularly for homopolymer repeats ^35,36,37,38^. Published results only sparsely cover the full range of lengths observed in the genome, largely focusing on disease relevant lengths and loci^40,41,42,43,44,45,46,^. In contrast, there is little information about mutation rates at short tract lengths despite comprising the vast majority of repeats in the genome.

**Fig. 2.**
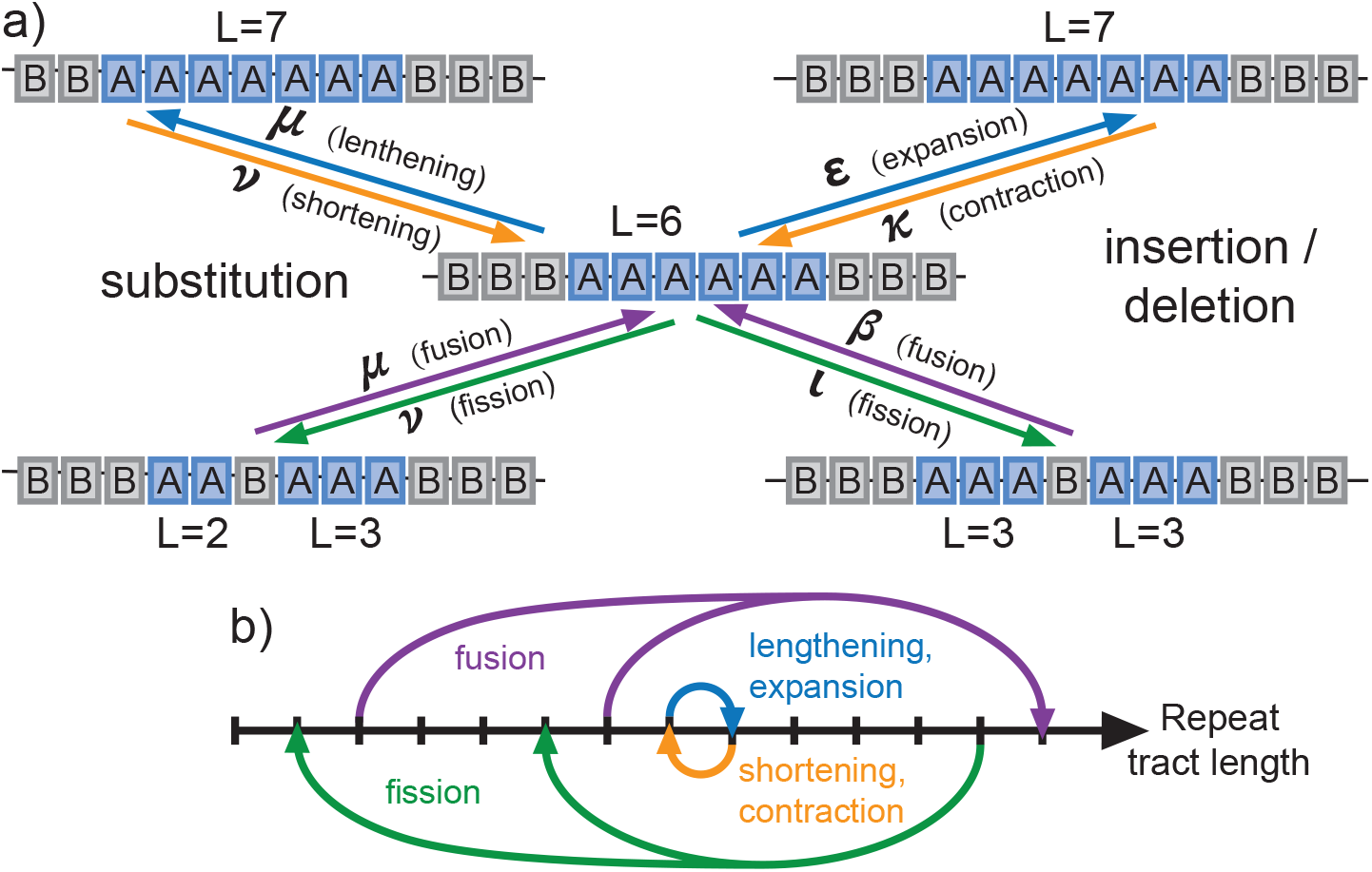
Mutational transitions between repeat tract lengths. **a)** Mutation types, using the example of transitions to and from repeat tract length *L*=6. ‘A’ represents a given STR motif; ‘B’ represents any other sequence with length equal to A. Arrows indicate mutations, either substitutions (left) or indels (right), that affect the length of A repeat tract(s). Mutations can lengthen/shorten (top) or interrupt/rejoin (bottom) repeat tracts. The latter we term repeat ‘fission’ and ‘fusion,’ respectively. **b)** Depiction of the same mutational processes as length transitions in the DRL. Lengthening/shortening mutations increase or decrease length by one unit (‘local’ transitions) and maintain the same total count of repeats (‘conservative’), while fission/fusion processes are non-local and non-conservative.

In order to study mutations across a wide range of tract lengths, we first subdivided insertions and deletions into repeat-relevant mutational categories; we refer to expansions and contractions as mutations that alter repeat length by whole motif units and maintain one contiguous repeat tract, in contrast to partial deletions and non-motif insertions. Rates of each mutagenic process were estimated by pooling existing short-read trio sequencing datasets (n=9,387 trios; henceforth, ‘pooled trio’ dataset). This data was sufficient to directly estimate length-dependent rates for short repeats (up to roughly *L*=6-8 units, depending on motif, where *L* is the number of repeated units in a tract) but observed that sequencing errors dramatically reduced mutation counts for longer repeat tract lengths (see **Methods**). We complemented these estimates by length-stratifying data from a recent study^46^ that used a population structure-aware caller (named ‘popSTR’) to study repeat mutations in the mid-to-long length range in short-read trio data (n=6,084). Due to a variety of technical considerations (see **Methods**), estimates were only reliable within a limited range of tract lengths for each motif, which differed by dataset.

It was previously observed that the majority of mutations within a repeat increase or decrease length by one unit (i.e., *L→L*±1)^47,48^. To expand on this, we length-stratified the mutation data and found that single-unit length changes dominate above a clear length threshold, consistent with the onset of repeat instability (**Figs. 3a, S4**). Accordingly, we estimated the rates of single-unit length changes, separately estimating the contraction, expansion, and non-motif insertion rates from the pooled trio data; popSTR-based estimates combine expansion and non-motif insertion rates due to technical limitations (see **Methods**). For mono-A repeats, all instability rates increase rapidly between roughly 5-10 nt (**Fig. 3b**, see **Fig S5** for all motifs), consistent with a threshold-like onset of repeat instability in this length range (detailed below). The popSTR-based estimates suggest that repeat instability rates continue to increase monotonically, at least until the length range where the data becomes noisy (**Figs. 3b, S5**).

**Fig. 3.**
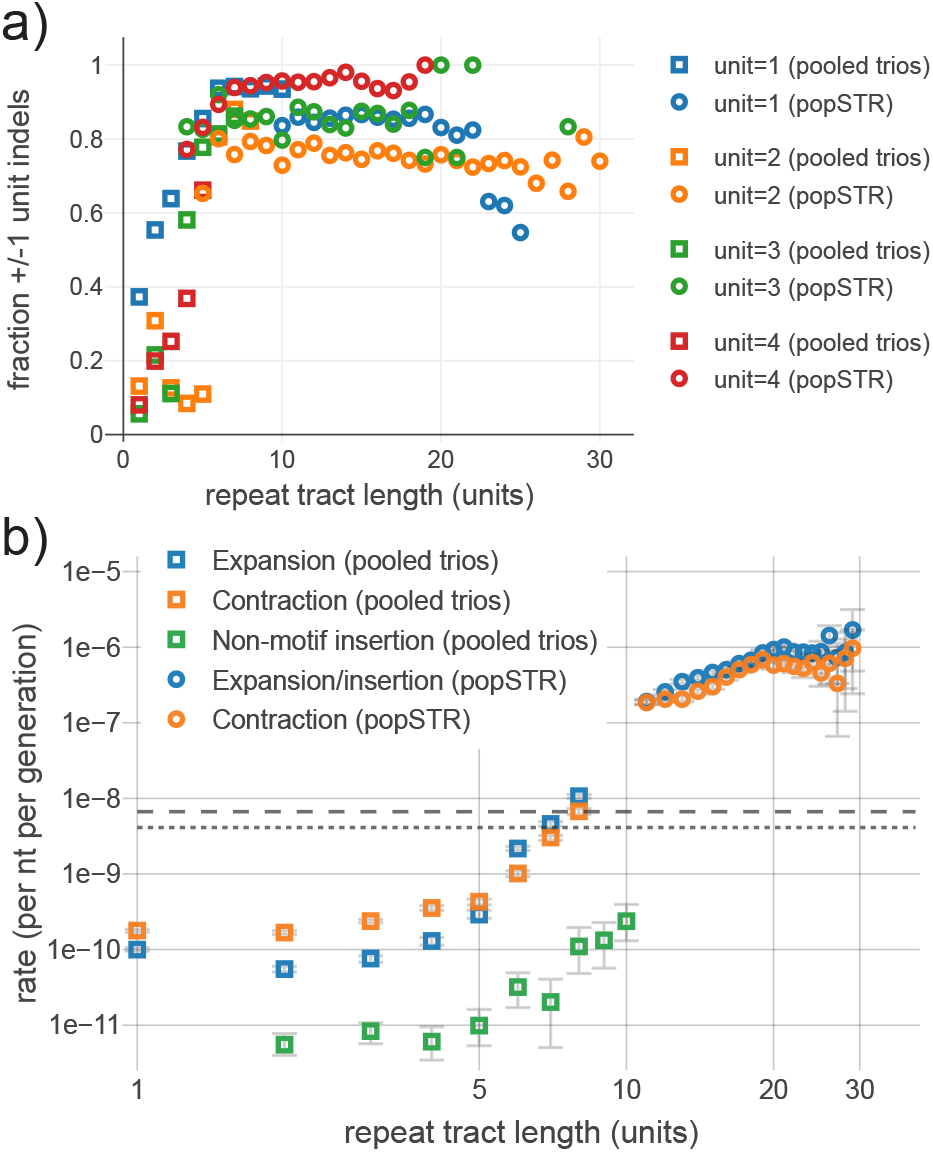
Estimated instability rates stratified by repeat tract length. **a)** Dependence of indel size on repeat tract length (x-axis) for repeat unit lengths 1–4 (different colors). Y-axis measures fraction of all indels that result in single unit length changes (i.e. number of inserted/deleted bases is equal to repeat unit length). Estimates shown for pooled trio (squares) and popSTR datasets (circles). Above a threshold of ∼5 units, repeat instability primarily consists of +/-1 unit changes. Tract lengths subject to severe technical artifacts were omitted for clarity. See **Fig. S3** for additional detail. **b)** Mononucleotide-A mutation rate estimates from pooled trio and popSTR datasets for expansions, contractions and non-motif insertions (note that popSTR dataset combines expansions and non-motif insertions). Rates calculated only from +/-1 unit changes. Statistical error bars show 95% CI assuming Poisson mutation counts. Gray dashed and dotted lines show average substitution rates ***μ*** (A>B, where B=C,G or T) and ***ν*** (B>A), respectively. Tract lengths subject to severe technical artifacts were omitted (see **Fig. S4** for complete estimates and additional motifs).

The combination of both datasets recapitulates the hallmark of repeat instability ^5,6,8,9^: a rapid increase in the rates of expansion and contraction as length increases. Beyond confirming this property, available data was insufficient to robustly estimate the length dependence of each mutational process across the tract length range observed in the genome. Such an estimate is a necessary component of any quantitative understanding of the approach of the DRL towards the steady state (as observed in primates). For further analyses, the length dependence of these mutation rates was extended to longer tract lengths by parameterization in an inference framework described below.

### Computational modeling of DRL dynamics

We sought to assess whether our three observations of the empirical human DRL, the existence of a long-term steady state, and the estimated repeat instability could be simultaneously incorporated into a self-consistent model of repeat length evolution. To this end, we built a computational model that incorporates length-changing effects of substitutions, expansions, contractions, and non-motif insertions to follow the evolution of the DRL towards a steady state. We modeled the distribution of mono-A repeats, in part, because they are subject to the simplest ensemble of mutational changes in length (related to the lack of distinction between tract length and the number of repeated units). As a consequence of counting of contiguous repeats to assemble the DRL, mutational processes alter the length of a given repeat in one of four ways (see **Fig. 2**): lengthening, shortening, joining two repeats into one (which we term ‘fusion’), or splitting one repeat into two (i.e., repeat interruption, which we term ‘fission’). This treatment of interruptions as effectively splitting a repeat is consistent with previous observations that interruptions result in locus-wide rates scaling with the longest contiguous subunit^49,50,51,52,53,54,55^ or, alternatively, a rate reduction that scales with distance from the repeat boundary^7^.

To reduce the computational time required to evolve a whole genome sequence and simultaneously count contiguous repeat tracts, we directly evolved the DRL by manipulating the occupancy of each length bin. In this formulation, length-altering mutations are reframed as transitions between length bins and the DRL evolves under repeated application of mutations over many generations. However, the elementary step of the process is deceptively complex, as repeat fusion precludes framing the mutational process as a standard transition rate matrix and both fission and fusion are non-conservative transitions. We treated the aggregate mutational effects as deterministic, ignoring stochasticity in the mutational process and due to factors like genetic drift, to approximate the expectation of the DRL at late times. We interpret this late-time expectation as an approximation to the steady-state distribution, if one exists, resulting from the modeled mutational processes.

### Bayesian inference and parametric model comparison

We constructed a Bayesian inference procedure (see **Fig. 4**) to constrain properties of repeat instability consistent with the steady-state evolution of the observed human DRL (see **Methods**). The computational model uses explicit length-dependent rates for each mutational process as inputs. We directly incorporated the subset of estimated rates from the pooled trio data shown in Fig. 3b (i.e., for *L*=1—8); the instability rate curves were then extended to longer lengths to model a rapid, monotonic increase with length. The inclusion of empirical estimates at low lengths, which include a rapid rate increase, limits the number of parameters required to describe complex length-dependent rates of repeat instability with both a rapid transition and a distinct asymptotic functional form for long repeats.

**Fig. 4.**
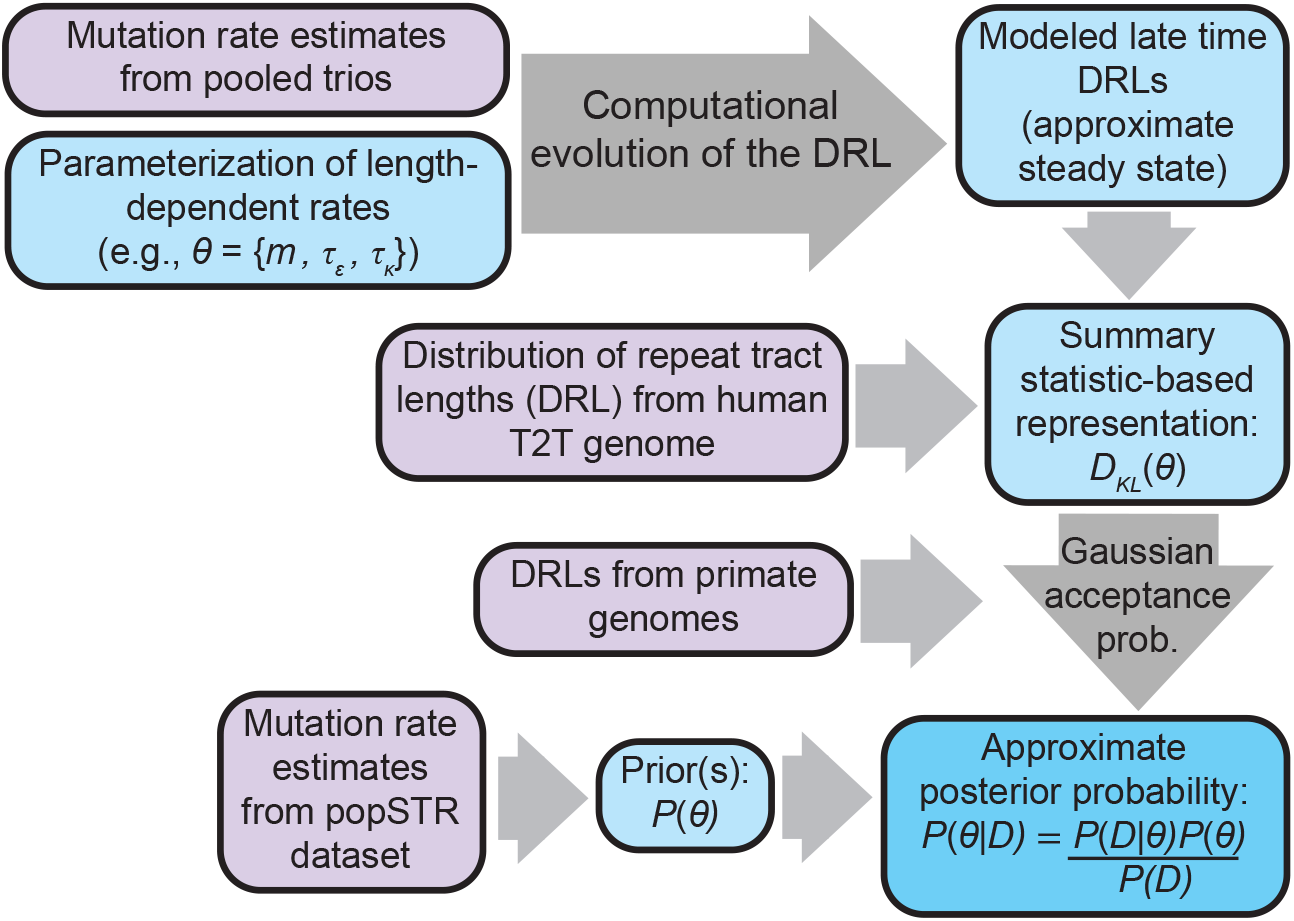
Schematic representation of Bayesian inference procedure. Inference of parameters representing length-dependent instability rates via Approximate Bayesian Computation (ABC). Empirical data sources informing the inference are shown in purple. First, expansion and contraction rates are parameterized at lengths where estimates are unreliable or unavailable. Each parameter combination specifies a complete set of mutation rates for repeats of all lengths. Given an initial state, mutational transitions (see **Fig. 2**) are repeatedly applied to evolve the DRL for a large number of generations. The late time DRL is treated as an approximation to the steady state DRL (when applicable). For each parameter combination, the difference between the late-time DRL and the human T2T DRL is summarized by the Kullback-Leibler (KL) divergence. The approximate posterior is proportional to the product of the prior and the probability of acceptance under an ABC rejection strategy [Wilkinson]. The prior is informed by popSTR-estimated mutation rates. The acceptance probability is treated as Gaussian-distributed in the KL divergence, with mean zero and variance defined by divergences for an ensemble of primates (see **Methods**). This quantity is calculated for each parameter combination and subsequently normalized over parameter space to approximate the posterior probability distribution.

Resemblance to the monotonic increase seen in popSTR estimates at intermediate lengths (**Fig 3b**) motivated a class of parameterizations with power-law increase in the rates (see **Table 1, Methods**). We first specified a model with minimal degrees of freedom that describes equal rates of expansion and contraction (i.e., rates for *L*>8 represent a simplistic model of replication slippage based on previous literature^8,20,23,28^). This was treated as a null model for comparison to parameterizations with additional degrees of freedom that characterize expansion-contraction bias. We used the popSTR-based rate estimates to define plausible, empirically based Bayesian priors for each parameter space, representing varying degrees of confidence in this dataset (**Fig. S6**), including a (naively uninformed) uniform prior. We then used the results of our computationally modeled DRLs in an Approximate Bayesian Computation framework (following the prescription in Wilkinson, 2013^56^) to compute a posterior probability distribution for each parametric model. We used the range of primate DRLs to define a rejection probability by comparing them to the human DRL (using Kullback-Leibler divergence, which quantifies the difference between two distributions). The same quantity was computed for each computationally modeled DRL and used to approximate the posterior probability distribution over the parameter space (see **Methods** for details).

**Table 1:**
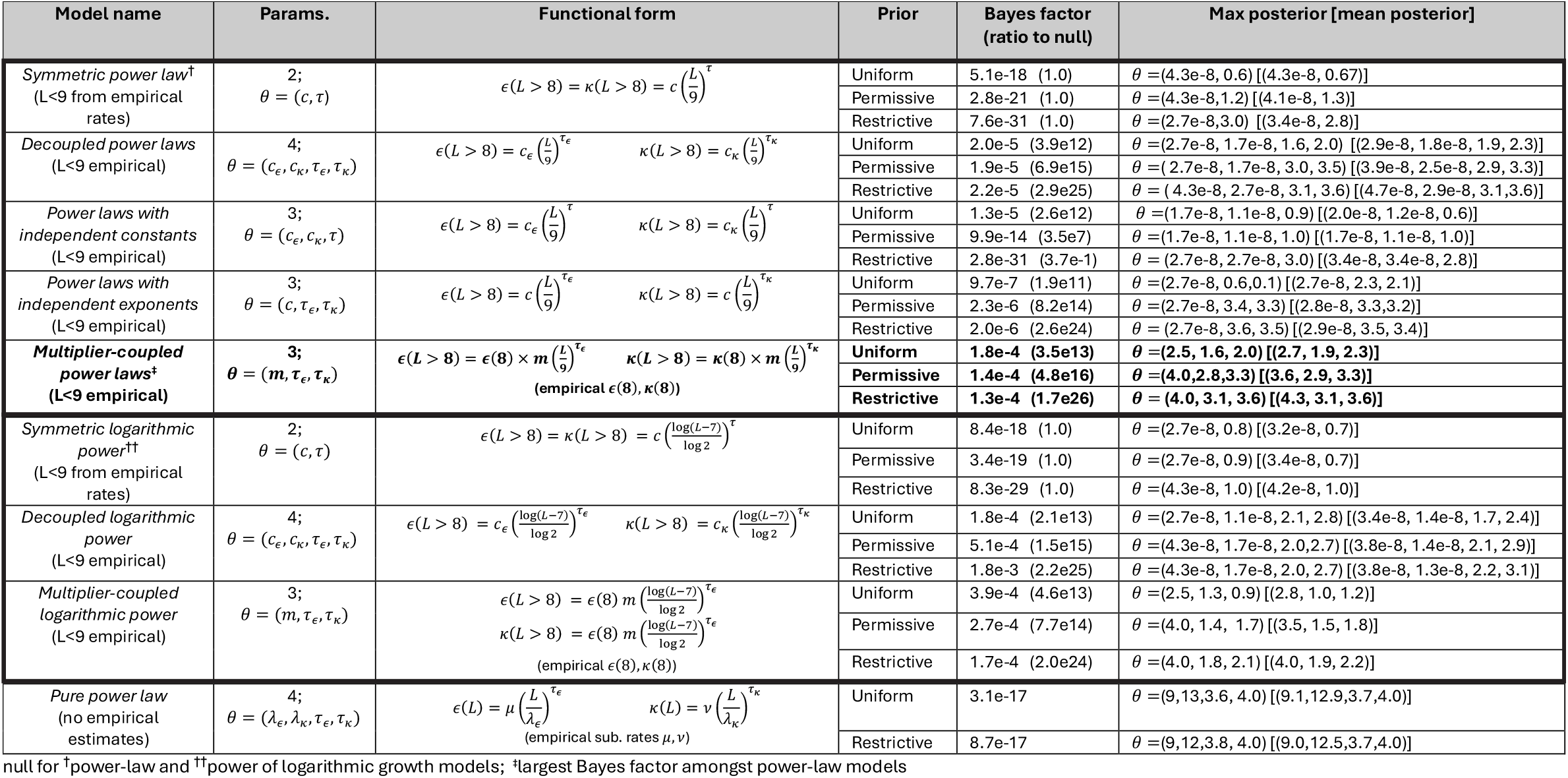
Parametric models of instability rates and summary of Bayesian inference results. For each parameterization used in our analyses, this table specifies the model name (as referred to in the text), the tract lengths described by the parameterization, the inference parameters, the functional forms for length-dependent expansion and contraction rates, and a summary of inference results. For each model, the following quantities are given for each prior: Bayes factor (and Bayes factor ratio to null model within the same nesting, denoted by symbols), parameter combination with maximum posterior probability, and mean posterior parameter combination. Distinct nestings separated by bold borders. Primary model considered is shown in bold text. Further detail on prior construction, calculation of Bayes factors (and model comparison), and expectation used to compute mean posterior parameters provided in **Methods**.

The results of our inference for each parameterization are summarized in **Table 1**. We assessed the relative statistical support for various model comparisons via the Bayes factor ratio (see **Methods**). The relative Bayes factors strongly suggested discarding the two-parameter null model in favor of further degrees of freedom that introduce asymmetry (i.e., bias) between the expansion and contraction rates. The power-law parameterization with the largest Bayes factor, regardless of prior, was a three dimensional model of expansion and contraction with distinct exponents and related multiplicative constants (see **Table 1, Methods**). This model is a modest improvement over the four-parameter description (i.e., completely decoupled expansion and contraction rates), which otherwise provides a dramatically better description than lower-dimensional parameterizations.

To test how reliant these conclusions are on the power-law functional form, we defined an alternate class of parameterizations with logarithm-based growth rates (i.e., with slower-growing rates at large lengths to better approximate saturation). Model comparison within this class of parameterizations provided qualitatively consistent results to those comparing power law parameterizations (see **Table 1**). The inequivalence of priors across functional forms (along with additional necessary approximations; see **Methods**) suggests caution should be taken in direct comparisons between models with distinct functional forms. Due to the relative analytic simplicity of the functional form, subsequent analyses were focused on power-law parameterization results.

### Inference of instability rates from the steady-state repeat length distribution

Amongst power-law models, we focused on the three-parameter multiplier-coupled model, as it showed the strongest statistical support, regardless of choice of prior. Above L=8, this model is parameterized by exponents, *τ*_*ϵ*_. and *τ*_*κ*_, and a common multiplier *m* (representing a discrete jump in rates immediately above the empirical estimates; explicit definition in **Methods**), which together characterize the length dependence of expansion *ϵ*(*L*; *m, τ*_*ϵ*_) and contraction *κ*(*L*; *m, τ*_*κ*_) (see **Methods**, Table 1 for full definitions). The common multiplier for expansion and contraction, which limits the dimensionality, may be interpreted as representing the onset of repeat instability due to some common biological mechanism. Our trio rate estimates rapidly rise in the length range where they lose accuracy; *m*, which describes a potentially dramatic jump immediately above this range, can provide an oversimplified characterization of a rapid transition to power-law like behavior. To limit further degrees of freedom, we assumed that the length dependence of non-motif insertions is dictated by *τ*_*ϵ*_, the expansion rate exponent, due to their parallel increase in de novo rates (**Fig. 3b**) and because they likely arise from the same biological mechanism (e.g., synthesis of the inserted nucleotides by an error-prone polymerase). The parameter space we explored includes the possibility of a constant per-nucleotide rate (i.e., *τ*=0, analogous to the constant per-nucleotide substitution rates), linearity (i.e., *τ*=1), a natural conceptual model for length dependence associated with repeat instability, and more rapid growth on par with popSTR-based estimates (**Fig. 3b**). However, the parameterization itself is not intended to represent a specific biological model; the true rate curves are likely more complex due to multiple contributing mechanisms.

To interpret the resulting posteriors, we first approximated highest density regions (HDRs) comprising 68%, 95%, and 99.7% of posterior probability on the finite grid for the multiplier-coupled model (**Figs. 5a, Fig. S7a**; alternate parameterizations shown in **Fig. S8, S9**). The posterior is largely localized along a ridge of constant values of Δ*τ* ≡ *τ*_*κ*_ − *τ*_*ϵ*_ (roughly, Δ*τ* ≈ 0.3 − 0.6) and roughly between multipliers *m* = 1 − 4, with larger average exponents for smaller multipliers (i.e., anticorrelation between *m* and *τ* values for the most probable parameters); the difference between expansion and contraction rate exponents appears to be more relevant than their specific values. Under the uniform prior, some parameter combinations within the 95% HDR deviate from this range of Δ*τ* (extending to both lower values of Δ*τ* and lower *τ*_*ϵ*_, *τ*_*κ*_; **Fig. 5a**) but are excluded when applying the popSTR-based prior, which requires consistency with the larger estimated instability rates (**Figs. 5a, S7a**).

**Fig. 5.**
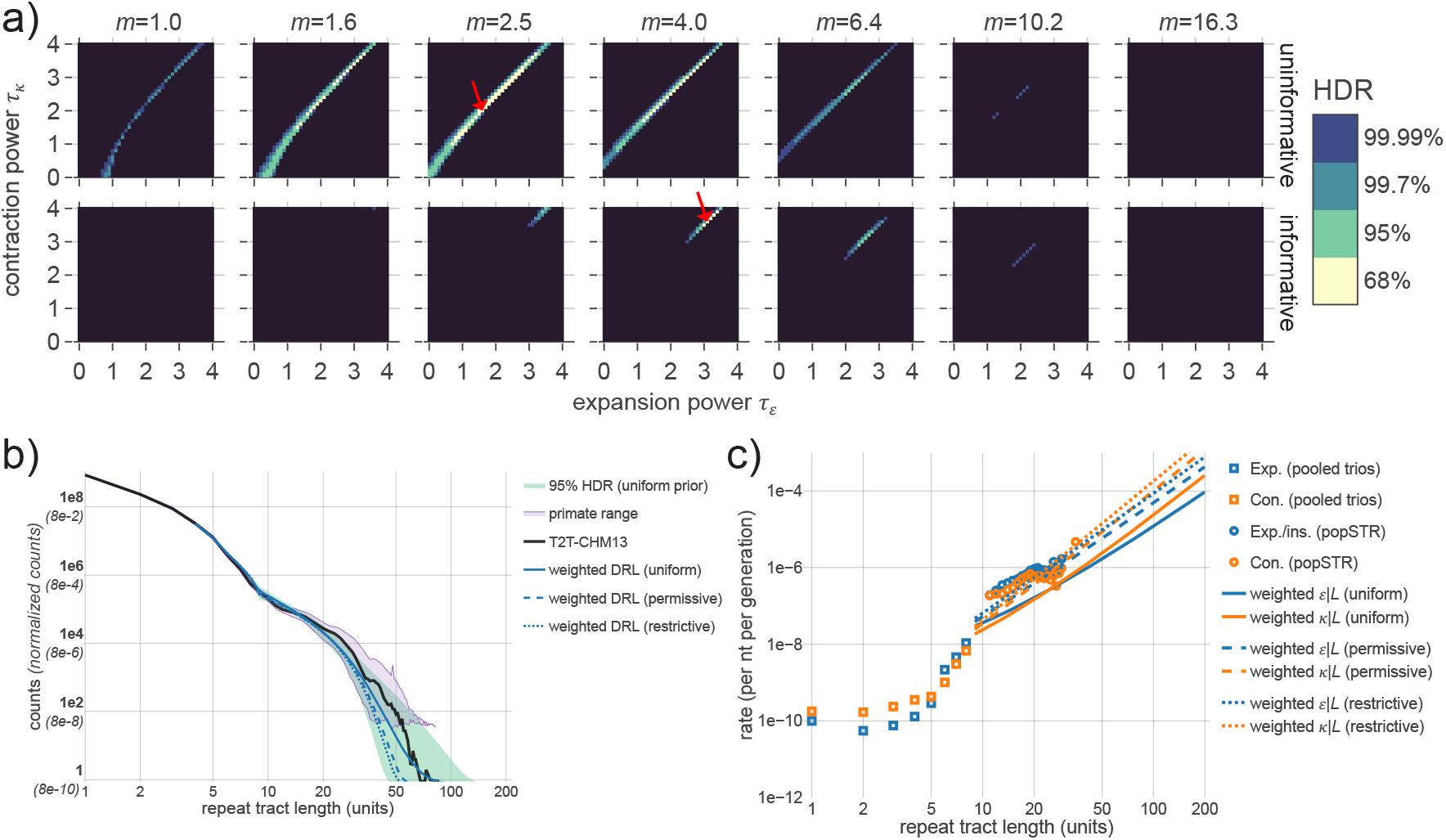
Inference results and self-consistency of a mutation-only model. **a)** Inference results for three-parameter multiplier-based power-law model of repeat instability rates. See **Fig. 4** for inference procedure. Plots display Bayesian posterior probabilities after applying uninformative prior (top row) and popSTR-based informative prior (bottom row; restrictive condition). Each coordinate represents a distinct set of length-dependent instability rates defined by parameters (*m*, τ_*ε*_, τ_*κ*_ ) shown in columns, x-axes and y-axes, respectively; τ_*ε*_ and τ_*κ*_ determine the power laws for expansion and contraction, respectively, and *m* is a multiplicative jump at L=9 (parameterization defined in **Table 1**). Color indicates highest density range (HDR) of the posterior for various total probabilities; black region sums to 0.01% of the probability. Red arrows show maximum posterior for each prior. Informative prior results in a more rapid increase in instability rates with length. See **Fig. S5** for construction of priors; see **Figs. S6-9** for posteriors under various parameterizations. **b)** Comparison of empirical DRLs for mono-A repeats to inference results. Counts for all DRLs are necessarily normalized for comparison (conditional on *L*>3, see **Methods)**; y-axis indicates normalized fractions (parentheses) and counts rescaled to match the number of repeats in the T2T genome (bold labels). Blue lines represent posterior-weighted DRLs (an average of all DRLs weighted by the posterior probability for each modeled parameter combination; see **Methods**) for informative and uninformative priors; modeled DRLs are largely consistent with the empirical T2T DRL (black). Range shown in green represents the minimum and maximum counts at each length bin across all parameters in the 95% HDR under the uniform prior. Purple region shows the min-max range generated from 36 primate genomes (after removing the two most-diverged DRLs and truncating each DRL where raw counts drop below 30; see **Methods**). The overlap between these two ranges indicates that the range of probable DRLs under the uniform prior largely reflects the ensemble of primates. **c)** Posterior-weighted repeat instability rates. Tract length dependencies of expansion (blue) and contraction rates (orange) shown for specified priors. For comparison, empirical estimates are shown for pooled trios (squares; directly incorporated in model) and popSTR data (circles; used to construct informative priors). Informative priors impose consistency with popSTR-estimated rates, while the posterior-weighted DRLs (panel b) remain consistent with the T2T genome.

We computed the posterior-weighted DRL (i.e., the expectation value of the DRL; see **Methods**) and the range of DRLs consistent with the 95% HDR (**Fig. 5b**). The posterior-weighted DRLs closely resemble the human genome-wide distribution, while the 95% HDR parameters roughly span the range of primate DRLs used in our inference procedure. This demonstrates that the coarse features of the empirical DRL can be recapitulated from mutational dynamics alone.

We then computed the posterior-weighted length-dependent rates of expansion and contraction for each prior and found rough consistency with popSTR-estimated rates (**Fig. 5c)**. One salient feature emerged, regardless of prior: expansion bias at intermediate tract lengths transitions to contraction bias at longer lengths due to the faster increase in contraction rate with length (i.e., *τ*_*κ*_ > *τ*_*ϵ*_; see **Figs. 5a,c**). This likely explains the preference for the multiplier-coupled model, which necessarily inherits a modest initial expansion bias directly from empirical rate estimates. However, if the apparent expansion bias at L=8 is simply a consequence of homopolymer sequencing errors, correcting the direction of this bias would lead to statistical rejection of the multiplier-coupled model in favor of the four-parameter model (which lacks the *a priori* imposition of initial expansion bias on the parametrized rates). Regardless, inference results under the four-parameter model recapitulate the importance of a transition from expansion to contraction bias (**Fig. S10**).

To gain intuition for the preference of expansion-to-contraction biased parameters in the multiplier-coupled model, we contrasted the DRLs to the parameter combinations outside of the 95% HDR. Excluding slowly evolving rates, the remaining parameter space broadly separates into three qualitative categories, largely characterized by Δ*τ*: Δ*τ* ≳0.6 yields DRLs that underestimate the long repeat tail (i.e., early truncation), while 0 < Δ*τ* ≲0.2 yields distributions that overestimate the long tail (**Fig. S11**). The roughly half of parameter space with Δ*τ* < 0 showed a clear reason for near-zero posterior probabilities: regardless of multiplier, these parameter combinations do not converge to steady state at late times and are subject to explosive growth in all length bins (**Figs. S11, S12a**).

In addition to Δ*τ* values outside of the high posterior ridge, larger values of *m* generally remained beyond the 95% HDR. This was likely due, in part, to the discrete jump between estimated and parameterized rates at L=9 that results in a discontinuous DRL, which can artificially inflate the Kullback-Leibler divergence. To investigate this, we repeated our inference after smoothing the instability rate length dependences via interpolation around this transition (**Fig. S8**). The results suggested that a (naively more realistic) interpolated length dependence results in less penalization for larger multipliers and posterior probabilities more robust to the choice of prior. Additionally, inference using interpolated rates under the restrictive informative prior results in posterior-weighted instability rate estimates that overlap the popSTR estimates (**Fig. S8c**); this suggests a smoother length dependence may represent more realistic instability rates. Indeed, this mutational model incorporates all available data and describes a self-consistent picture of a steady-state DRL shaped only by mutational dynamics.

To better understand the approach to steady state for realistic parameters within the 95% HDR, we followed the temporal evolution of the DRL, starting from a highly diverged initial state (see **Methods, Fig. S12b**). This analysis suggested a two-stage equilibration process with two distinct timescales. The bulk of the long repeat tail establishes exponentially quickly, followed by a slower fine-scale equilibration of mutational processes at each length. Finally, we tested for robustness to potential confounders (e.g., differing initial conditions, use of a step-wise speed-up factor, lack of stochastic fluctuations, etc.) and found no major changes in the qualitative results (see **Methods, Fig. S13**). Collectively, these results show that mutational dynamics, rather than natural selection, may be responsible for maintenance of an excess of mid-to-long tract length repeats in the human genome.

### Maintenance of the repeat length distribution in steady state

To understand the complex interplay between mutational processes that shapes and stabilizes the distribution of repeat lengths, we constructed an analytic model of the dynamics. This analytic approximation captures the behavior of *DRL* after the mutational process reaches steady state (see **Methods** and **Supplementary Note SN1**), focusing primarily on the previously described multiplier-coupled three-dimensional parameterization. A number of previous studies have constructed mathematical models of repeat instability to study repeat length evolution^20,23,26,26,27,28,29,30,32,34,57^, including a notable study by Lai and Sun^31^ that incorporates many of the elements detailed herein. However, the combination of empirical rate estimates, a robust genome assembly, and our phylogenetic observations motivated the construction of a model from first principles that is directly informed by this collection of observations. In addition to differences in mathematical machinery, the analytic construction differs from previous efforts by incorporating pervasive length-dependent expansion-contraction bias (**Figs. 3b, S5**) and explicit effects from non-motif insertions.

We first constructed a discrete equation for the change in the number of repeats at a given length in a single generation due to the deterministic action of mutations (i.e., in the absence of selection and stochasticity in the mutational process, consistent with our computational model). We then imposed a steady state condition by requiring that the sum of all changes in and out of each length class vanishes at each time step after equilibration. Despite the simplifying assumption of steady state, the full dynamical equation cannot be solved generically. However, our estimates of de novo mutation rates suggested a dichotomy exists in the primary driver of changes in length between short and longer repeats (i.e., primarily substitutions for L<8 A-mononucleotide repeats vs. expansions and contractions for L>10; see below for direct inference of this length range). Accordingly, short and long repeat dynamics can be treated as separable (i.e., under the approximation of a separation of repeat length scales), leading to simpler approximations of both length regimes. Transitions between the short and long repeat regimes, while present, remain negligible in all realistic scenarios (see **SN1**).

For short repeats, we treated indel mutations as negligible and showed that a geometric distribution (see **Methods, Equation 8**) exactly solves the steady state equation under two-way substitutions alone (see **SN1**). For longer repeats, we constructed a partial differential equation (PDE) that approximates the discrete equation and studied its time-independent properties in steady state; dynamical equations are derived in **Supplementary Note SN1** in terms of generic parameterization of the length-dependent instability rates. Focusing on the multiplier-coupled parameterization, we obtained numerical solutions to the steady-state dynamical equations under various approximations and under the assumption that fusion-based contributions are negligible to long repeats (**Methods, Equations 9**—**11; SN1**). These solutions, along with the geometric distribution for short repeats, accurately describe the late-time DRLs produced by our computational model across the range of parameters that approach a steady state (**Fig. 6**; see **SN1** for additional comparisons). Using these comparisons, we found that, within some parameter regimes, the dynamics simplify to a less complex balance of mutational processes (**Methods, Equations 10**—**11; SN1**) and assessed the appropriate regime of validity (**Figs. 6a,b; SN1**). To more directly test the accuracy of the PDE, we used our computational results to decompose the per-generation fluxes in and out of each length class into relative contributions from each mutational type. This allowed for identification of the dominant mutational processes maintaining steady state (**Fig. 6c; SN1**); the accuracy of each approximation was confirmed by analyzing the net magnitudes of fission and fusion within each length class and regime (**Supplementary Note Figs. SN1, SN4**—**6**).

**Fig. 6.**
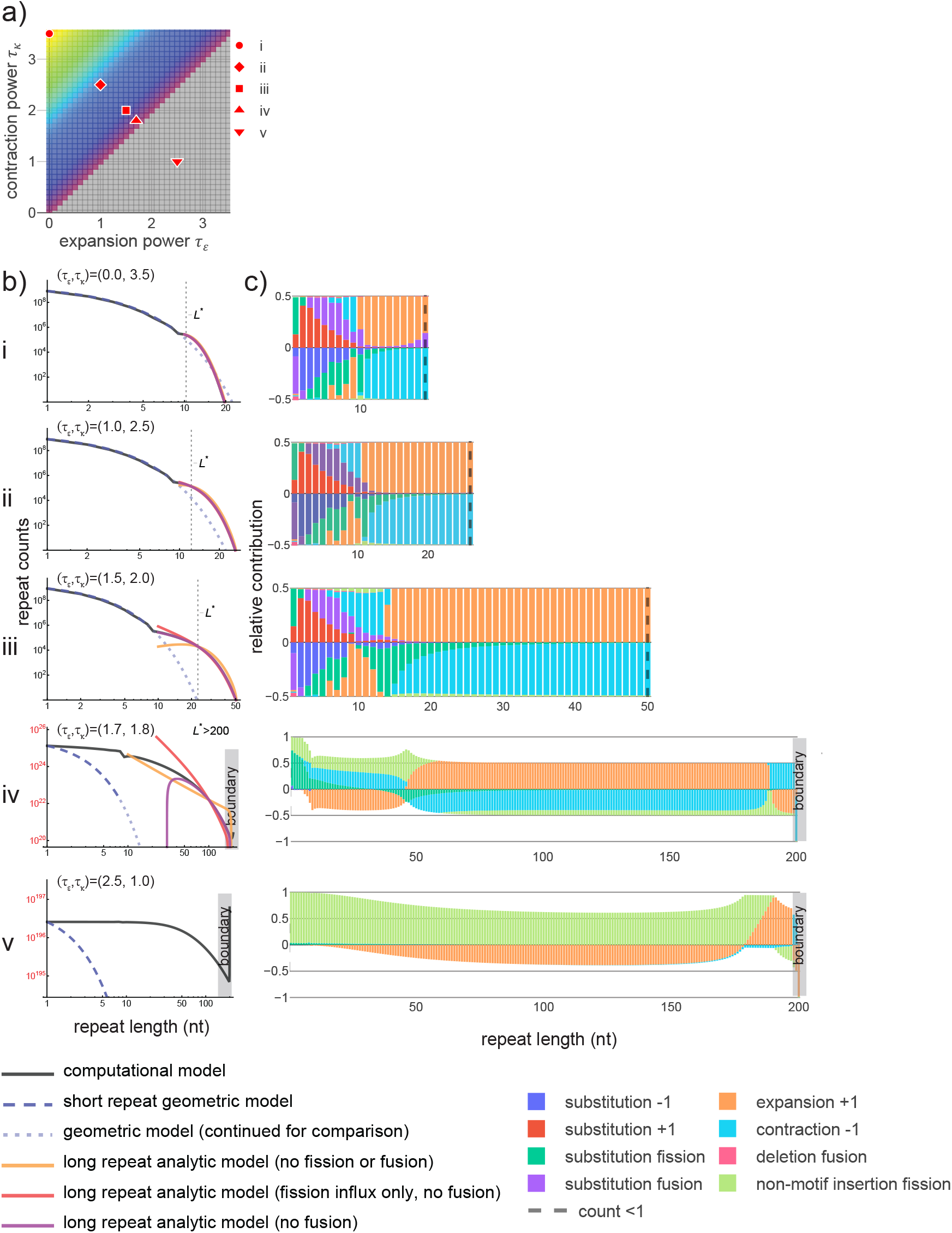
Dynamical regimes distinguishable by dominant mutational effects. **a)** Slice of parameter space for the three-parameter multiplier-based model (parameterization in **Table 1**). Five example parameter combinations with *m*=4 and τ_*ε*_ +τ_*κ*_ =3.5 are shown, corresponding to plots in (b) and (c). Colors roughly divide the parameter space into dynamical regimes. **b)** Comparisons between computational model results and numerical solutions of approximate steady state equations (see **Methods, Supplementary Note SN1**). Short length regime at equilibrium is geometrically distributed (blue dashed lines). For long repeats, numerical solutions are shown for three nested approximations to the steady state equation in the continuum limit (*L*≫1) in the absence of fusion (due to negligible rates). Local transitions (*L* to *L*+/-1) were modeled as a combination of symmetric (diffusive) and asymmetric (directional bias) components. L* represents length at which expansion and contraction rates are equal. For strong asymptotic contraction-bias (b:i), all three approximations remain valid indicating that the dynamics are well approximated by neglecting fission entirely (orange curve aligns). For moderate contraction bias (b:ii), outflux due to fission becomes non-negligible (orange begins to deviate). For realistic parameter combinations with lower contraction bias (b:iii), outflux due to fission is required at all lengths (orange fails); influx due to fission is required at intermediate lengths (red deviates); fusion remains negligible (purple remains accurate). Plots (b:iv-v) display non-equilibrium dynamics leading to rapid increase in repeat counts and explosive growth in genome size. Under universal expansion bias (b:v), DRL extends indefinitely above the boundary imposed at *L*=200 (for computational feasibility). Steady-state analytics do not apply. **c)** Relative contributions from each mutational transition to net flux (in minus out) per length bin, produced by computational model. Dashed line indicates DRL truncation (counts <1). Consistent with analytic predictions, fission is subdominant under strong contraction bias, has relevant outflux under moderate to weak contraction bias, and relevant influx at intermediate lengths under weak contraction bias. Equilibrium distribution is stabilized in detailed balance (net influx = outflux). Influx > outflux (c:iv-v) leads to nonequilibrium dynamics and indefinite genome growth.

We used this model to study the shape and stability of the empirical DRL and distinctions between repeats in different length regimes under mutational forces alone. Expansions and contractions remain non-negligible for any long repeat across the space of parameters that lead to stable late-time DRLs, highlighting the importance of repeat length instability to the maintenance of long repeats. For extreme parameters that stabilize (i.e., *τ*_*κ*_≫*τ*_*ϵ*_), the dynamics of all long repeats are dominated by expansion and contraction, alone, leading to a DRL that truncates more rapidly than under substitutions alone (i.e., a *depletion* of long repeats relative to a geometric distribution). In contrast, for realistic parameters (i.e., within the 95% HDR for A-mononucleotide repeats), an intermediate length regime emerges, characterized by the relevance of repeat fission. An accurate description of the shape of the DRL requires fission to account for the loss of repeats from the extreme tail (i.e., the longest populated length bins) and gain of intermediate length repeats. The relative contributions of fission due to substitutions and non-motif insertions are parameter-dependent; within the rough neighborhood of the maximum posterior parameters (informative prior), substitution is the primary driver of fission up to lengths of ∼20 nt, while longer repeats are primarily interrupted by non-motif insertions (see **SN1**). Fission-based losses in the extreme tail are insufficient to fully counteract length increases due to expansion, independent of the mutational mechanism and parameter values. Instead, contraction is primarily responsible for truncating the DRL at finite repeat length but can be bolstered by both substitution- and non-motif insertion-based fission. The dynamics of the long repeat regime decouples from that of short repeats such that rapidly mutating long repeats effectively become independently evolving genomic elements, categorically distinct from random sequences of the same length. The abundance of long repeats in the genome may therefore be a consequence of their largely unencumbered evolution caused by rapid changes in length.

### Inferring the onset of instability from the shape of the DRL

We sought to better characterize the length at which repeats become independently evolving genomic elements. Our analyses thus far suggests that this occurs roughly at the length where expansion and contraction rates exceed substitution rates. This length was explicitly fixed in our inference via reliance on empirical rate estimates at L=1—8; however, this precluded exploration of the onset length of repeat instability. To study the encoding of this information within the shape of the DRL, we defined fully parameterized rate curves (omitting all empirical rate estimates) that include the length at which instability rates exceed substitution as an explicit parameter. Expansion and contraction rates are each parameterized by an independent power law at all lengths (i.e., with no reference to empirical estimates) in terms of an exponent, *τ*, and, *λ*, the length at which the rate intersects the relevant substitution rate (*μ* for expansion, *ν* for contraction; full parameterizations defined in **Table 1, Methods**). This four-parameter model depicts an oversimplified rate dependence but serves as a toy model to probe the instability onset length *λ*.

Applying the same Bayesian inference pipeline, we estimated the posterior probability using both a uniform prior (i.e., excluding all rate data) and a prior informed by the combined set of empirical rate estimates (i.e., from both the pooled trio and popSTR data; see **Methods**). We marginalized the posterior to specify values of the onset lengths for expansion (*λ*_*ϵ ϵ*_) and contraction (*λ*_*κ*_) and found highly restrictive marginal distributions with 95% HDRs isolated to *λ*_*ϵ*_ = 9, *λ*_*κ*_ = 12 − 13 (informative prior: *λ*_*ϵ*_, *λ*_*κ*_ = 9, 12; **Fig. S14a, Table 1**). This recapitulates the range of lengths observed in direct empirical rate estimates, despite excluding all such data from the inference (i.e., isolating the influence of the shape of the DRL). The posterior-weighted DRLs reproduce the informative features of the human DRL (e.g., deviation from the substitution-driven geometric distribution at roughly 10 nt; **Fig. S14b**), despite the oversimplified model of instability. This suggests that the transition in shape of the DRL corresponds to the onset length of repeat instability, allowing for rough estimation of this key feature from visual inspection of the distribution.

### Application to repeats with longer unit length

Given that the DRL is informative about the onset length of repeat instability, we next compared this quantity across motifs of differing unit lengths (e.g. dinucleotides, trinucleotides). Empirical rate estimates for all motifs showed qualitatively similar properties to mono-A repeats (i.e., predominantly single-unit expansions and contractions with rates that scale rapidly with tract length; **Figs. 3a, S4, S5**), suggesting an analogous dynamical competition between substitutions and repeat instability that shapes the steady state DRL. We compared two distinct measures of the onset of repeat instability, representing long- and short-timescale information: first, rough lengths at which empirical DRLs first deviate from geometric decay (i.e., the expected DRL under substitutions alone) and second, the approximate lengths at which per-repeat expansion and/or contraction rates first exceed either substitution rate (i.e. rates comparable to *μ* or *ν* that perturb the geometric dependence on *L*). Both measures showed reasonable agreement confined to a range of onset lengths between roughly 6 and 12 nt, despite differing unit lengths (**Fig. S15a**). This suggests a universal description of repeat dynamics that shapes the extended tail of the DRL, despite apparent differences in the geometric portion at short lengths. Rapid geometric falloff is an immediate consequence of increasing the unit length (**Fig. S15b**): while a single substitution is sufficient to shorten tract lengths regardless of motif, lengthening of repeat tracts can require multiple substitutions (up to the unit length of the motif). Given initially comparable expansion and contraction rates across motifs, longer motifs show a more immediate transition to the repeat instability-dominated dynamics when measured in the number of repeated units but largely agree when measuring the onset length of repeat instability in nucleotides (**Fig. 1a**).

## Discussion

Motivated to understand the origin, prevalence and maintenance of simple tandem repeats in the genome, we constructed a model of repeat evolution under mutagenesis alone that bridges short- and long-timescale observations of repeat length instability. We demonstrate that mutations alone are sufficient to explain the shape of the genome-wide distribution of tract lengths. The abundance of long repeats in the genome reflects the rapid onset of repeat instability with an initial expansion bias, rather than natural selection. This observation does not preclude selection at specific loci, whether beneficial or disease-associated, provided these comprise a small portion of repeats in the genome.

Length-dependent expansion-contraction bias is evident in our de novo estimates; incorporating this property into the mutational model is sufficient to truncate the distribution at finite lengths due to substantial contraction-bias. The long length tail of the distribution is produced and maintained in a dynamic balance between expansion, contraction and fission. This implicitly prevents the growth of repeats to disease-relevant lengths, suggesting natural selection as a disease-prevention mechanism may not be essential. If selection, rather than contraction bias, is responsible for terminating the distribution below disease length, it would have to be enormously efficient to counteract instability-driven expansion rates and act globally across all sufficiently long repeats. If pervasive selection plays a role in shaping the distribution, this must be inferred as a deviation from the DRL under mutation alone built on a more complete model of repeat mutagenesis.

Our analysis of the genome-wide properties of repeats is complementary to studies of individual loci harboring disease, which generally occurs at or above the longest lengths present in the reference genome (i.e., stochastically driven length classes in the present context). Such elongated repeats can form motif-specific secondary structures that can disrupt replication and repair, causing instability with qualitatively distinct properties^8,9^. Furthermore, even amongst repeats of the same motif, locus-specific properties can introduce variability in the length-dependent rates of expansion and contraction and directional differences in bias (e.g., for long CAG repeat loci^7^). One well-studied example is the CAG repeat locus responsible for Huntington’s disease. A recent analysis showed that a secondary phase of expansion-biased, accelerated instability rates best explains somatic repeat expansion and its association with disease progression^57^. This locus-specific inference does not conflict with our observation that contraction bias terminates the bulk of the genome-wide distribution; indeed, this may indicate that, at lengths well-above those studied in the present manuscript, additional directional flips in bias may occur. This, along with potential inter-locus variability, may contribute to the modest number of repeats at lengths above the truncation point.

Our analysis offers a potential explanation for the prevalence of repeats at lengths that risk progression to disease. First, the dynamics of short and long repeats decouple due to the rapid onset, and subsequent dominance, of repeat length instability. Short repeat dynamics are dictated by substitutions alone such that repeats within this regime are roughly indistinguishable from random strings of nucleotides of the same length. Longer repeats are primarily subject to distinct mutational forces, exhibiting rapid expansions and contractions and a higher rate of repeat fission, which increases the total number of repeat tracts. Amongst long repeats, those of mid-length primarily experience substitution-based fission, while mutations in the longest repeats are effectively substitution-independent (i.e., fission is driven by non-motif insertion). This is inconsistent with previous literature that suggested substitutions prevent disease by providing a stopping force that counteracts indefinite expansion^45,49,55,58,59,60^; instead, our analyses suggest this is primarily a consequence of contraction bias at long tract lengths (similar to previous proposals based on very early data^25,27,33,47^.) Given the negligible role of substitution, there is little overlap in the mutational forces—and, subsequently, the underlying mechanisms—between the shortest and longest repetitive sequences included in our analyses. In this sense, long repeats emerge as independently evolving genomic elements (with parallels to the concept of selfish genetic elements^61,62,63^). Monotonically increasing instability rates generates length-dependent dynamics under which expansions lead to further instability, while decreasing length is effectively stabilizing; the former results in frequent forays into long length bins that may be the precursors to disease. The onset of this process leads to a natural definition for the shortest ‘unstable’ repeat (roughly 6—12 nt, far below disease length). This dynamical definition is distinct from measuring the lowest length where expansion or contraction rates start exceeding the background indel rate (as low as two units for many motifs; **Figs. 3b, S5**), which may better inform the molecular underpinnings of repeat instability. This difference in scientific goals underlies the debate in the literature concerning the definition of unstable repeats^64^.

Provided selection plays little role in directly modifying repeat length, the conservation of the distribution in steady state implies that the underlying mutational mechanisms (i.e., DNA replication and repair) are highly conserved. Generically, such mechanisms play a broad role in maintaining sequence fidelity of the entire genome, primarily preventing single nucleotide mutations; due to the substantially larger target size, it is unlikely that machinery responsible for both single site mutations and instability-driven length changes are optimized to properties of the latter. The abundance of long repeats may thus be an inescapable consequence of the pleiotropic function of the machinery maintaining genome-wide sequence fidelity.

It remains unclear which biological mechanisms control the key properties of repeat length instability described in our study. The proposed mechanism(s) should be able to explain length dependencies of instability rates (**Figs. 3b, S5**) that show: a) rapid onset from ∼6—12 nt, surpassing the rate of substitutions, b) greater-than-linear increase in the expansion/contraction rate per target above ∼10 nt, c) generically asymmetric rates of expansion and contraction with initial expansion-bias, followed by terminal contraction-bias, and d) single-unit expansions/contractions, regardless of tract length (**Figs. 3a, S4**). Surprisingly, these observations appear to be largely independent of both motif sequence and unit length (**Fig. S5**), suggesting a common biological origin.

Two widely studied mechanisms, replication slippage and mismatch repair (MMR), likely explains part of the story^8,9,65,66,67,68,69^. Slippage, when newly synthesized DNA partially unwinds and realigns out of register, should strongly depend on the unit length; however, we see only minor variation associated with unit length (**Fig. S5**). While slippage during DNA replication produces loop-outs on both strands symmetrically^68^, subsequent small loop-processing by MMR preferentially results in contractions^70^ due to bias towards the nascent strand^71^. Slipped-strand structures may be a motif-independent source of loop-outs subject to the same MMR-processing; in contrast, other secondary structures are motif-specific and therefore cannot be the primary source of repeat instability but can potentially explain differences between motifs^72^ (**Fig. S1b**). Importantly, the observation of mostly single-unit expansions/contractions argues against mechanisms involving larger structures (e.g., long hairpins that cannot be processed by MutS*β*_73,74,75,76,77,78_), as these would be expected to generate multi-unit indels.

Single-unit expansions have also been observed in a different context: Okazaki fragment maturation by flap-endonuclease FEN1^79^. Imprecise removal of the flap formed by the displaced 5’-flank of an Okazaki fragment may lead to expansion bias^80^ and introduce an associated length scale. A secondary mechanism takes over when flaps exceed 30 nt^81^; speculatively, long repeats could give rise to long flaps. Likewise, another flap-endonuclease, FAN1, which recently emerged as a genetic modifier of several repeat expansion disorders, was implicated in processing of various slip-outs and demonstrated differential activity depending on flap length^82^. Altogether, this illustrates how different mechanistic explanations may apply to repeats of distinct lengths, generating emergent properties like length-dependent expansion-contraction bias.

In addition to advancing a mechanistic understanding, substantial effort continues to be dedicated to both assembling datasets and developing estimation techniques specific to repeat instability, due to the inherent difficulties associated with repetitive DNA. Given the difficulty of this task, the present work demonstrates how direct rate estimates can be informed by orthogonal data. The comparative robustness of estimates of the distribution of repeat lengths provides constraints on properties of instability that can serve as a new means for evaluating the quality of differing rate estimates. The DRL may also serve as a summary statistic informative about the evolutionary history of mutation rates and mechanisms, including in species where no population data exists. Indeed, our rapidly improving understanding of repetitive elements, which have historically evaded sequencing efforts, unlocks a range of new questions about the composition and evolution of the genome.

## Methods

### Genome sources

Genome fasta files for T2T-CHM13_2.0 were downloaded from UCSC: http://hgdownload.soe.ucsc.edu/downloads.html#human. Alternate human assemblies and mammalian genomes were downloaded from the NCBI genome database: https://www.ncbi.nlm.nih.gov/datasets/genome/

### Motif labeling

Throughout the present study, repeat motifs are given a standardized label according to alphabetical order within the list of all cyclical permutations of a given motif (e.g., CAG, AGC, GCA) and their reverse complements (e.g., CTG, GCT, TGC). Outside of coding regions, cyclical permutations of a motif become mostly indistinguishable, both bioinformatically and biologically (after exceeding some minimal length relevant to processes such as protein binding site recognition). Likewise, if not considering specific hypotheses such as transcription direction, reverse complementary motifs should be treated as equivalent because Watson and Crick strands are assigned to each chromosome arbitrarily. The present study does not investigate any of these specific biological hypotheses, and so we combine results for all equivalent motifs under a single label to increase statistical power. In this arrangement, well-studied motifs may receive a label that differs from that commonly used in the literature (e.g. Huntington’s disease (CAG)_n_ repeats and myotonic dystrophy (CTG)_n_ repeats are both labeled ‘AGC’.)

### Generation of empirical repeat tract length distributions

Repeat tract length distributions were generated by counting consecutive complete motifs (i.e., perfect motifs, no interruptions and no partial motifs). Each distribution was assembled by counting contiguous, uninterrupted repetitions of a specified motif. Instead of introducing an arbitrary tolerance for interruptions when counting repeats of a given length, this strict definition allows for straightforward bioinformatic assembly of the DRL. The *regex* pattern ‘([ATGC]{1,6}?)\1+’ detects arbitrarily long tracts of repeated nucleotides, finding any motif with unit length of 1-6 nt. Using a *regex* implementation in Python 3 (https://pypi.org/project/regex/), all motifs can be detected simultaneously by using the ‘finditer’ command with the ‘overlapped=True’ option. Because this pattern detects repetitions of motifs, separate *regex* patterns were used to detect single instances of each motif (i.e. *L*=1), taking the form ‘([ATGC]{*n*})\1{0}(?!\1)’, where *n* is each motif unit length 1-6. The results of all regex searches were combined to generate a histogram of counts for each motif (pooled under the appropriate label) at all tract lengths present in the genome (i.e., the distribution of repeat tract lengths, DRL). Histograms representing counts of non-motifs (i.e., the lengths of contiguous regions where a particular motif is absent; required for computational modeling) were generated on a per-motif basis, using the *regex* pattern ‘(?:(?!’ + motif + ‘)[ATGC])+’ and combining the results for all cyclical and reverse-complementary permutations of the given motif.

Bootstrap confidence intervals were generated around the T2T-CHM13 repeat length distribution. The genome was divided into 1Mb contiguous non-overlapping segments, discarding any sub-1Mb chromosome ends. DRLs were measured for each segment. A distribution for the full-length genome was then reconstituted by randomly sampling from these segments, allowing replacements, and summing the distributions from each segment. This process was repeated 1000 times, after which 95% confidence intervals were generated by separately taking the minimum and maximum in each length bin by removing the top and bottom 25 counts.

For the various mammalian genomes, the same counting procedure was applied. Assemblies generated from short-read sequencing frequently contain many short contigs which typically originate from poorly-sequenced regions containing transposable elements; any contig of length <10 kb was discarded. Taxonomic data was retrieved from https://ftp.ncbi.nlm.nih.gov/pub/taxonomy/. The median distribution of a given taxonomic group was assembled by gathering the normalized DRLs (see below) for every member of the group (i.e., for primates this includes humans, and for mammals this includes primates), and taking the value of the median species for each length bin.

### Distribution normalization

After initially computing the DRL for each motif from the T2T genome, we sought to compare the shape of each histogram of raw counts to those assembled from distinct human reference assemblies (**Fig. S1c**) and from references for various species (**Figs. 1b, S2**). To compare distributions estimated from assemblies with differing total target size, it was necessary to normalize each distribution (i.e., divide by the total number of counts, summing over length bins) to standardize the overall scale. We refer to the normalized DRL as the probability distribution of repeat tract lengths, which we interpret as an estimate of the probability of randomly sampling a repeat of length L from the set of all contiguous motifs (including *L*=1) in the assembly; when specific length classes are omitted, this becomes a related conditional probability distribution (i.e., *P*(*L*|*L* > *L*_*min*_)). Shorter assemblies (particularly due to lower quality and read depth at repetitive loci) have a reduced overall number of sequenced repeats and a threshold for statistical (and potentially stochastic dynamical) noise at lower length. To ensure we are comparing estimates robust to statistical noise, we truncate each DRL above the lowest length bin containing less than 30 counts (roughly 1.5 in logarithmic space). This results in otherwise comparable normalized DRLs (assuming the same motif) with distinct truncation points based on non-normalized counts. Qualitative differences between the shape of the resulting normalized DRLs in remaining comparable length bins are indicative of differences in the evolutionary parameters (e.g., mutation rate, selection, etc.) or systematic error profiles (or both) between compared assemblies.

In addition to normalization for empirical comparisons, the empirical DRL and parameterized theoretical DRLs generated by our computational model were normalized by summing only over length classes above a specified minimum length to produce a comparable normalized DRL, conditional on *L* ≥ *L*_*min*_. This improved the summary statistic used to characterize differences between these distributions. Further details and justification for the specific choice of *L*_*min*_ are provided in *Bayesian inference procedure*, below. To make figures easier to interpret, normalized DRLs from the computational model were subsequently rescaled (where noted) to match the non-normalized counts for T2T-CHM13 by multiplying each normalized DRL by the sum of counts for bins *L* ≥ *L*_*min*_ in the T2T-CHM13 DRL.

### Bioinformatic estimation of substitution and indel rates

De novo mutation datasets were acquired as VCF files (or equivalent) from various published sources^83,84,85,86,87,88,89,90^, representing a total of 10,912 parent-child trios with available SNV data and 9,387 trios with available indel data. This dataset was compiled in McGinty and Sunyaev (2023)^91^ and comprised of all freely available trio samples at the time of analysis; samples from distinct VCFs were pooled to increase statistical power. We assumed that all individuals have the same underlying mutation rates. Variants were mapped to GRCh38 either in the original study, or subsequently, using pyliftover (pypi.org/project/pyliftover/). The average substitution rate was estimated to be 1.2x10^-8^, calculated as: number of substitutions / approximate number of sequenceable nucleotides in the diploid genome (see below) / number of offspring genomes in the dataset. We classified substitutions according to six categories based on trinucleotide context and the motif in question, as follows: for the example of A_n_ motifs, using B to represent non-A nucleotides, we determined rates (in parentheses) of ABB>AAB and BBA>BAA (4.58x10^-9^) representing repeat-lengthening events, AAB>ABB and BAA>BBA (7.74x10^-9^) representing repeat-shortening events, ABA>AAA (2.74x10^-9^) representing fusion events, AAA>ABA (4.35x10^-9^) representing fissions, BBB>BAB (3.80x10^-9^) representing the rate of A_1_ creation, and BAB>BBB (6.17x10^-9^) representing the loss of A_1_. Rates of substitutions of B which do not create an A were not estimated.

We calculated indel rates as a function of repeat tract length. Using positional information, upstream and downstream sequences for each event were pulled from the reference genome, under the assumption that the sequence of the parental genome is identical to the reference genome. For every focal motif, we used the reference sequence to determine tract length. Indel rates per tract length per motif were estimated by dividing by the number of repeats of that length, obtained by generating DRLs in a GRCh38 genome masked for low quality regions (see below). Each indel was classified as an expansion, contraction or non-motif insertion, additionally measuring how many motif units were added/removed in the event. We limited mutations in our computational model to +1/-1 unit changes in length at appropriate rates. We also measured the rate of indels for all B positions (with respect to each motif; A_n_ rates in parentheses), separately estimating the rates of BB>BBB (1.38x10^-10^), BBB>BB (4.37x10^-12^) and ABA>AA (2.76x10^-10^) events. Because B strings were not modeled as having length-dependent instability, we measured the average rate, i.e., the rate per unit.

Limitations of the VCF file format, namely the lack of any information at unmutated positions, forces the treatment of the pooled-trio VCFs as a complete record of variants in all individuals. At a coarse level, this problem was minimized by assuming that 100kb regions lacking any substitutions across the combined dataset suffer from regional mappability issues. These regions were masked in GRCh38 when estimating the denominator for rate calculations. At the fine level, this issue persists: mutations may have been filtered (prior to populating the VCF file) due to localized drops in sequencing quality, resulting in false negative calls and undercounting in the pooled estimates. This results in an underestimate of mutation rates, because counts from GRCh38 used in the denominator remain static. This may particularly affect estimation of instability rates as repeat tract length increases, because long repeats are known to interfere with several facets of the sequencing and bioinformatic processes^35,36,37,38^. We believe this systematic error mode, leading to progressively more severe underestimation of instability rates with increasing tract length, is the underlying cause of non-monotonicity observed in these rate estimates (**Fig. S5**). Mononucleotide repeats may be especially susceptible to systematic rate underestimation, as they are among the most difficult motifs to sequence^38^.

The popSTR repeat instability dataset, representing 6,084 parent-child trios, was acquired from the supplement of Kristmundsdottir et. al. (2023)^46^. This dataset was incorporated into our inference due to the unique methodology, which provided high quality calls of mutations extending beyond short tract length repeats that allowed us to produce length-stratified rate estimates. Files ‘bpinvolved_extended’ and ‘mutRateDataAll.gz’ were downloaded from https://github.com/DecodeGenetics/mDNM_analysisAndData. Due to our focus on uninterrupted repeats, we measured the longest contiguous repeat tract within the provided coordinates for each event. We limited the dataset to loci where the popSTR-reported reference tract length agreed with our own measurement in GRCh38. The ‘bpinvolved_extended’ file contains a mix of phased and unphased data; where the parental length for a given mutation was not assigned by phasing, we assumed that it originated from the parental copy which minimizes the difference in tract length between the proband repeat and any of the parental repeats. Skipping this phasing step and assuming that all events originated from the reference length allele (but retaining the size and direction of the event), as we do for the pooled-trio dataset, results in relatively minor differences in counts per length bin. The ‘mutRateDataAll.gz’ file contains information on the number of trios where all three samples passed sequencing quality filters at a given locus, and the length of the repeat tract at each locus in GRCh38, but lacks information on the parental genotypes for each of these loci (i.e., the file does not report pass/fail counts stratified by parental tract length). For the denominator of the popSTR mutation rates, we thus generated a distribution of passing counts (using the reference length for each locus), multiplied by two parental alleles. This assumption leads to some amount of misestimation of rates: loci containing long repeats show higher tract length variance in the population (due to higher instability rates), and thus individuals are more likely to differ from the reference genome. It is unclear whether a related effect (owing to the absence of loci with reference tract lengths below 10 nt in the popSTR dataset), or some other unknown error mode, is responsible for apparent overestimation of popSTR-based rates at shorter tract lengths where direct comparisons to reliable pooled-trio estimates are possible (**Fig. S5**).

We note that the popSTR dataset differs from the pooled trio data in several aspects: the popSTR caller provides no estimates for reference tract lengths below 10 nt; mutations are classified by length change, failing to distinguish between expansions and non-motif insertions; and, due to data access limitations, we were unable to assess the nature and magnitude of potential systematic errors detailed above. These distinctions precluded direct merging of instability rate estimates with the pooled trio data; popSTR-based estimates were instead incorporated into our inference by informing the prior (see below). Insertion rates, which are the sum of expansion and non-motif insertion rates, were used as a surrogate for expansion rates under the assumption that non-motif insertion remains far more infrequent than expansion (consistent with estimates from the pooled trio data at roughly 1% of total insertions).

For substitution and indel rate estimates based on either the pooled-trio or popSTR datasets, we calculated confidence intervals based on 200 Poisson samples of the mutation counts, removing the top 5 and bottom 5 values per length bin (see **Figs. 3b, S5**). We note that error bars provided on each estimate represent only Poisson distributed statistical error bars associated with point estimated counts in the numerator of each rate and are therefore subject to the above and any additional systematic errors underlying variant calling.

### Computational modeling of repeat length dynamics

We used a custom-written script in Python 3 that models repeat dynamics by directly manipulating the distribution of repeat lengths. We simultaneously tracked and manipulated the length distribution of B strings. As detailed above, we assumed a binary genome consisting only of A and B sites, where A is a repeat unit and B represents any non-A unit; as a result, B strings do not *a priori* represent repetitive sequences. Mutations are applied in aggregate such that, in each generation, repeats transition between integer length bins according to rules associated with each mutational process, while the B distribution is updated accordingly (e.g., a substitution that lengthens a repeat simultaneously shortens a B string). Mutation rates were restricted to be sufficiently low to model only a single mutation event per repeat per generation. The non-normalized distribution was evolved and subsequently normalized to create a probability distribution for comparison to empirical data. This approach is far more computationally efficient than simulating an entire genomic sequence, subsequently applying mutations and generating a distribution; computational time in our script scales with the number of length bins rather than with the length of the genome. Tracking only the distribution discards information about the location of particular mutations, instead generating an expected number of mutations for each category per length bin per generation. Except where specified, we used a deterministic approximation to assess the behavior of the expectation value of each bin as the distribution evolves toward steady state via repeated application of the mutation kernel. To understand the impact of stochastic fluctuations on the steady state distribution, we additionally implemented a model that represents fluctuations by Poisson sampling the expected change to each length bin per generation. We model stochastic fluctuations around the applicable rates by sampling mutational counts, but without constraining individual transitions (i.e., a net number of mutations may leave a given class, but the number introduced elsewhere, as a result, is appropriately distributed only on average due to an independent sampling procedure). All subsequent analyses were performed using the deterministic results, as modeling independent fluctuations in each bin showed no qualitative differences (**Fig. S13e**).

Mutations affect the distribution via the following well-defined rules for substitutions and indels (see **Fig. 2** for illustration). These rules assume that each mutation adds, subtracts or substitutes a single, complete repeat unit (i.e., the most prevalent class of length changes, seen in **Figs. 3a, S4**) . Using the example of a repeat of *L*=6, a lengthening substitution subtracts one count from the *L*=6 bin and adds one to the *L*=7 bin. A shortening substitution subtracts one from the *L*=6 bin and adds one to the *L*=5 bin. A substitution causing repeat fission subtracts one from the *L*=6 bin and adds two new repeats, either one *L*=1 and one *L*=4, or one *L*=2 and one *L*=3 (when evolving the distribution as a whole, both occur simultaneously with appropriate relative rates). The reverse process of fission is fusion, in which an L=6 repeat can be generated by fusing an *L*=1 with an *L*=4, or by fusing one *L*=2 and one *L*=3 repeats, while the mutated B unit is replaced with an A unit and added to the repeat length. Lengthening and shortening substitutions act locally (i.e., counts leave the *L* bin and move to the adjacent *L*+1 and *L*-1 bins, respectively). Substitution of an *L*=1 in the A distribution also corresponds to fusion of B strings; the reverse, i.e., substitution of a length one B string, generates fusion in the A distribution. Fission and fusion substitutions inherently act non-locally in length space: fission results in the loss of one count in the *L* bin and gain of two counts that are evenly distributed across all bins of length <=*L*-2; fusion evenly subtracts two counts from bins <=*L*-2 to add a count to *L*. The net effect of substitutions conserves the total length of the genome, i.e., the sum of the length of all A repeats plus the sum of the length of B strings remains constant under substitutions alone.

The rates of lengthening, shortening, fission and fusion substitutions per generation are separately estimated using the three-unit context: BBA>BAA (or ABB>AAB) for lengthening substitutions, AAB>ABB (or BAA>BBA) for shortening substitutions, AAA>ABA for fissions, and ABA>AAA for fusions. All substitution rates were assumed to be independent of repeat length, based on our previous observations showing little to no rate increase with increasing repeat length^91^. The target size for lengthening substitutions is two per repeat (i.e., the two sites adjacent to each repeat boundary). Likewise, the target size for shortening substitutions is also two per repeat, representing the two boundary units of the repeat (assuming *L*>1). The target size for fission substitutions is *L*-2 per repeat, representing all non-boundary units within the repeat. The target size for all fusion events is proportional to the *L*=1 count of the B distribution. Equations governing these processes are described in detail in **Supplementary Note SN1**.

Indel mutations operate under an analogous logic, but with a few important distinctions. Indels, by definition, do not conserve the length of the genome. Expansions and contractions act strictly locally, but the location of the event is indistinguishable within the repeat, affecting any of the units rather than just the boundaries; this results in a per-repeat target size *L* for these mutations, rather than 2. Non-motif insertions (i.e., AA>ABA) cause fission, resulting in the loss of one count in the *L* bin and gain of two counts that are evenly distributed across all bins of length ≤*L*-1; deletion of a B string of *L*=1 (i.e., ABA>AA) causes fusion, which evenly subtracts two counts from bins of length <=*L*-1 and adds one count to bin *L*. Indel rates for expansions and contractions are incorporated in a length-dependent manner, described above, in contrast to substitution rates. We did not model length dependence for B indels, as most B strings represent a combination of nucleotides and not necessarily STRs with any biological relevance. This assumption should not impact the evolution of the A distribution after normalization, which is only coupled to the *L*=1 class of the B distribution; this length class is dominated by substitution rate dynamics and not subject to repeat instability.

### Time-rescaling using a constant speed-up factor

Due to the large number of iterations required to reach a steady-state DRL, propagating the mutational process directly was computationally prohibitive. Instead, we approximated the DRL by first rescaling time by multiplying all mutation rates by the same constant 10^*r*^ such that each iteration represents 10^*r*^ generations of evolution for a total of *T*=10^*Ir*^ generations (assuming a constant *r* at all time points); *r* was limited to integer values for convenience. This defines a set of time-rescaled substitution, expansion, contraction, and non-motif insertion rates as a function of length.

Due to the rapid growth of instability rates with increasing length, these rates quickly saturate, reaching probabilities of one at some length *L*. To avoid multiple mutations per repeat per iteration, we defined a saturation length *L*_*max*_ as the length at which the sum of all mutation rates first exceeds 0.1 (e.g., *L*_*max*_ →∞if the sum of rates remains below 0.1 at all lengths). *L*_*max*_ thus demarcates the linear mutation regime, below which multiple mutation events remain rare. In addition to dependence on the instability rate parameters, *L*_*max*_ is dependent on *r*: increasing the speed-up factor increases mutation rates by 10^*r*^, which decreases *L*_*max*_. To ensure reasonable computational time, *L*_*max*_ was limited to a maximum of 200 (i.e., min{*L*_*max*_,200}, computed separately for *A*-repeats and *B*-string lengths), which extends well beyond the empirical DRLs. To prevent loss of mass associated with the finite length grid, we imposed a reflective boundary condition at *L*_*bound*_ = min{*L*_*max*_,200}, (i.e., all transitions from lengths *L* < *L*_*bound*_to *L* ≥ *L*_*bound*_ were assigned to *L*_*bound*_). This results in artefactual behavior near the boundary but provides a reasonable approximation when the expected number of counts drops below one at lengths far below *L*_*bound*_ Substantial counts at the boundary are indicative of unrealistic distributions (often associated with diverging total genome size), provided *L*_*bound*_is sufficiently far from the maximum well-populated lengths in the comparable empirical distribution.

For a given parameterization, producing a grid of DRLs requires choosing a constant speed up *r* and the boundary length *L*_*bound*_appropriate for each parameter combination (the required computational time is largely determined by the number of parameter combinations with the lowest value of *r*). This procedure can be used to produce a coarse grid of parameters (e.g., for comparison to alternative approximations) but proved computationally prohibitive for the dense grid needed for inference.

### Step-wise speed-up procedure

To produce finer grids of DRLs (for several parameterizations), we implemented a procedure that reduces overall computational time, while producing approximately the same DRL as that under a constant speedup (described above). This procedure models the evolution of the DRL by performing several, discrete phases of evolution, each with successively smaller time-rescaling factors 10^*r*^. Each stage is allowed 10^6^ iterations of evolution under the specified *r* (and the associated reflective boundary at *L*_*bound*_ = min{*L*_*max*_,200} for each parameter combination); *r*=3 for the first stage, and is reduced to *r*=2, 1, and 0 for subsequent stages. *L*_*bound*_is altered at each stage to maintain the linear mutation regime (i.e., ensuring *L*_*max*_ ≥ *L*_*bound*_). In total, parameter combinations can experience up to four stages (equivalent to 4 × 10^6,^ iterations, or 1.111 × 10^9^ generations).

For computational efficiency, we first separated parameter combinations that rapidly equilibrate in a single stage of 10^6^ iterations under a sufficiently large rescaling factor *r* ≥ 3 (easily identified by *L*_*max*_ ≥ *L*_*bound*_ = 200; equivalent to propagation using a single, constant speed-up factor). These were each run at the largest allowed integer *r* for the equivalent of 10^9^ generations and removed from the grid. All other parameter combinations were subjected to several stages with progressively decreased *r*; after each stage, parameter combinations deemed equilibrated were removed from the grid (again identified by *L*_*max*_ ≥ 200, determined by the preceding *r*-rescaled instability rates). For parameter combinations with *L*_*max*_ < 200 in the absence of any speed-up factor (*r*=0), counts in all length bins between *L*_*max*_ and 200 were set to zero prior to analysis of the DRL.

To ensure that the multi-step procedure provides a reasonable approximation to the DRL produced under a constant speed-up, we compared inference results over a coarse grid of parameter combinations and found negligible differences (**Fig. S13**). Intuitively, this procedure takes advantage of the faster mutation rates at longer lengths, which equilibrate on much more quickly than shorter length bins.

### Initial conditions at t=0

The computational model was initialized with an initial distribution that is approximately geometric (created by propagating substitutions alone, setting all instability rates to zero) for the equivalent of 10^10^ generations (using the largest allowable rescaling, *r* = 5). Using this approximation to the substitution-only steady-state as a pre-simulation substantially reduces equilibration times because the lowest length bins (i.e., those dominated by substitutions) require the most time to equilibrate due to low mutation rates.

Although the eventual steady-state DRL should not depend on the initial state, the choice of initial distributions can dramatically affect equilibration times. We confirmed that the final timepoint DRLs are effectively independent of choice of initial condition by comparing the results of two distinct initial conditions with similar equilibration times (geometric vs. geometric plus uniform; Fig. S12), finding only minor differences in the deterministic late-time distribution.

### Computational model inputs and outputs

The script relies on the following as inputs: an initial distribution for A and B (i.e., motif and non-motif) repeat lengths, per-target substitution rates in three-unit context, and per-target mutation rate curves for expansions, contractions and non-motif insertions. Substitution rates and length-dependent indel rate curves are imported from external files (see above for estimated substitution rates, below for generation of parameterized rate curves); these files, along with the initial repeat length distribution table, can be replaced with appropriate tables for other purposes, if desired. This table must specify rates for each mutational process at all lengths intended to be computationally modeled (i.e., from 1 to *L*_boundary_). For normalized length distributions that reach steady state, the initial distribution can be chosen arbitrarily, in principle, but any specific choice affects equilibration time; due to equilibration time differences, minor differences between the deterministic late time distributions arise from distinct initial conditions (see **Fig. S13** for comparison between two initial distributions).

Stochastics can be introduced using a command line option to model fluctuations in the mutational process; the number of mutations in and out of each length bin are separately Poisson sampled (using *numpy*.*random*.*poisson*) around the expected number of mutational counts in each iteration.

After each run, we output a file containing repeat length counts reported at various time points to show the temporal evolution of the distribution. We subsequently normalized the resulting distributions by dividing each length bin by the total number of repeats in the distribution (see Methods on normalization).

The relative contribution of each mutational force was assessed by producing a single-generation plot of the transitions in and out of each length bin at the final time point (i.e., once steady state was reached, if applicable). To produce these plots (see **Fig. 6**), we applied the mutation kernel for a single generation and separately computed the number of fission, fusion and local changes for substitutions and indels. For each length bin, the magnitude of total flux in and out was normalized to one. Length bins that have equilibrated should contain equal fluxes in and out; steady state occurs only when all bins show equilibrated fluxes.

### Bayesian inference procedure

Given our observations indicating a stable distribution of repeat lengths over phylogenetic time scales, we sought to identify mutation rates capable of explaining this observation. To study the extent to which mutational processes alone can recapitulate the repeat tract length distribution, we constructed a Bayesian inference framework to compare models (i.e., parameterizations) of the length-dependent rates of repeat instability. Each parameterization describes the length-dependent rates of expansion and contraction as following a simple functional form; as discussed above, substitution rates are assumed to be length independent in all cases. Within the Bayesian framework, a prior probability distribution on the parameter space is specified (several priors were used for interpretation) and used to weight the likelihood to calculate a posterior probability that a given parameter combination accurately describes the length dependence of the repeat instability rates. In the present setting, the likelihood is constructed by comparing the empirical repeat tract length distribution to the late-time distribution generated by computationally modeling a given parameter combination. Due to the analytic intractability of this likelihood, we used Approximate Bayesian Computation (ABC)^56^ to approximate the posterior probability distribution, which avoids specifying the likelihood explicitly. Additionally, length bins are presumably correlated due to the complex mutational transitions underlying the distribution, complicating naïve construction of the likelihood. In contrast, ABC-based inference circumvents this issue by approximating the posterior in terms of summary statistics that appropriately characterize the DRL (see discussion of summary statistics below). After specifying summary statistics, the late-time distribution for each parameter combination was summarized for comparison to the empirical distribution (e.g., mononucleotide A repeat tract lengths in T2T-CHM13). For each parameterization, we specified a discrete grid of parameters for comparison to the empirical distribution, the result of which was weighted by the prior probabilities for those parameters (equivalent to randomly sampling the prior as prescribed in ABC^56^; see below for specific priors used) to compute the posterior.

We chose to use the Kullback-Leibler (KL) divergence, a well-established statistic for distribution comparison, to characterize the difference between the empirical and parameterized repeat tract length distributions. The KL divergence quantifies the extent to which each parameterized distribution diverges from the empirical distribution and was calculated for all parameter combinations (denoted θ below) on the discrete grid using the following definition.

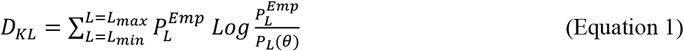

We note that comparing the empirical distribution to itself results in a divergence of zero such that the KL values are equivalent to the difference between the modeled and empirical distributions (i.e., 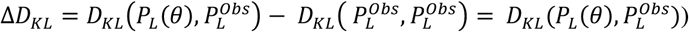. To define a cutoff for ABC rejection, we estimated the divergence between the human empirical distribution and the ensemble of primate genomes using the same statistic. Under the assumption that primates evolved towards the same steady state (i.e., the mutational parameters remain constant across the phylogeny), we proceeded under the assumption that differences between the repeat tract length distributions in distinct species are due to a combination of stochastics and bioinformatic errors due to the lower coverage and short read technologies used to assemble primate reference genomes. Due to the difference in assembly lengths, we added a pseudocount of one to all length classes in all species to avoid divergence of the statistic and confirmed that our results were qualitatively independent of the choice of pseudocount between 0.01 and 100. L_*max*_ was set the longest modeled length bin, *L*=200. We set the lower bound to *L*_*min*_ = 4 to ensure that the ordering of *D*_*KL*_ statistics computed for all primates remains roughly consistent: setting *L*_*min*_ ≥ 4 resulted in the smallest values for the human HG38 reference (which was not included in subsequent analyses) and the largest values for the most divergent primates (i.e., those on the loris branch). We used the range of 36 computed primate KL values to very roughly define a rejection threshold by throwing out the largest 2 values; we considered the remaining 34 values (i.e., roughly, the closest 95% of ranked primates) and used this to approximate the variance of *D*_*KL*_ (i.e., 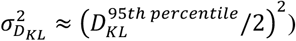 associated with stochasticity and sequencing errors.

We approximated the posterior probability (up to a normalization constant) following the prescription in Wilkinson^56^ wherein ABC is applied with a soft rejection threshold by rejecting values of *D*_*KL*_ (*θ*) with the following probability based on the primate-estimated variance.

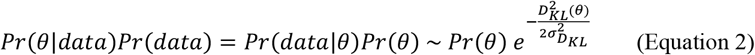

Here, *Pr*(*θ*|*data*) is the posterior probability distribution over the grid of parameter combinations *θ, Pr*(*θ*)is the prior, and the Gaussian falloff is the rejection probability (up to normalization) for a given parameter combination. This soft-rejection procedure provides a slightly better approximation for the posterior distribution than rejection with probability one (e.g., reject all parameter combinations with 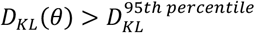, roughly corresponding to the primate-estimated 95% confidence interval). Additionally, Wilkinson argues that the Gaussian rejection probability quantifies model misspecification inherent in the procedure, as all parameterizations (i.e., models) employed herein are imperfect approximations of the true instability rate parameters.

### Model comparison using Bayes factors

To assess the relative explanatory power of each parameterization *M*_*θ*_ modeling the length dependence of repeat instability, we computed a Bayes factor for each model *BF*(*M*_*θ*_) using the definition below.

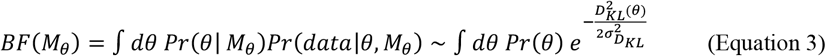

Here, the right-hand side is our previously computed approximation to the posterior, integrated over the parameter space for a given model. As we were only interested in the relative Bayes factor between models, proportionality constants can be ignored, including the overall normalization and an assumed uniform prior over model space. The Bayes factor for a model naturally controls for the number of degrees of freedom in each parameterization because integration over the weighted posterior is performed in parameter spaces with differing dimensionalities. Once computed, models were compared by interpreting the Bayes factor ratio (BFR) as indicative of the relative statistical support between two parameterizations of interest.

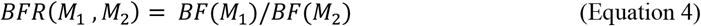

We then used the Jeffrey’s scale to interpret the strength of statistical support for each model.

### Parameterizations of repeat instability rates

We tested several parameterizations to assess consistency with the empirical distribution of repeat tract lengths. We focused on mononucleotide A repeats, as both the distributions and rate estimates were supported by the most empirical data. For computational convenience, we defined a sequence of nested parameterizations (see **Fig. S15**) that could be computed simultaneously across the grid of the parameter combinations under the model with the largest number of degrees of freedom (DoF). To define the most general set of length-dependent instability rate models, we parameterized expansion and contraction rates, *ϵ*(*L*) and *κ*(*L*), respectively, as independent power law functions at all lengths L>8.

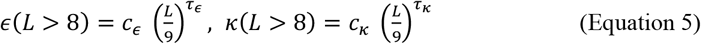

Rates for L=1—8 were taken directly from empirical rate estimates; the expansion and contraction rates at longer lengths were parameterized in terms of *c*_*ϵ*_ and *c*_*κ*_, which denote their respective values at L=9 (i.e., the first parametrized length bin). Guided by our empirical estimates, we assumed that the rate of non-motif insertion ι(*L*) is directly related to the rate of expansion with the same length dependence at 1% of the rate (i.e., ι(*L*) = *ϵ*(*L*)/100 at all lengths. This results in instability rates characterized by four independent parameters (*c*_*ϵ*_, τ_*ϵ*_, *c*_*κ*_, τ_*κ*_). We then constructed a series of nested lower-dimensional models for comparison. A natural way to reduce the dimensionality of the parameter space is to introduce symmetries corresponding to *c*_*ϵ*_ = *c*_*κ*_ and/or τ_*ϵ*_ = τ_*κ*_. The simplest model assumes fully symmetric expansion and contraction rates (i.e. both *c*_*ϵ*_ = *c*_*κ*_ ≡ *c* and τ_*ϵ*_ = τ_*κ*_ ≡ *τ*) with a two-dimensional parameter space (*c, τ*). We treat this as a null model corresponding to a frequently discussed^8,20,23,28^ biological interpretation of repeat slippage. The parameter space can be reduced to three DoF by restricting to either *c*_*ϵ*_ = *c*_*κ*_ or τ_*ϵ*_ = τ_*κ*_, which need not have straightforward biological interpretations. We constructed an additional 3 DoF model parameterized by(*m*, τ_*ϵ*_, τ_*κ*_) by treating the expansion and contraction rates at L=9 as increased by a common multiplier *m* relative to their values at L=8 (i.e., *ϵ*(9) = *m ϵ*(8), *κ*(9) = *m κ*(8)). For computational expediency, we embedded this model within the four-dimensional grid of parameters by appropriately choosing intervals for *c*_*ϵ*_ and *c*_*κ*_ when defining the grid discretization.

We used two distinct, non-nested parameterizations for subsequent analyses. To test the reliance of our inference on the functional form of the length dependence (i.e., power-law parameterization), we defined an additional parameterization by replacing the length dependence for L>8 with logarithmic growth in the following form:

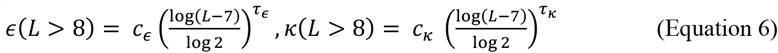

Here, the dependence on Log 2 ensures that *c*_*ϵ*_ and *c*_*κ*_ parameterize the values at L=9 of the expansion and contraction rates, respectively. Under this parameterization, empirical estimates were again used for all lengths *L* ≤ 8. This functional form retains monotonicity while growing more slowly at longer lengths to model a saturation-like effect.

We analyzed a second version of the power-law parameterization that extends the functional form to all lengths such that the rates are fully independent of empirical estimates. We re-parameterized the functional dependence in terms of the parameters (*λ*_*ϵ*_, *τ*_*ϵ*_, *λ*_*κ*_, *τ*_*κ*_), where *λ*_*ϵ*_ and *λ*_*κ*_ correspond to the length at which each instability rate exceeds the relevant substitution rate (i.e., *ϵ*(*λ*_*ϵ*_) = *μ* and *κ*(*λ*_*κ*_) = *ν*).

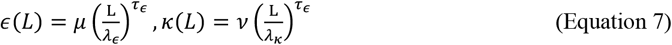

Here, *μ* and *ν* are point estimates of the average lengthening and shortening substitution rates, respectively, for a given motif; note that this parameterization is defined at all tract lengths (including *L* ≤ 8). This allowed us to directly infer the length scale of the instability-substitution rate crossover and assess the extent to which our inferences from the above parameterizations rely on direct use and accuracy of empirical rate estimates below L=9. After confirming that expansion-biased parameters do not approach steady state DRLs, the parameter space was further limited to asymptotically contraction-biased parameter combinations (i.e., with *τ*_*ϵ*_ > *τ*_*κ*_) to limit computational time.

The functional form of each aforementioned parameterization is specified in Table 1.

### Construction of prior distributions

For each parameterization, we constructed an uninformative prior by treating each parameter combination as equally probable with probability equal to *1/n*, where *n* is the number of computationally modeled points on a discrete grid. We next generated informative priors using approximations derived from our empirical estimates of the expansion and contraction rates. For power-law parameterizations that include empirical estimates at low lengths (Models 1—5 in **Table 1**), we performed a linear fit to the popSTR rate estimates (at lengths *L*=11—29) in log-log space to estimate parameters of best-fitting power laws. Curve fitting was performed in Python 3 using *scipy*.*optimize*.*curve_fit()* with the sigma option to specify an array of approximately symmetric log error bars (i.e., approximating a rescaled Poisson as log-normal). We note that Poisson regression does not appropriately model statistical noise due to target size rescaling when estimating rates from mutational counts. We fit using rate estimates at all available lengths. However, we artificially inflated the variance at lengths above and below *L*=13—21 to model potential systematic errors that generate observed non-monotonicity, likely due to miscalling at the shortest and longest lengths accessible to popSTR. The optimization package produced a covariance matrix for the best-fit line expressed in terms of the slope and intercept in log-log space. We used this covariance matrix to approximate lines representing the 95% confidence bounds around the best-fit line. Using these lines, we estimated the value of the best-fit line and standard deviation at *L*=9. Assuming no correlation between expansion rate and contraction rate parameters, we used these values to approximate a block diagonal covariance matrix for the parameters (*c*_*ϵ*_, *τ*_*ϵ*_, *c*_*κ*_, *τ*_*κ*_) in the four-dimensional model. We then inflated the variance by a constant (100-fold for ‘restrictive’ informative prior; 1000-fold for ‘permissive’ prior)_and used the rescaled covariance matrix to define a multivariate normal distribution centered at the point-estimates for the best-fit parameters. The informative prior for the four-parameter model was constructed by normalizing this multivariate normal over the discrete grid. Analogous priors for nested models were defined by restricting to the appropriate subset of parameter space, maintaining the relative weights specified by the normal distribution, and normalizing by the number of discrete grid points in this subset.

The approximate nature of our ABC-based inference procedure prohibited construction of strictly uninformative priors (i.e., Jeffreys priors) for fair comparison between models with differing parametric functional forms. We instead treated the uniform prior as naively uninformative; however, despite the similarity of their parameters, we caution that uniform priors are inequivalent for distinct functional forms. To facilitate very rough model comparison, we constructed a restrictive prior for the logarithm-based model by again fitting the functional form to popSTR rates using *scipy*.*optimize*.*curve_fit()*. This produced a point estimate of the best-fit parameters that was used to specify the mean of a multivariate normal distribution. To attempt to define a multivariate normal very roughly comparable to the priors for the power-law parameterizations, we defined the normal distribution in terms of the covariance matrix estimated under the four-parameter power-law model. This comparison relies on the fact that both parameterizations use the same parameters to represent nearly identical quantities (i.e., parameters *c*_*ϵ*_, *τ*_*ϵ*_, *c*_*κ*_, and *τ*_*κ*_ define constants and exponents in the same way). The procedure described above was then used to define restrictive and informative priors (with 100-fold and 1000-fold inflated variances, respectively) for two- and four-dimensional logarithm-based parameterizations from four-dimensional multivariate normal distributions with appropriately shifted means.

To construct priors for pure power-law models, we again used a uniform prior over the discrete grid to define a naively uninformative prior over the parameter space. Informative priors were defined by again using *scipy*.*optimize*.*curve_fit* to estimate best-fit parameters and a covariance matrix from empirical instability rate estimates. Unlike the previous models, rates at all lengths (including *L* ≤ 8) were fully parameterized by **Equation 7** (and **Table 1**); data from both the pooled trio and popSTR datasets was used to estimate the mean and covariance matrix by fitting the functional form to expansion and contraction rate estimates for *L* = 4 − 15. The covariance matrix was again inflated by a factor of 100 and used to define a four-dimensional multivariate normal around the point-estimated mean values of (*λ*_*ϵ*_, *τ*_*ϵ*_, *λ*_*κ*_, *τ*_*κ*_).

### Calculation of expectation values from posterior probability distribution

In addition to identifying maximum posterior probability parameter combinations, we used the posterior distribution to weight various quantities to calculate their expectation values. Expectations of an arbitrary parameter-dependent function *f*(*θ*) were computed as follows.

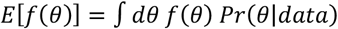

Here, the ABC-approximated posterior was used for *Pr*(*θ*|*data*) after normalizing over the discrete computational grid of parameters *θ* such that *E*[1] = ∫ *dθ Pr*(*θ*|*data*) = 1. We used this to compute: the length dependence of the posterior-weighted repeat instability rates *E*[*ϵ*(*L*; *θ*)] and *E*[*κ*(*L*; *θ*)] for comparison; and the posterior-weighted distribution of repeat tract lengths *E*[*P*(*L*; *θ*)] (see **Fig. 5**).

### Analytic modeling of repeat length dynamics

To better understand the underlying dynamics that generate with the genome-wide repeat length distribution, we attempted to analytically model the effect of each mutational type on the number of repeats at a given length *L* from first principles. We were interested in describing the steady state distributions that emerge for a subset of parameter combinations, as seen in the results of our computational model. Our goal was to capture the balance between relevant mutative forces, which can vary by repeat length, by writing an appropriate approximation to the steady state equation; the solutions to these equations describe the shape of the normalized repeat length distribution, *P*(*L*), restricted to the regime of validity of each approximation. Within this section, we have used the notation *P*_*L*_ to represent the distribution *P*(*L*) more compactly when detailing the relevant equations. Each parameter combination defines a functional form for the per target (i.e., per unit) expansion, contraction, and non-motif insertion rates at lengths 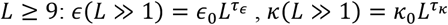, and 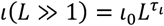, respectively, where the constants *ϵ*_0_, *κ*_0_, and ι_0_ are set by the empirical value of these rates at *L*=8 and the multiplier *m* (noting that we set *τ*_ι_ ≡ *τ*_*e*_ to limit the number of free parameters; see inference Methods). Again, these length-dependent rates, in either discrete or continuous form, are denoted with a subscript *L* (e.g., *ϵ*_*L*_ ≡ *ϵ*(*L*)) in this section for brevity. For substitutions, we refer herein to rates *μ* ≡ *μ*_*A*→*B*_ and *ν* ≡ *μ*_*B*→*A*_ for lengthening and shortening mutations, respectively, but later specify separate mutation rates based on three-unit context (e.g., *μ*_*ABB*→*AAB*_) when comparing directly to computational model results. While the mutation rates may be well defined by these rates, the combined effect of substitutions and indels on the repeat length distribution requires a description of a number of complicated behaviors, including both local and non-local transitions between lengths across the distribution, non-conservation of the number of repeats due to fission and fusion, and non-linear dependence on the state of the distribution due to fusion (i.e., the generic dynamics are non-Markovian). As a result, our aim was not to describe an exact solution, but instead an expression for the effective dynamics that dominate the maintenance of the distribution in steady state, specifically in the asymptotic regimes associated with the shortest and longest length repeats. Note that this analytic description was motivated by and is strictly applicable to mononucleotide repeat dynamics, where the species of repeat length-changing mutations are fewer, but the conceptual findings may be generalizable to longer motif repeats (**Fig. S3**).

### Short repeat regime

First, we focused on the regime of asymptotically short repeats, as their behavior is more straightforward. By assessing the relative rates of substitution and indel processes in the estimated per-target rates (**Fig. 2b**), one can immediately see that substitutions must dominate the dynamics for the lowest length repeats. Short repeats can be characterized by a straightforward balance between opposing types of substitutions, µ and *ν*, which is equivalent to sequence evolution under a two-way point mutation process. At steady state, the resulting distribution is equivalent to the probability of randomly assembling specific strings of length *L* when the whole genome is randomly sampled between A and B bases with probability *p*_*A*_ = *μ*/(*μ* + *ν*) and *p*_*B*_ = *ν*/(*μ* + *ν*), respectively. The frequency of a length *L* string of A’s (i.e., an A repeat) is geometrically distributed in proportion to *p*_*A*_^*L*^ (i.e., sampling an A, *L* successive times).

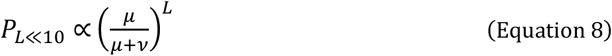

Here, we have omitted a normalization constant that determines the relative weight of this geometric distribution to the weight of the long repeat tail. For comparison to the computational model (or the empirical distribution), we fixed the normalization constant using the mass of the *L* = 1 bin. The approximation that the effects of expansion, contraction, and non-motif insertion are negligible breaks down at a length determined by the estimated relative rates in **Fig. 3b**; the regime of validity for this approximation extends roughly to lengths of order *L* = 10.

### Long repeat regime

The dynamics of long repeats, i.e., for asymptotically large repeat lengths *L* ≫ 1, the analysis is complicated by the numerous length-dependent (and parameter-dependent) forces that can potentially contribute to stabilizing the distribution. While expansion and contraction describe inherently local transitions from *L* to *L* + 1 and from *L* to *L* − 1, respectively, the effects of non-motif insertions and substitutions on extended repeats are not strictly local. To model this regime, we first wrote a finite difference equation that describes the change in the distribution in a single time step Δ*t*: Δ*P*_*L*_ ≡ *P*_*L*_ (*t* + Δ*t*) − *P*_*L*_ (*t*), where P_*L*_ (t) = *P*(*L, t*; µ, *ν*, ϵ_*L*_, κ_*L*_, ι_*L*_) is implicitly dependent on the length scaling of each rate (see **SN1**). From this discrete equation, we derived a partial differential equation (PDE) in the large-length continuum limit Δ*L* = 1 ≪ *L* that approximates the dynamics in the large length regime (derivation provided in **SN1**). This PDE includes explicit terms depicting the combined local effects of repeat instability due expansion and contraction, each occurring at distinct length-dependent rates, and the separate effects of repeat fission and fusion, each introducing an integral that captures the aggregate effects of non-local transitions in length. Expansion and contraction collectively generate both symmetric (i.e., bidirectional) and asymmetric local length transitions, which correspond to a diffusion term represented by a second derivative and directional flux term expressed as a first derivative, respectively, each appropriately accounting for length dependent rates.

While local effects from substitutions and non-motif insertions exist (specifically, transitions *L* → *L* + 1 or *L* → *L* − 1), as well, they are negligible in comparison to expansion and contraction due to their low relative rates at long lengths and finite target size of two per repeat. Fission due to substitutions and non-motif insertions were both accounted for as separate non-local contributions to the change in *P*_*L*_. Importantly, the probability of fission due to substitution is proportional to the target size (*L* − 2) ≈ *L*; for insertions, the rate itself harbors an additional length dependence such that the per-repeat rate of fission scales as 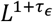. As a result, the relative importance of fission compared to local contributions is highly dependent on the parameters *τ*_*ϵ*_ and *τ*_*κ*_; similarly, the relative importance of substitution- and insertion-based fission are parameter dependent due to distinct dependencies on length. Thus, a unified description across parameter space requires the inclusion of fission in full form and captures all four mutational effects. While we were able to explicitly describe the integral effects of length changes due repeat fusion in the continuum (see **SN1**), the inherent non-locality is additionally complicated by the nonlinearity introduced by pairing two repeats randomly sampled from the distribution. To make further progress, we proceeded under the assumption that fusion remains subdominant at large lengths, which we confirmed via our computational model to be generically true across parameter space. Stochastic fluctuations in the mutation rates were omitted, resulting in a deterministic approximation for the expected repeat length distribution.

Next, we imposed the assumption of steady state (i.e., *dP*/*dt* = 0), reducing the PDE to an ordinary differential equation in length to solve for the shape of the distribution in equilibrium. Despite excluding complications from fusion, the remaining approximation to the steady state equation is, strictly speaking, a second order integro-differential equation, for which no explicit closed-form solutions exist. The following equation approximates the steady state dynamics in the absence of fusion (i.e., when fusion is subdominant). Here, ∂_*x*_ represents a derivative with respect to *x* (noting that partial derivatives with respect to *L* become total derivatives in steady state) and *P*_*L*_ is the steady state value of the continuous repeat length distribution at large length *L* ≫ 1 up to an overall normalization constant (along with an arbitrary constant set to zero). Again, all continuous functions describing mutation rates (e.g., ϵ_*L*_, κ_*L*_) are expressed here as per-target rates.

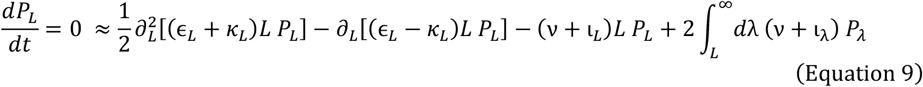

In order from left to right, the terms appearing on the right hand side describes: length-dependent diffusion (arising from local transitions due to expansion and contraction), a length-dependent local directional flux (due to the bias between expansion and contraction), a net loss of due to fissions that break up length *L* repeats (i.e., substitutions or insertions that interrupt the repeat sequence; referred to herein as *fission out*), and a net gain due fissions of repeats longer than *L* (referred to as *fission in*). Fission in represents the sole integral effect, which substantially complicates our analysis; elimination of the integral dependence is discussed below and results in a third order ordinary differential equation (ODE) that maps to this second order integro-differential equation.

### Contraction-biased rates stabilize the distribution

Importantly, we found that steady state could only be reached for the subset of parameter combinations with *τ*_*κ*_ > *τ*_*ϵ*_, corresponding to cases for which local transitions are asymptotically contraction-biased: 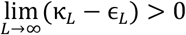 (note that the edge case where *τ*_*κ*_ = *τ*_*ϵ*_ is asymptotically expansion-biased based on observations at *L* = 8 and implications of our parameterization). We therefore denote this as the contraction-biased regime, which is characterized by defining the variable Δτ ≡ τ_*κ*_ − τ _*ϵ*_. When Δτ > 0, the distribution is stabilized at some arbitrarily large length *L* = *L*_*trunc*_ by sufficiently large contraction rates in excess of all processes that increase repeat length; a truncation of the distribution (i.e., when less than one repeat is expected in a genome of given size) occurs due to the more rapid increase of contraction rates than expansion rates that leads to contraction-biased dynamics at some point *L* < *L*_*trunc*_. The necessity of asymptotic contraction-bias contrasts the notion that length-dependent interruptions (due to substitutions and non-motif insertions) counteract expansion at sufficiently long lengths, stabilizing the distribution^45,49,55,58,59,60^ based on our estimated mutation rates, this effect does not lead to a steady state in the absence of contractions, as the per-repeat rate of expansions far exceeds that of repeat fission (i.e., interruptions) at long lengths. As discussed below, the length at which the contraction rate is equal to the expansion rate *L*^*^ (i.e., *L*^*^ is the unique length *L* ≥ 8 where κ_*L*_ = ϵ_*L*_, which may occur at non-integer values) is highly informative about the dynamics in each regime, as well as the behavior when all effects captured in **Equation 9** are simultaneously relevant; *L*^*^ is exponentially dependent on Δ*τ* and more weakly controlled by the multiplier *m*, notably occurring at the same length across lines of constant Δ*τ* in the parameter space (for a given *m*). For *m* > 2.5, the dynamics are nearly identical for parameter combinations with the same Δτ, effectively collapsing the (*τ*_*ϵ*_, τ_*κ*_) plane to a single dimension. The functional dependence of *L*^*^ on the parameters and further discussion is provided in **Supplementary Note SN1**.

### Effective equations approximating steady state dynamics

Given the complexity of **Equation 9** introduced by the nonlocal effects of fission, we first searched for subsets of the contraction-biased parameter space that could be well approximated under a further reduction of the dynamics. Such simplifications are, in principle, possible because the length scaling of each term in **Equation 9** is distinct; specifically, parameter combinations exist where the nonlocal behavior (i.e., the integral representing fission in) becomes subdominant and can be neglected in our analysis. Neglecting the integral results in a second order ODE approximation to the steady state equation. We identified two distinct dynamical regimes within the Δ*τ* > 0 region, which are each well-approximated by a subset of contributions that dominate the dynamics in their respective regimes of validity.

### Balance between local dynamics in the highly contraction-biased regime

For parameter combinations with very large positive values of Δ*τ* (i.e., for *τ*_*ϵ*_ ≫ *τ*_*κ*_), the dynamics are entirely dominated by the diffusion and local directional flux terms appearing in **Equation 9**, as the contraction rate quickly outcompetes both the rate of fission in and fission out. This results in an effective steady state equation dominated only by local transitions.

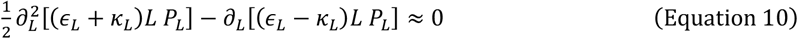

In this case, the contraction rate exceeds the expansion rate almost immediately above the short length regime (i.e., *L*^*^ is of order 10) such that the dynamics are effectively uniform across the long length regime. The long length tail of the distribution decays in a super-exponential fashion such that the truncation occurs at low values of *L*_*trunc*_ ∼ 20, which dramatically limits the lengths of repeats that occur in a genome of realistic size. In this regime, a further simplification leads to an approximate closed-form analytic solution for the rough asymptotic shape of the distribution, however this approximation is only valid near the truncation point and rapidly loses accuracy. A more general solution was obtained by numerically solving the effective steady state equation (**Equation 10**) for comparison to computational model results. To obtain numerical values, two additional constraints must be applied, as with any second order ODE, which conceptually correspond to an overall normalization constant (in this case, fixing the relative weights of the short length and long length distributions) and a linear coefficient that defines the relative weights of two real solutions, if both exist. These constraints can be imposed by fixing the value of the distribution at two specific lengths, *L*_1_ and *L*_2_, (i.e., fixing 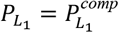 and 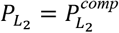, where 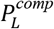 is the value of the computationally modeled distribution at length *L*), with both lengths chosen to lie long length regime *L* > 10 where the continuum approximation remains valid. For consistency, we chose to constrain the numerical solutions at the two lengths of theoretical interest in stable distributions: *L*_1_ = *L*^*^ (rounded to the nearest integer) and *L*_2_ = *L*_*trunc*_, both of which definitionally remain in the long length regime at a location with finite occupancy in a realistic genome and are well defined for all values of Δτ > 0. All numerical solutions were obtained using the *NDSolve* function in Mathematica 14.0^92^. Comparisons between computational model results and numerical solutions to **Equation 10** showed that this approximate steady state equation remains highly accurate across the Δτ ≫ 1 regime (see **SN1**).

### Relevant effects of fission out in the intermediate contraction-biased regime

We found that, at less extreme values of Δτ, roughly on the order of Δτ ∼ 1 (e.g., roughly 1.5 > Δτ > 0.7 for *m*=4), the integral contributions to **Equation 9** remained subdominant, but the effects of fission could not be omitted completely. In this regime, fission out non-negligibly impacts the dynamics, leading to an effective steady state equation that only omits incoming contributions from fission.

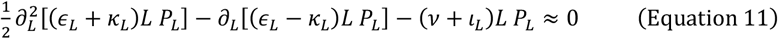

In this regime, contraction is aided by the length-reducing effects of fission out. However, the relevance of this contribution is limited roughly to lengths below *L*^*^ ; above *L*^*^, the distribution remains well-described by **Equation 10** (see **SN1**). This indicates that contraction is largely responsible for truncating the distribution, even when fission is involved in shaping the distribution. This defines a range of intermediate lengths below *L*^*^ with distinguishable dynamics from asymptotic lengths, but this range is limited by the relatively small values of *L*^*^ on the order of *L*^*^ ∼ 15 − 20. The approximation in **Equation 11** is again a second order ODE but is complicated by the introduction of an additional length scaling associated with substitution-based fission. However, even when substitution rates are negligible (e.g., for *m* ≫ 1), no exact solution could be found due to the generic power laws associated with our parameterization. For comparison to the computational model, numerical solutions were obtained by again constraining the solution at lengths *L*_1_ = *L*^*^ and *L*_2_ = *L*_*trunc*_. We found that the effective steady state equation (**Equation 11**) is a highly accurate approximation to the dynamics in this regime of moderate values of Δτ. Additionally, this approximation remains accurate at large values of Δ*τ* (i.e., **Equation 11** is applicable to the full subspace Δτ ≳ 1), as the approximation in **Equation 10** is nested in **Equation 11**; the latter includes the additional effect of fission out, which becomes negligible for Δ*τ* ≫ 1.

### Inclusion of the nonlocal dynamics in the weakly contraction-biased regime

For values Δτ ≲ 0.5 (roughly Δτ < 0.7 for *m*=4), the nonlocal effects described by the integral term in **Equation 9** become relevant to the maintenance of steady state. To further analyze this regime, we first eliminated the integral dependence by applying an overall length derivative to all terms on the right-hand side of **Equation 9** such that the equation becomes the following.

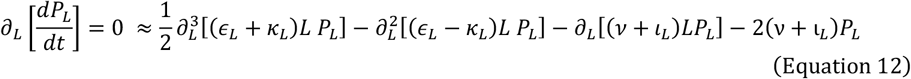

This third order ODE now represents a constraint on the net flux, which must equal a time-independent constant. This can be seen by swapping the order of the derivatives on the left-hand side of **Equation 12**: ∂_*L*_ [*dP*_*L*_/*dt*] = *d*[∂_*L*_*P*_*L*_]/*dt* = 0. Taking this overall length derivative maps the nonlocal contributions from the fission of all repeats longer than L to an effectively local boundary effect on the net flux ∂_*L*_*P*_*L*_ through length *L*. However, this is not equivalent to steady state until applying an additional constraint that this net flux vanishes (i.e., the special case where the constant is zero, ∂_*L*_*P*_*L*_ = 0). Obtaining numerical solutions to this third order ODE requires three constraints, which now includes the constraint that the net flux vanishes. For comparison to the computational model, this was imposed by again specifying *L*_1_ = *L*^*^ and *L*_2_ = *L*_*trunc*_ along with the additional constraint 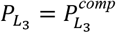 at length *L*_3_ = *L*_2_ − 1, chosen for convenience. We found good agreement between the resulting numerical solutions and our computational model results. Additionally, solutions to this equation accurately describe the parameter regimes that are well approximated by **Equations 10** and **11**, as the latter represent nested dynamics characterized by **Equation 12** that discard negligible contributions. Thus, **Equation 12** has a regime of validity that extends across the entire set of parameter combinations that result in stable distributions Δτ > 0. As a corollary, the accuracy of this approximation to the full steady state dynamics across the space of computational model results indicates that the effects of repeat fusion remain negligible throughout. However, this statement is only applicable to the long repeat dynamics for L>10; the effects of repeat fusion are everywhere relevant for short repeats, which, in part, shape the geometric distribution at steady state.

Details on the derivation, relevant approximations, dynamical regimes, and comparison between numerical and computational model results are provided in depth in **Supplementary Note SN1**.

## Supporting information

Supplementary Data File SF1: Mammalian repeat length counts

Supplementary Data File SF2: Rate data

## Data Availability

The datasets analyzed during the current study are freely available from the NCBI (https://www.ncbi.nlm.nih.gov/datasets/genome/), the UCSC Genome Browser (https://genome.ucsc.edu), and other studies as cited. Instructions for accessing specific datasets are further detailed in the code repository (see Code Availability). Repeat length distributions for mammalian genomes analyzed in this study are available in Supplementary Data File SF1. Repeat length instability rates calculated in this study are available in Supplementary Data File SF2.

## Code availability

The code to perform the analysis in the current study is available in a Github repository (https://github.com/ryanmcggg/repeat_distributions). For software/packages (with version numbers), please visit the Github repository.

## Author contributions

RM conceived the study and performed bioinformatic analyses. RM and DB jointly designed and implemented the computational model and prepared the manuscript. DB and SS constructed the analytic model and mathematical analyses. RM, DB and SS devised the statistical inference procedure. SM and SS supervised the project, discussed results and prepared the manuscript.

## Acknowledgements

Work in the SS lab is supported by grants from NIGMS R35GM127131, NIMH R01MH101244 and NHGRI U01HG012009. Work in the SM lab is supported by grants from NIGMS R35GM130322 and NSF-BSF 2153071. We thank Alexey Kondrashov and Alisa Lyskova for helpful discussions at the early stages of the project.

---

**Fig. S1.**
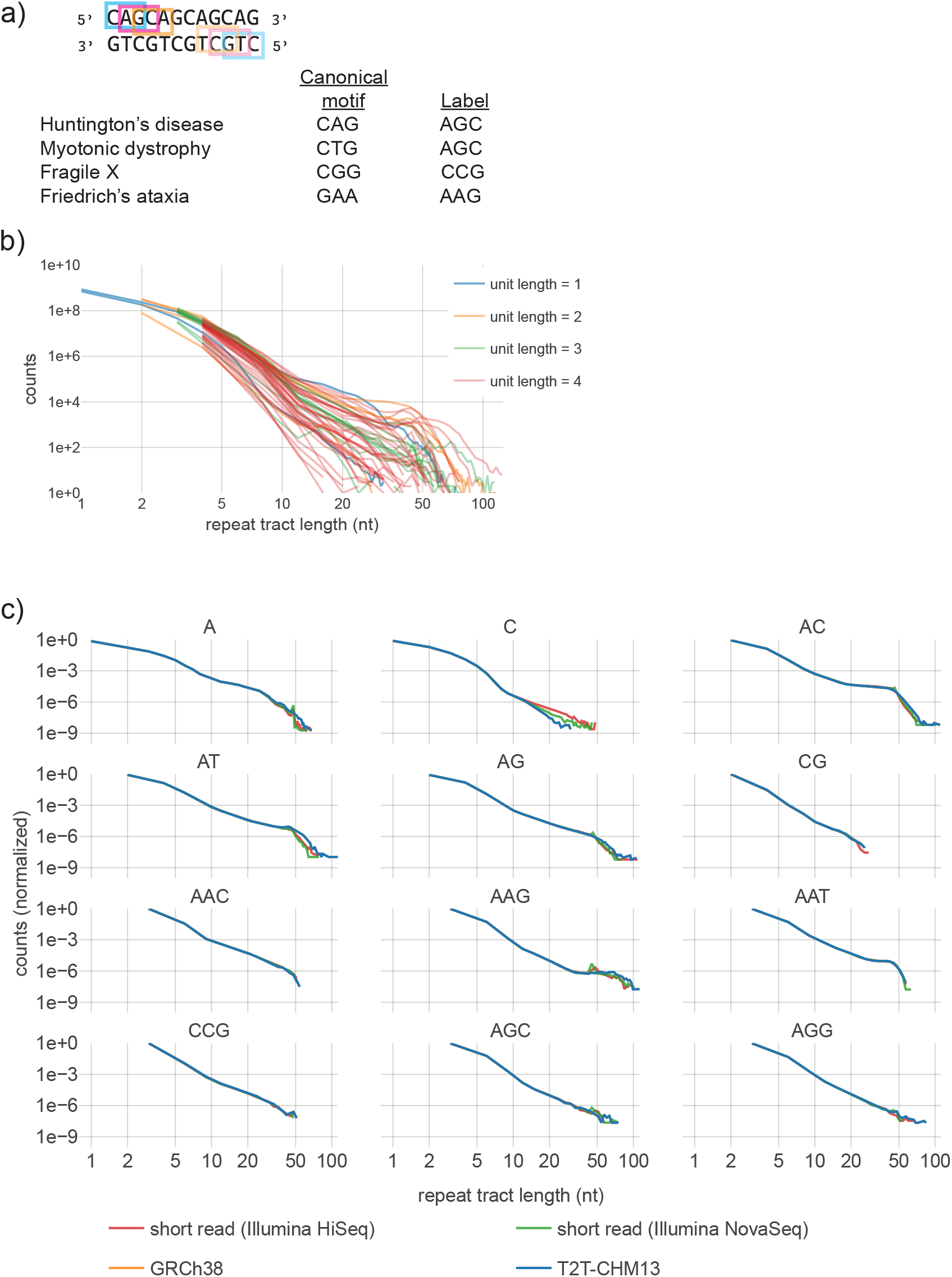
Motif labeling and distributions of repeat tract lengths (DRLs). **a)** Motif labels describe estimates pooled across equivalent cyclical permutations and their reverse complements (example shown for ‘AGC’), using the alphabetically-first label. Labels of well-studied repeat expansion disorders are provided as examples. **b)** DRLs in human T2T genome for all distinct motifs with unit lengths 1–4 nt. Motifs of the same unit length are shown in the same color. DRLs vary by motif but share similar qualitative features. **c)** DRLs in four distinct human genome assemblies with different sequencing technologies. Normalization is required to account for differences in total sequence length (see **Fig. S2**). T2T-CHM13 was assembled using multiple long read technologies, while other assembles employed shorter read technologies. Results shown for all motifs of unit length 1–3 (labeling described in panel a). Read length does not appear to affect the accurate counting of repeats of sub-disease lengths (with the exception of long mono-C repeats in one example short-read genome).

**Fig. S2.**
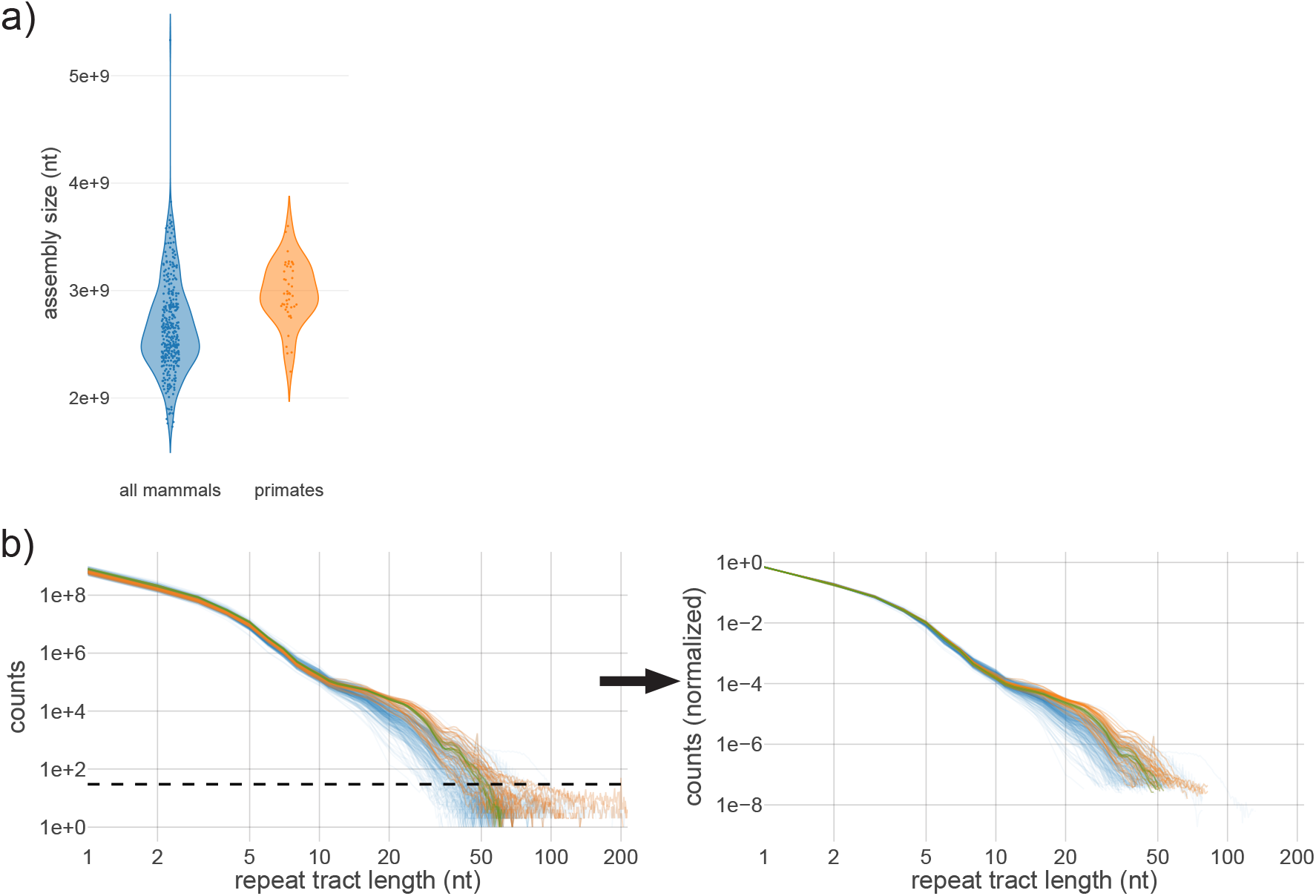
DRL normalization for comparison across genomes of varying lengths. **a)** Differing assembly size for ∼300 mammalian genomes (i.e., total sequence length in each FASTA file, rather than complete *in vivo* genome size). Comparisons between genomes necessarily require normalization due to expected differences in total number of repeats. **b)** Example of DRL normalization for mono-A repeats (used to produce **Fig. 1b**). Left panel shows dashed line at 30 counts; DRLs were truncated to remove highly stochastic bins below this threshold prior to normalization (right panel; see **Methods**).

**Fig. S3.**
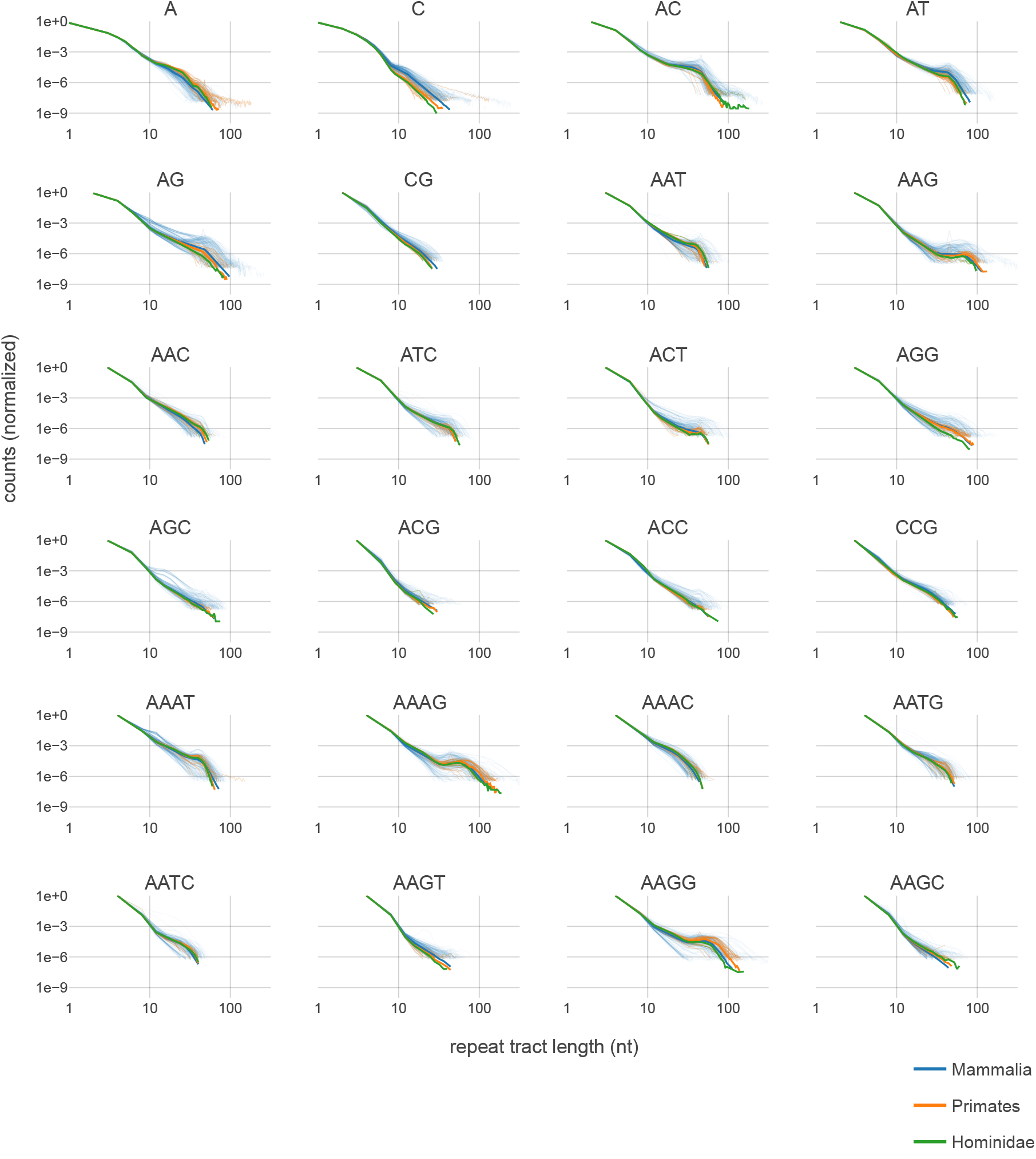
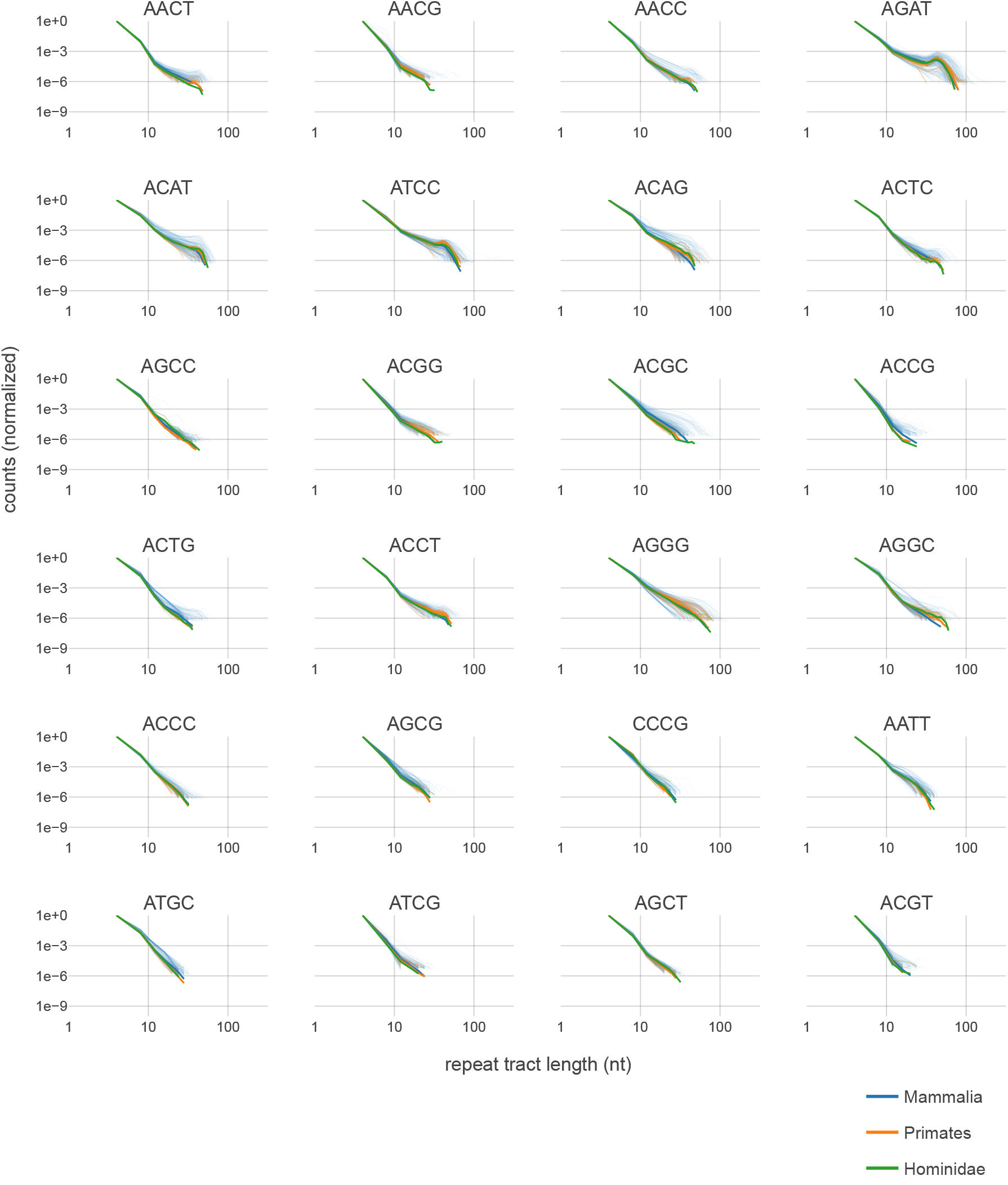
Distributions of repeat tract length per motif across phylogenies. Normalized DRLs for all motifs of unit length 1–4 in mammals, primates and hominids. Motif labels include equivalent cyclical permutations and their reverse complements (see **Fig. S1a**). Counts are necessarily normalized to account for different genome lengths (see **Fig. S2, Methods**). Solid line indicates median values per length bin for each phylogeny. Median calculations within phylogenies are inclusive (e.g., primates are included as a subset of mammals). Thin transparent lines show individual species; DRLs are truncated at the shortest bin containing 30 counts prior to normalization (see **Fig. S2**). Overlapping medians between primates and hominids suggests long-term stability of the DRLs, while individual mammalian species display variability.

**Fig. S4.**
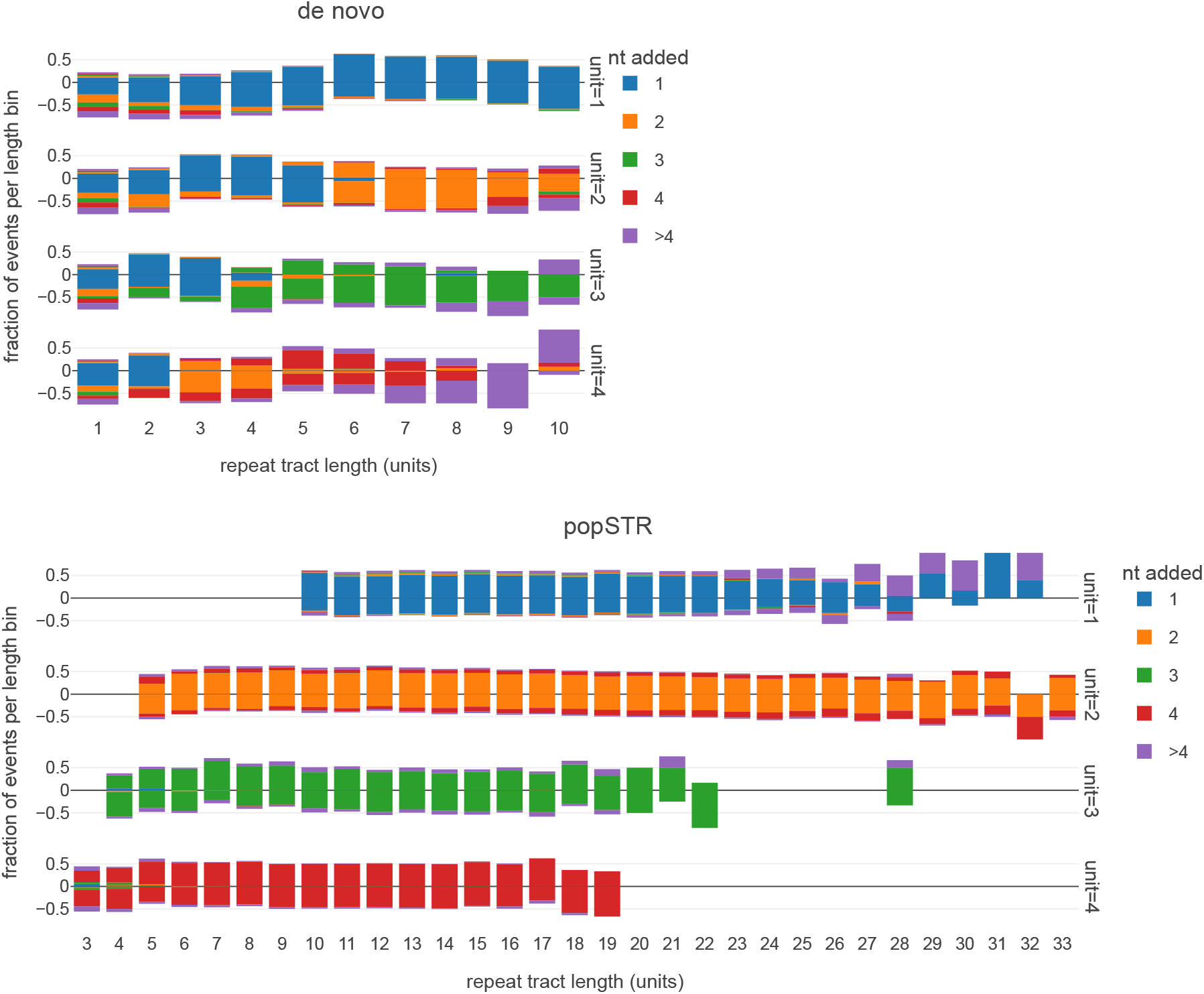
Nucleotides gained or lost per indel event. De novo indels measured from two different data sources: pooled trio sequences (**top**) and popSTR estimates **(bottom**). Row labels indicate the motif unit length. X-axis displays the tract length of the repeat in number of units. Y-axis displays the fraction of events per length bin, according to the number of nucleotides gained or lost per event (indicated by color). Y-axis sign indicates insertion (positive) or deletion (negative); overall shift up or down indicates bias. Most events involve the gain or loss of a single complete unit, independent of total repeat length; both partial-unit and multiple-unit changes are less frequent. A length threshold is apparent (see **Fig. 3a**), above which single unit indels characterize repeat instability. Note that indel calling in the de novo dataset loses accuracy at longer tract lengths; likewise, the popSTR database is not sufficiently populated below 10 nt and above ∼30 nt.

**Fig. S5.**
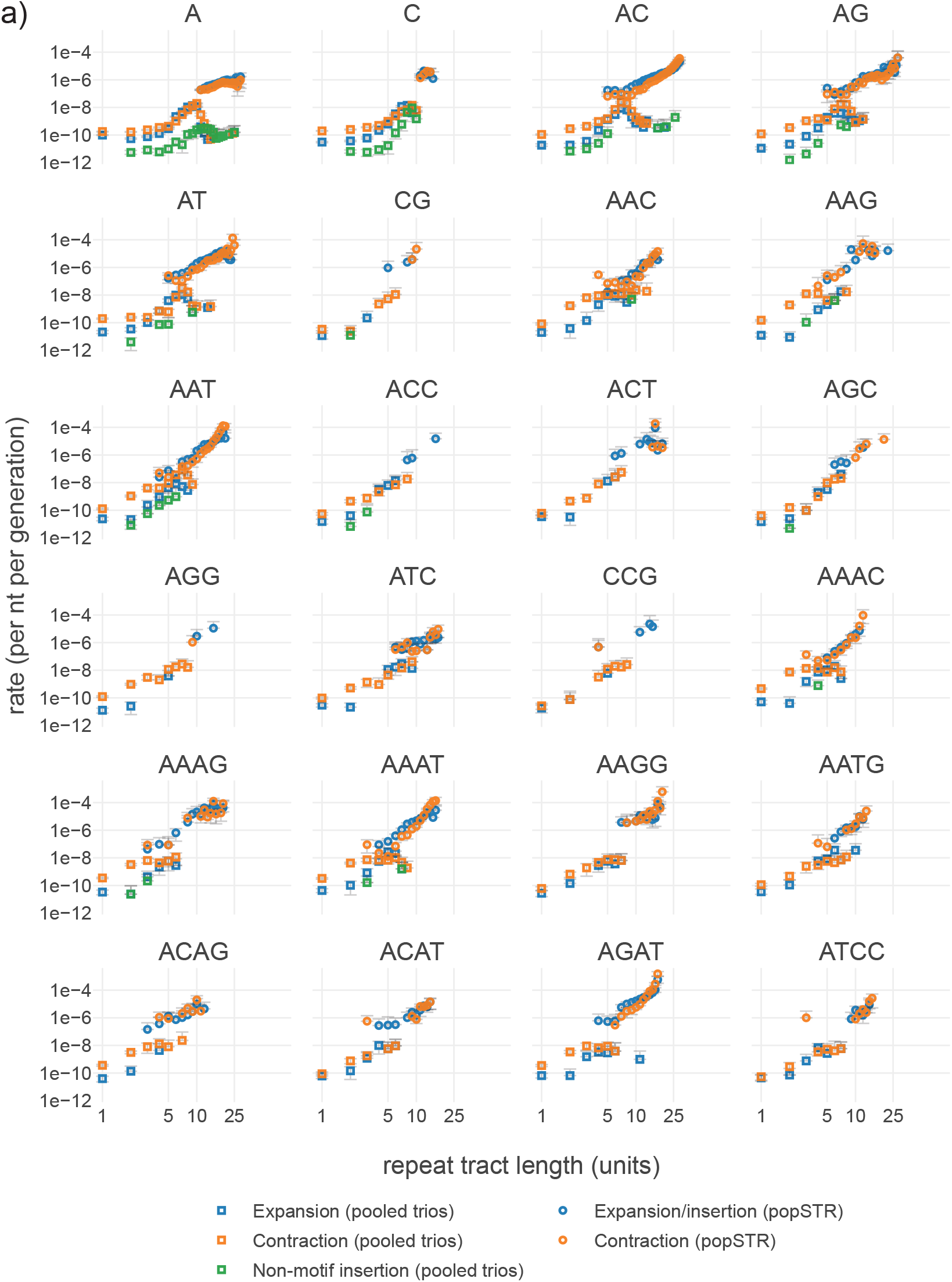
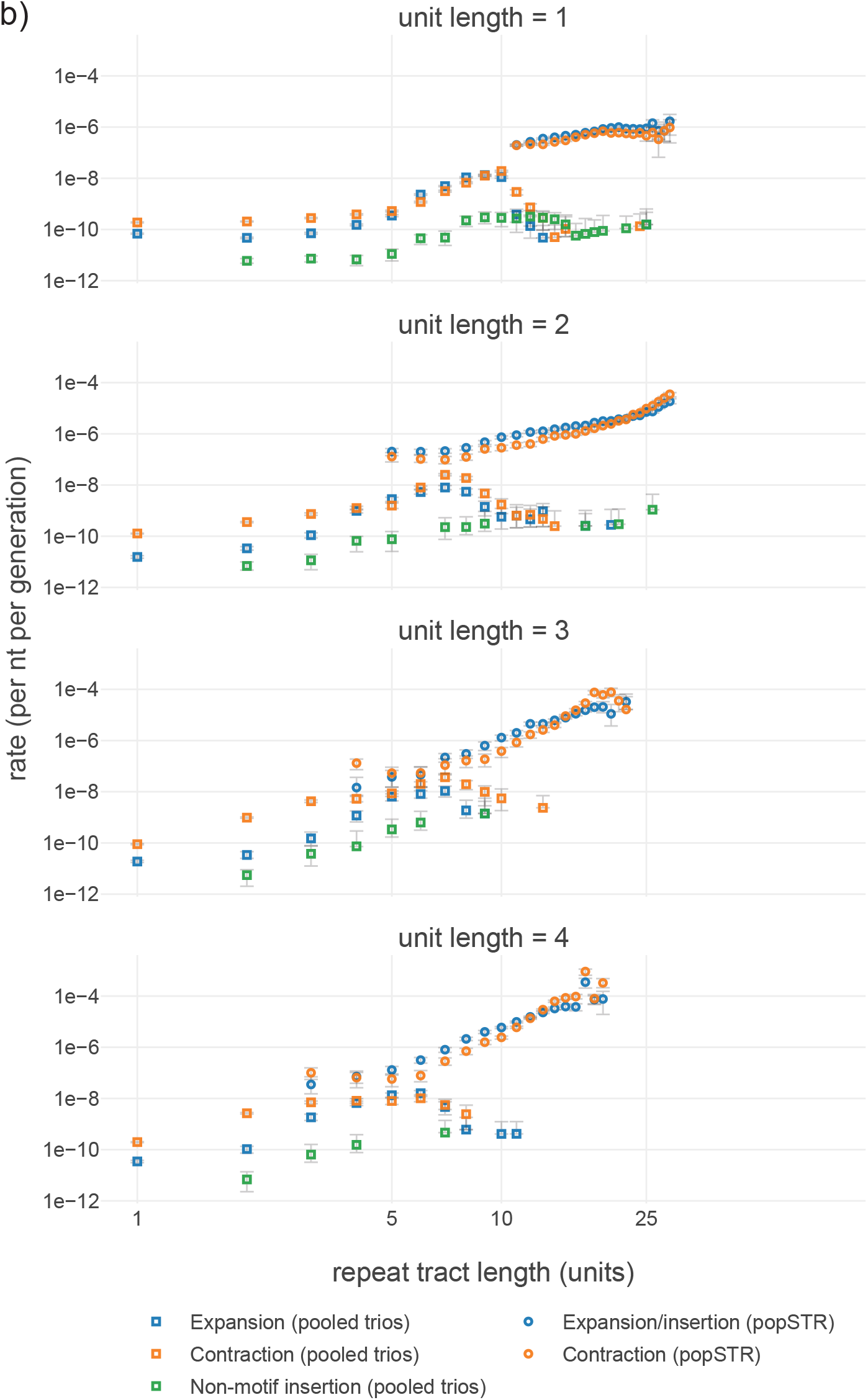
Instability rate estimates. **a)** Rate estimates from pooled-trio and popSTR datasets for expansion, contraction and non-motif insertions for all repeat motifs of unit lengths 1–4 nt. Statistical error bars show 95% CI assuming Poisson mutation counts. Motif labels include equivalent cyclical permutations and their reverse complements (see **Fig. S1a**). Complete estimates shown (no points omitted due to systematic errors and/or low sequencing quality). **b)** Rate estimates from de novo and popSTR datasets for expansion, contraction and non-motif insertions, pooled by motif unit length. Statistical error bars show 95% CI assuming Poisson mutation counts. No points omitted due to systematic errors and/or low sequencing quality.

**Fig. S6.**
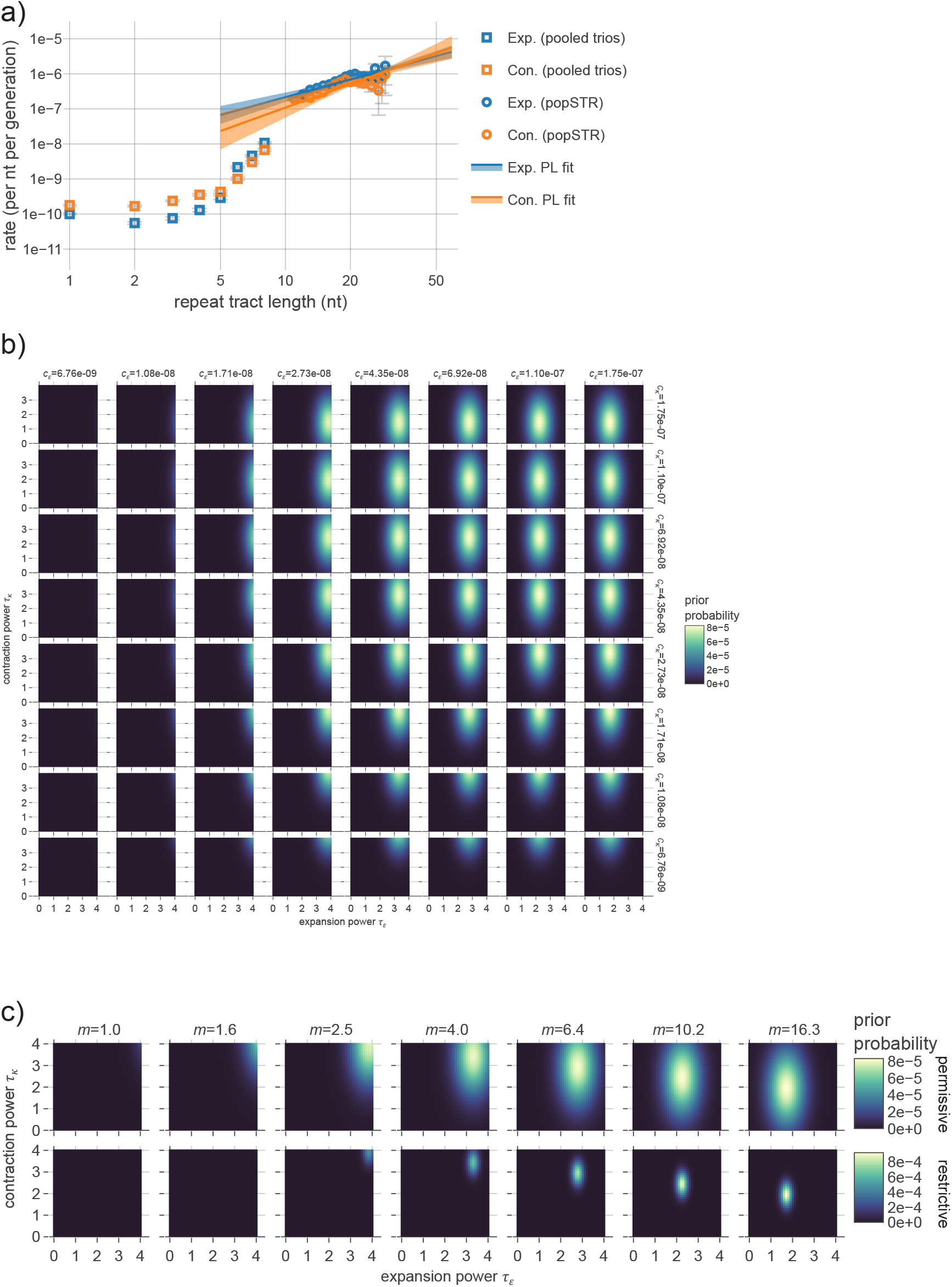
Construction of informative priors. **a)** Rate estimates from pooled-trio and popSTR datasets for expansion, contraction and non-motif insertions for mono-A repeats, as shown in **Fig. 3b**. Lines show best-fit to popSTR expansion (plus insertion) and contraction rate estimates under a linear model in log-log space (i.e. power law; see **Methods**). Transparency approximates 95% confidence region for each linear fit. **b)** Bayesian informative prior for the four-parameter de-coupled power law model (parameterization in **Table 1**). Parameters (c_*ε*_, c_*κ*_, τ_*ε*_, τ_*κ*_ ) shown as (columns, rows, x-axes, y-axes). Prior was constructed as a multivariate Gaussian probability distribution with mean parameters corresponding to best fit in panel a; co- variance matrix estimated from confidence band after artificially inflating variances and covariances (1000x for permissive; 100x for restrictive, not shown). See **Methods** for further detail. **c)** Informative priors for three-parameter multiplier-coupled power law model. Due to nested parameterizations (see **Fig. S16**), these priors were defined as appropriately re-normalized slices of the four dimensional prior (permissive - top; restrictive - bottom).

**Fig. S7.**
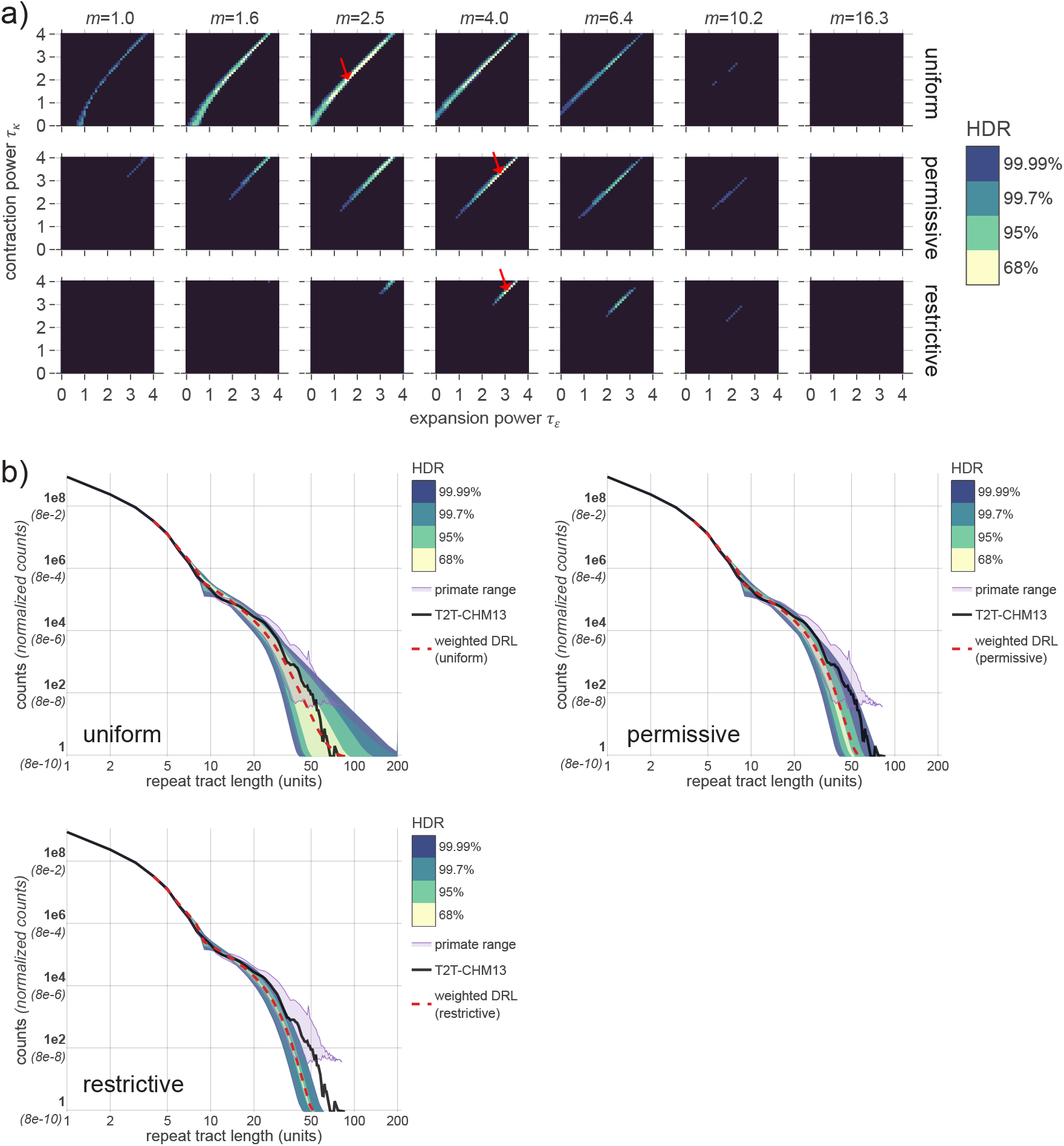
Inference results for three-parameter multiplier-coupled power-law model of repeat instability rates. **a)** Plots display Bayesian posterior probabilities after applying uninformative (top row), permissive (middle row) and restrictive (bottom row) informative priors (bottom; see **Fig. S6**). Length-dependent instability rates defined by parameters (*m*, τ_*ε*_, τ_*κ*_ ) shown as columns, x-axes and y-axes, respectively. Color bar indicates highest density range (HDR) of the posterior for specified total probabilities; black region sums to 0.01% of the probability. Red arrows show maximum posterior for each prior. **b)** Comparison of empirical DRLs to various inference results under specified priors. Counts for all DRLs are necessarily normalized for comparison (see **Methods)**; y-axis indicates normalized fractions (parentheses) and counts rescaled to match the number of repeats in the T2T genome (bold labels, black curve). Dashed line represents posterior-weighted DRL (an average of all DRLs weighted by the posterior probability for each parameter combination; see **Methods**). Colored ranges represent the minimum and maximum counts at each length bin across all parameters in the specified HDR (color corresponding to panel a). Purple region shows the min-max range generated from 36 primate genomes (after removing the two most-diverged DRLs and appropriately normalizing; see **Fig. S2, Methods**).

**Fig. S8.**
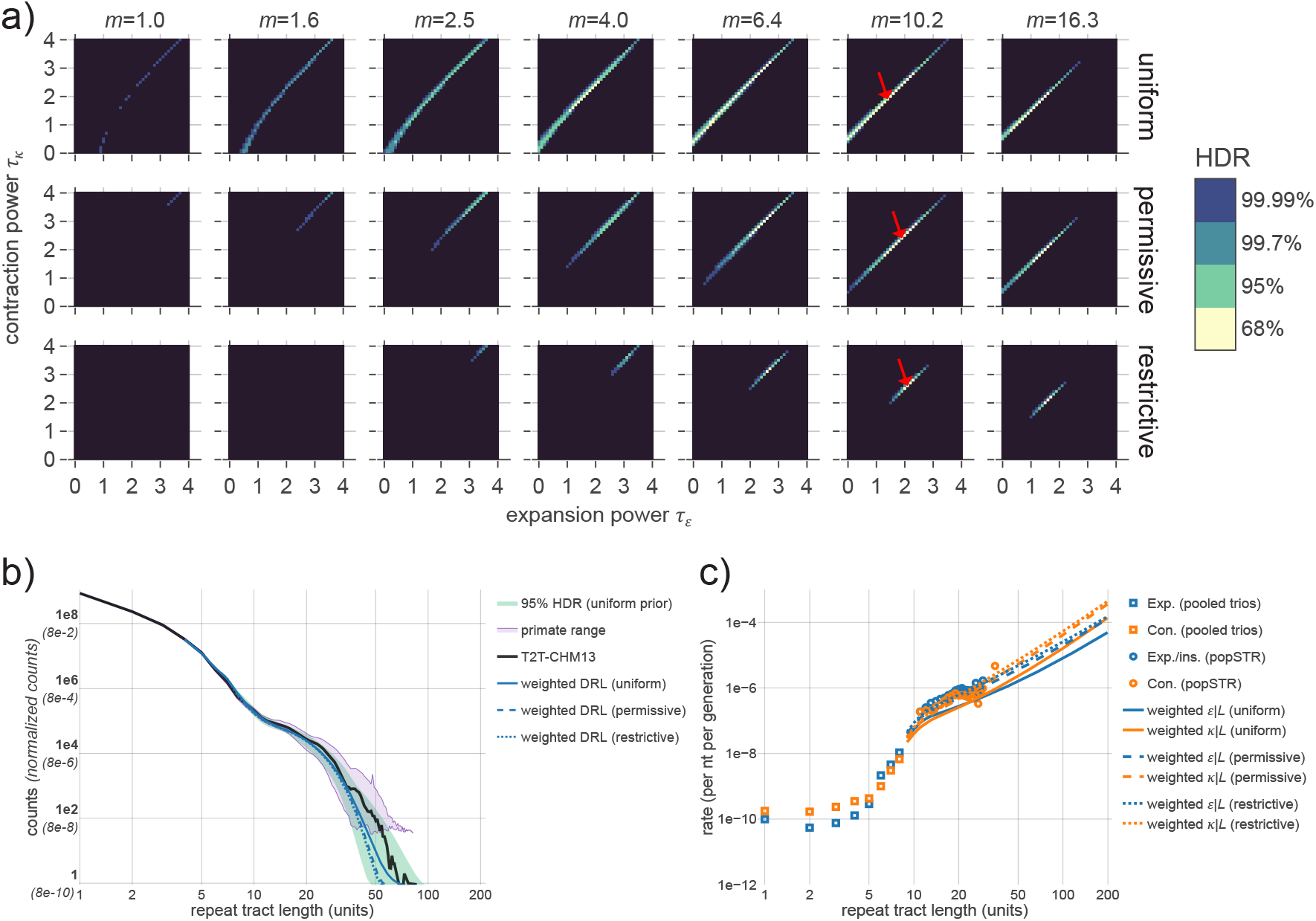
Posterior probability distributions for interpolated length-dependent rates. **a)** Inference results for three-parameter multiplier-based power-law model of repeat instability rates after interpolating between lengths 8 and 13 nt (see **Methods**). Plots display Bayesian posterior probabilities after applying uninformative (top row), permissive (middle row) and restrictive (bottom row) informative priors (see **Fig. S6**). Each coordinate represents a distinct set of length-dependent instability rates defined by parameters (*m*, τ_*ε*_, τ_*κ*_ ) shown in columns, x-axes and y-axes, respectively; τ_*ε*_ and τ_*κ*_ determine the power laws for expansion and contraction, respectively, and *m* is a multiplicative jump at L=9 (parameterization defined in **Table 1**). Color indicates highest density range (HDR) of the posterior for specified total probabilities; black region sums to 0.01% of the probability. Red arrows show maximum posterior for each prior. Interpolation shifts posterior density to higher *m* values. **b)** Comparison of empirical DRLs to inference results for each prior. Counts for all DRLs are necessarily normalized for comparison (see **Methods)**; y-axis indicates normalized fractions (parentheses) and counts rescaled to match the number of repeats in the T2T genome (bold labels). Blue lines represent posterior-weighted DRLs for informative and uninformative priors; modeled DRLs are largely consistent with the empirical T2T DRL (black). Range shown in green represents the minimum and maximum counts at each length bin across all parameters in the 95% HDR under the uniform prior. Purple region shows the min-max range generated from 36 primate genomes (after removing the two most-diverged DRLs and appropriately normalizing; see **Fig. S2, Methods**). Interpolation removes discontinuity in modeled DRLs between *L*=8 and 9. **c)** Posterior-weighted repeat instability rates. Tract length dependencies of expansion (blue) and contraction rates (orange) shown for specified priors. For comparison, empirical estimates are shown for pooled trios (squares; directly incorporated in model) and popSTR data (circles; used to construct informative priors). Informative priors show greater consistency with popSTR-estimated rates after interpolation, while the posterior-weighted DRLs (panel b) remain consistent with the T2T genome.

**Fig. S9.**
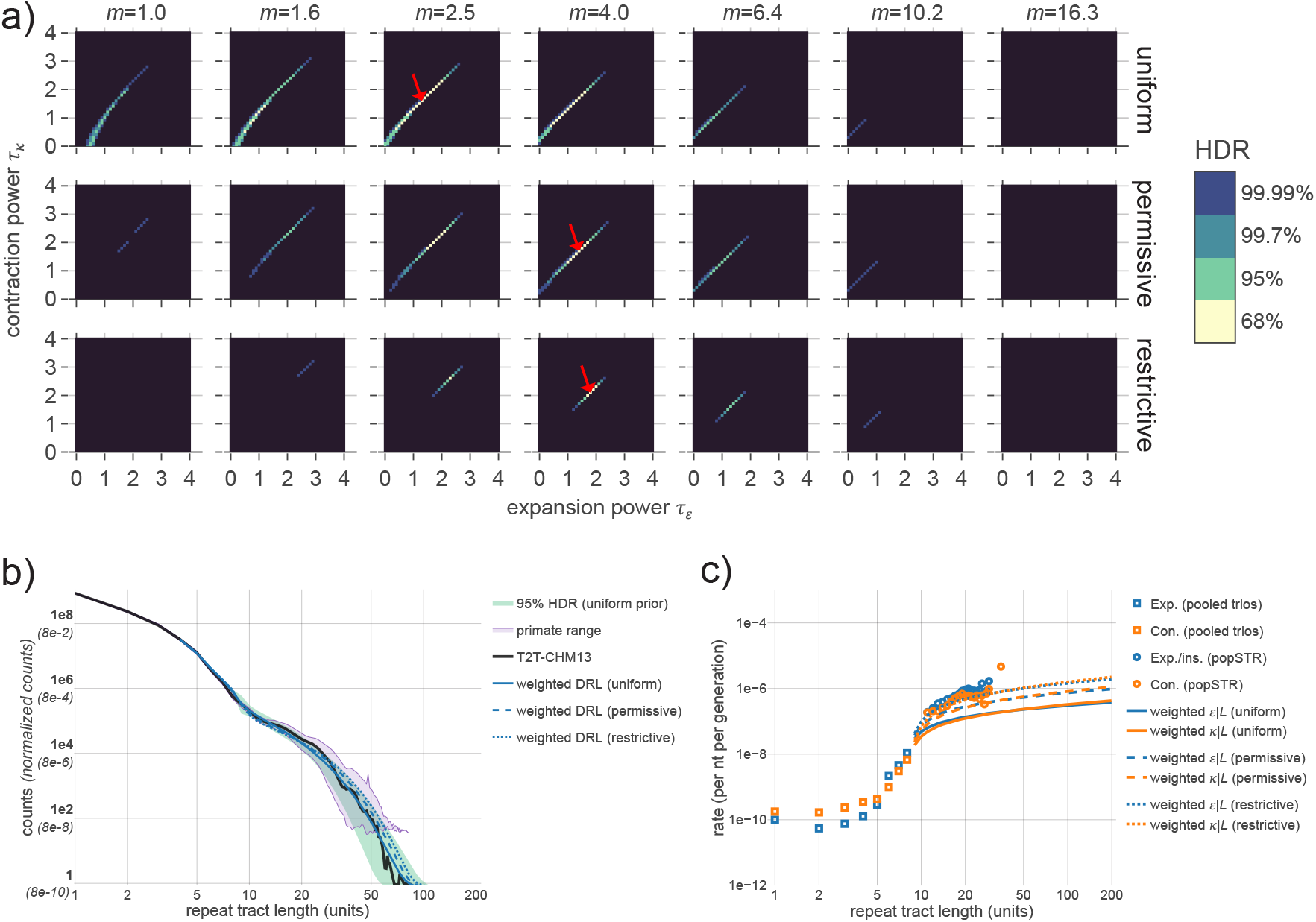
Posterior probability distributions under logarithm-based parameterization. **a)** Inference results for three-parameter multiplier-based model of repeat instability rates using a log-based functional form (see **Methods**). Plots display Bayesian posterior probabilities after applying uninformative (top row), permissive (middle row) and restrictive (bottom row) informative priors (see **Fig. S6**). The parameters (*m*, τ_*ε*_, τ_*κ*_ ) are defined analogously to the power-law model (see **Table 1** for parameterization). Color indicates highest density range (HDR) of the posterior for specified total probabilities; black region sums to 0.01% of the probability. Red arrows show maximum posterior for each prior. Posterior distribution is highly similar to that for the power-law parameterization, indicating that specific functional forms should not be over-interpreted. **b)** Comparison of empirical DRLs to inference results for each prior. Counts for all DRLs are necessarily normalized for comparison (see **Methods)**; y-axis indicates normalized fractions (parentheses) and counts rescaled to match the number of repeats in the T2T genome (bold labels). Blue lines represent posterior-weighted DRLs for informative and uninformative priors; modeled DRLs are largely consistent with the empirical T2T DRL (black). Range shown in green represents the minimum and maximum counts at each length bin across all parameters in the 95% HDR under the uniform prior. Purple region shows the min-max range generated from 36 primate genomes (after removing the two most-diverged DRLs and appropriately normalizing; see **Fig. S2, Methods**). **c)** Posterior-weighted repeat instability rates. Tract length dependencies of expansion (blue) and contraction rates (orange) shown for specified priors. For comparison, empirical estimates are shown for pooled trios (squares; directly incorporated in model) and popSTR data (circles; used to construct informative priors). Like the power-law parameterization, this parameterization allows simultaneous consistency with popSTR estimates and empirical DRLs.

**Fig. S10.**
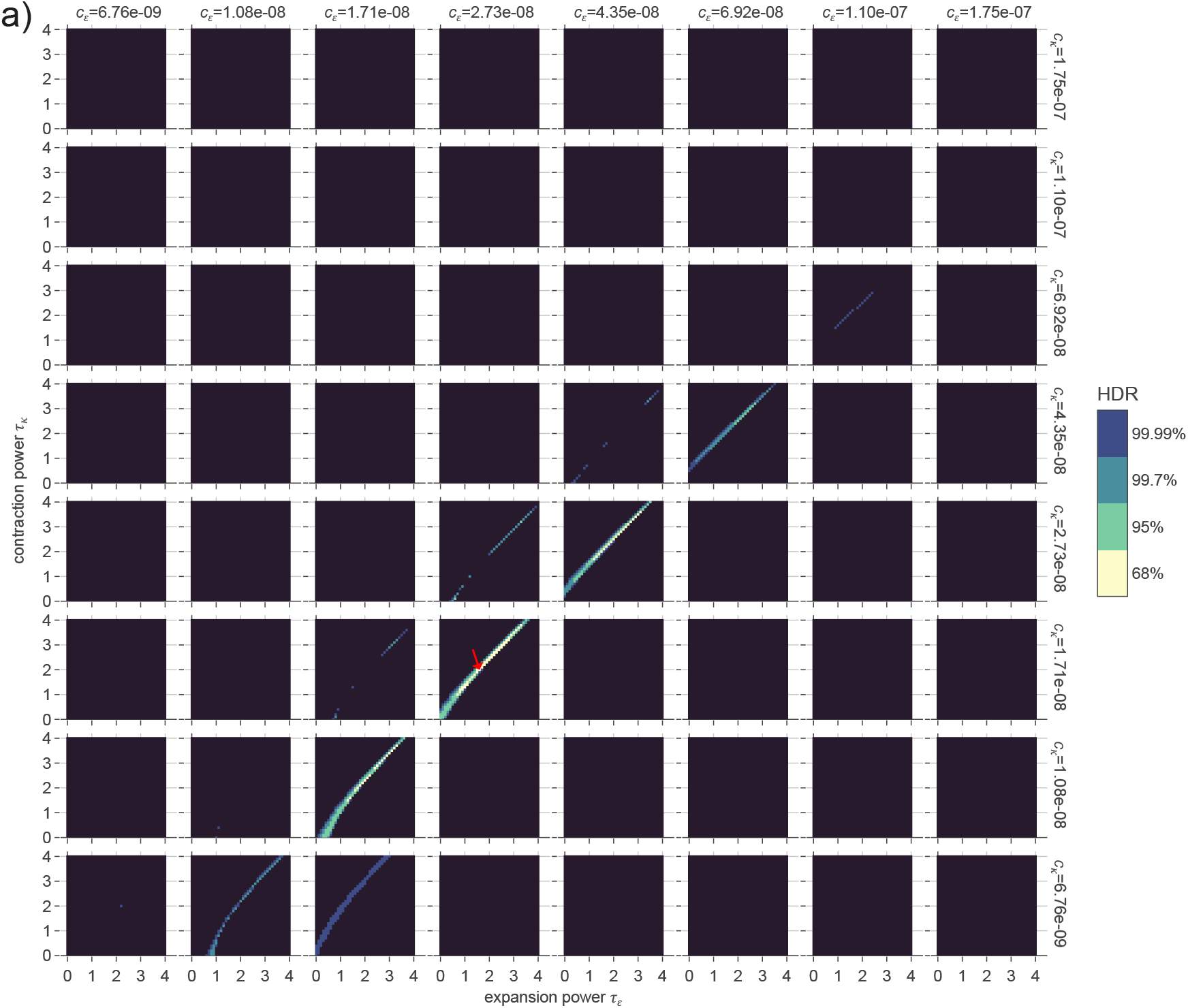
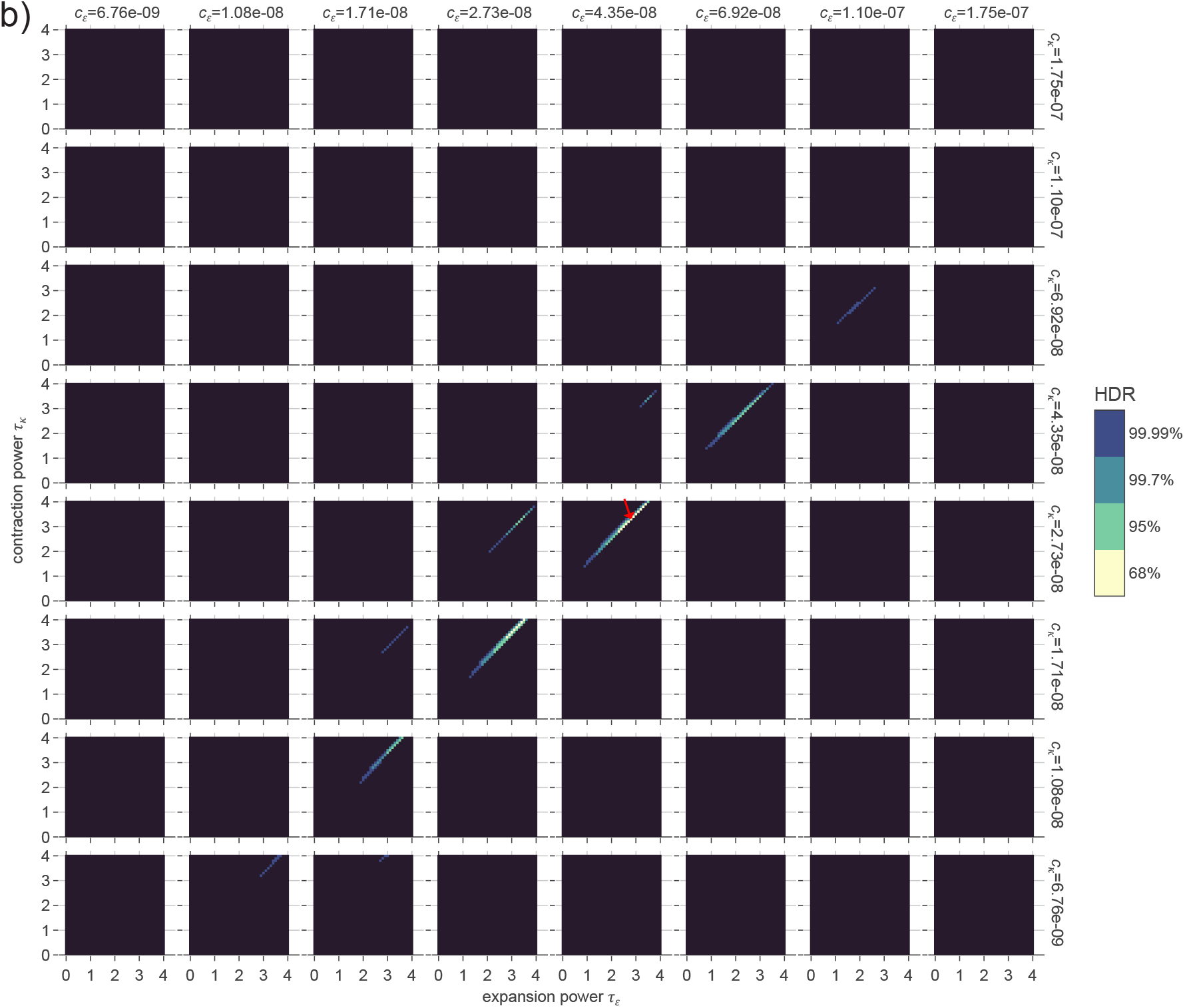
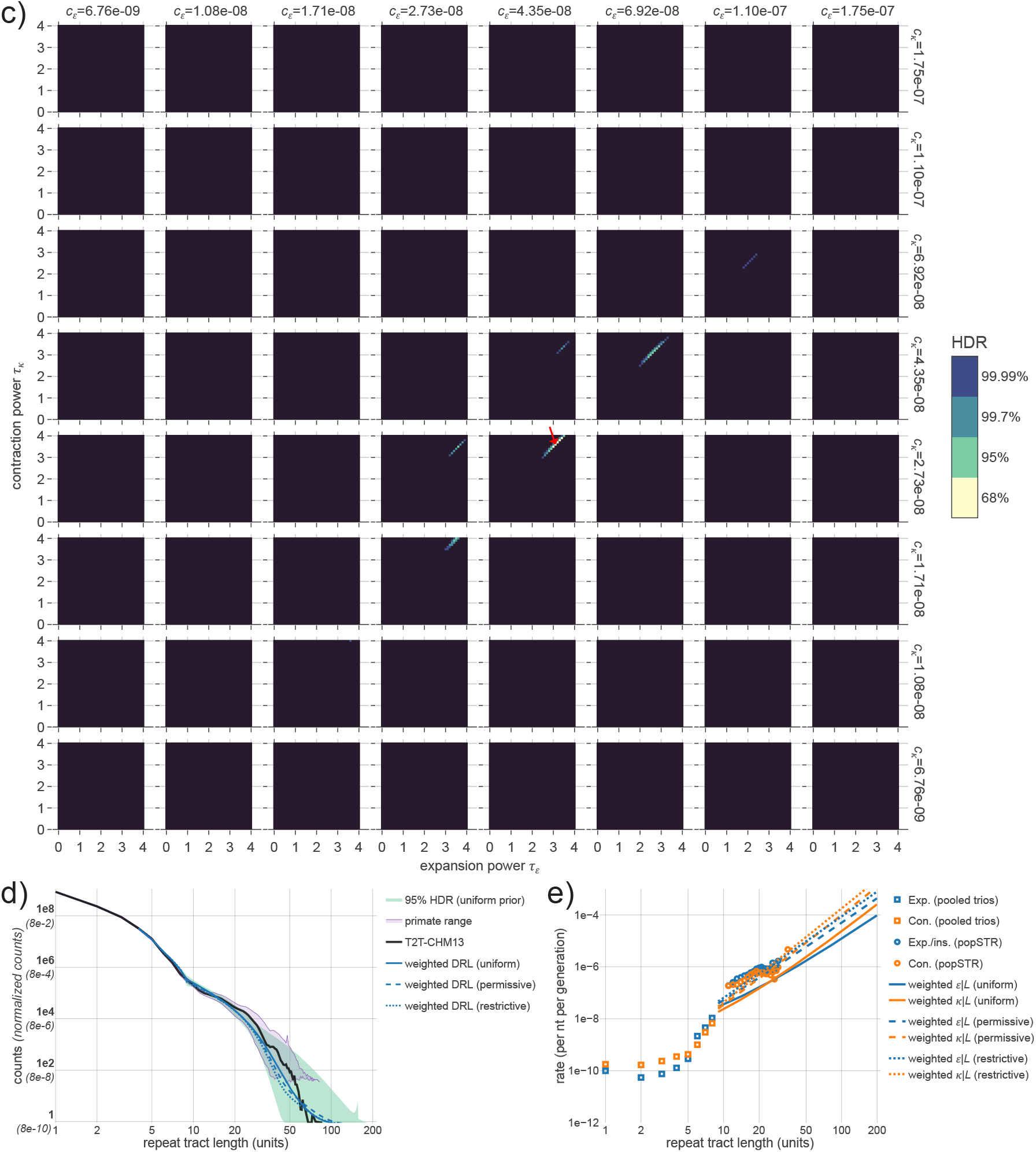
Inference results for four-parameter decoupled power-law model of repeat instability rates. **a)** Bayesian posterior probabilities for the uninformative prior. Each coordinate represents a distinct set of length-dependent instability rates defined by parameters (c_*ε*_, c_*κ*_, τ_*ε*_, τ_*κ*_ ) shown in columns, rows, x-axes and y-axes, respectively; τ values determine power law exponents, and *c* values determine the rates at *L*=9 (parameterization defined in **Table 1**). Color indicates highest density range (HDR) of the posterior for specified total probabilities; black region sums to 0.01% of the probability. Red arrows show maximum posterior. **b)** Bayesian posterior probabilities for the permissive informative prior. **c)** Bayesian posterior probabilities for the restrictive informative prior. For all three priors, most of the density lies within the nested parameterization corresponding to the three-parameter multiplier-coupled model. **d)** Comparison of empirical DRLs to inference results for each prior. Counts for all DRLs are necessarily normalized for comparison (see **Methods)**; y-axis indicates normalized fractions (parentheses) and counts rescaled to match the number of repeats in the T2T genome (bold labels). Blue lines represent posterior-weighted DRLs for informative and uninformative priors; modeled DRLs are largely consistent with the empirical T2T DRL (black). Range shown in green represents the minimum and maximum counts at each length bin across all parameters in the 95% HDR under the uniform prior. Purple region shows the min-max range generated from 36 primate genomes (after removing the two most-diverged DRLs and appropriately normalizing; see **Fig. S2, Methods**). **e)** Posterior-weighted repeat instability rates. Tract length dependencies of expansion (blue) and contraction rates (orange) shown for specified priors. For comparison, empirical estimates are shown for pooled trios (squares; directly incorporated in model) and popSTR data (circles; used to construct informative priors). This parameterization provides a qualitatively similar set of observations to the three-parameter multiplier-coupled model (e.g., similar Bayes factors, posterior-weighted DRLs and instability rates are consistent with empirical estimates and display asymptotic contraction bias), but is less dependent on empirical rate estimates at *L*=8.

**Fig. S11.**
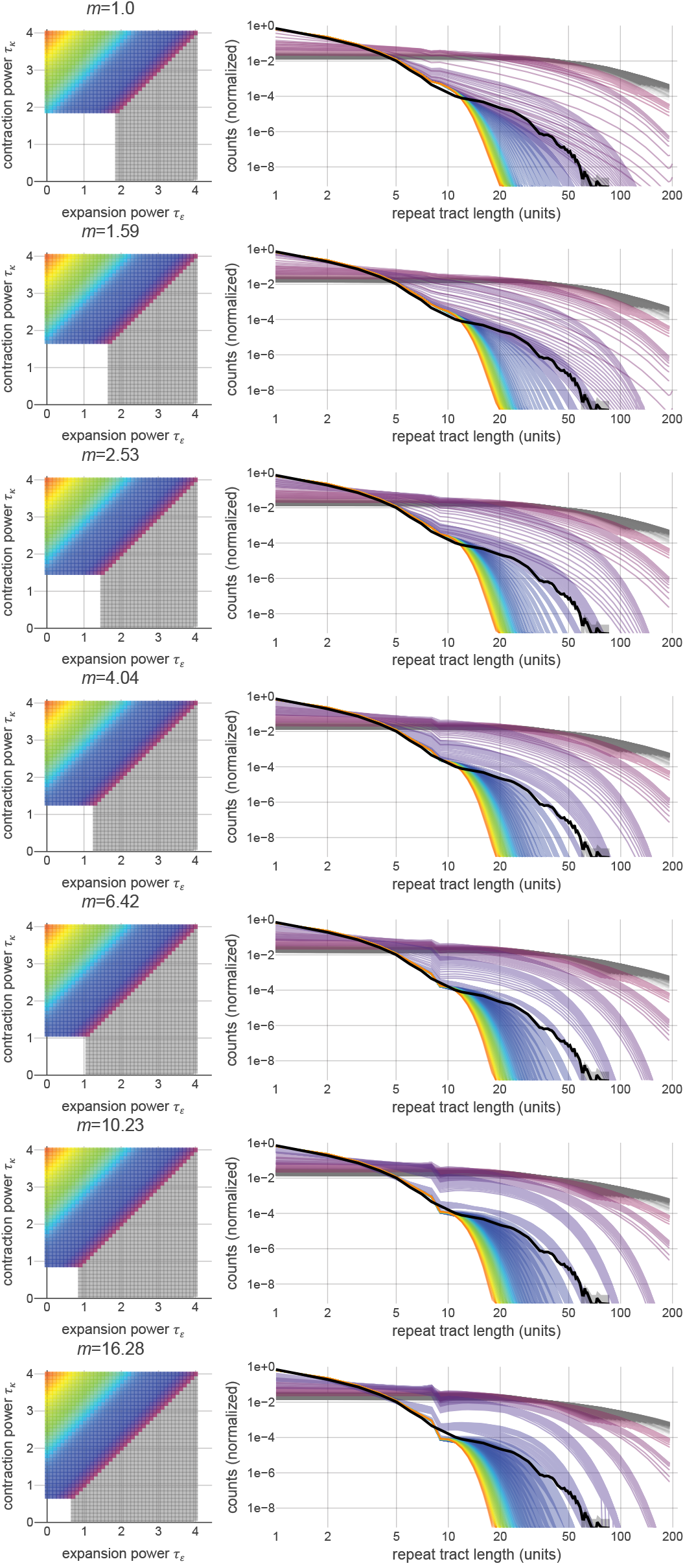
Late-time DRLs from computational model across parameter space. Results from three-parameter multiplier-coupled power law model. **(Left)** Grid of (τ_*ε*_, τ_*κ*_ ) parameter values for various multipliers *m*. For clarity, lower left region omitted due to low rates insufficient to equilibrate in the allotted time. Lines of constant Δτ=τ_*κ*_ -τ_*ε*_ are displayed in the same color. Red corresponds to large values of Δτ, purple corresponds to low values of Δτ, and gray represents negative values of Δτ. **(Right)** Plots of DRLs at the final time point, one for each parameter combination in the grid, using the same color-coding scheme. Black line depicts the empirical T2T DRL. Larger Δτ results in more rapid truncation of the distribution at lower lengths; smaller values of Δτ result in a more extended tail of long repeats. Negative values of Δτ result in unrealistic inflation of long tail; subsequent normalization leads to lower relative counts in short tract length bins. Higher *m* values result in larger discontinuity in the DRL between *L*=8 and 9 units.

**Fig. S12.**
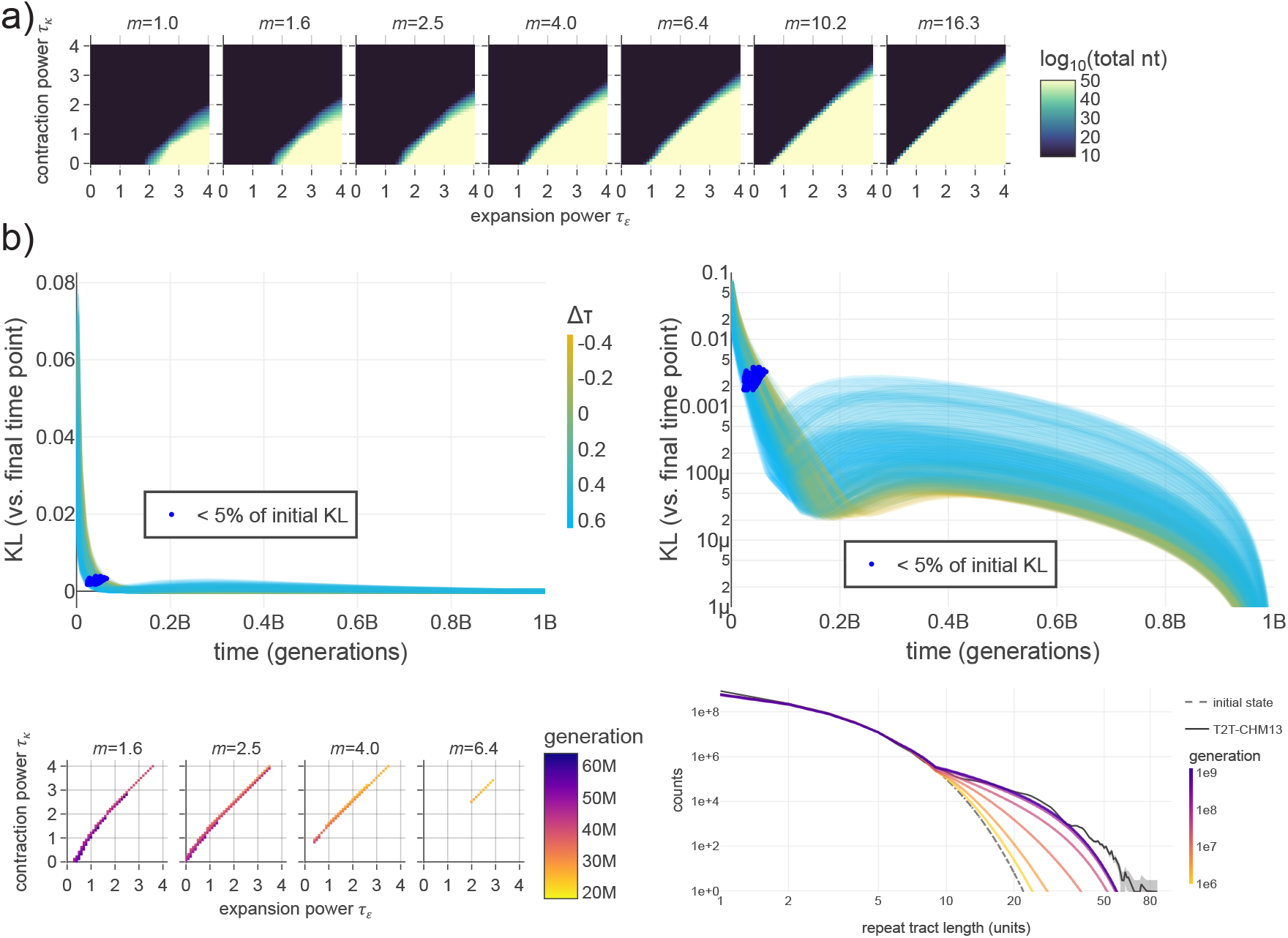
Constraints from genome size and equilibration time. **a)** Plot of total number of repeat bases in genome at final time point of computational model. Displayed as a heatmap across parameter space (*m*, τ_*ε*_, τ_*κ*_ ) for the three-parameter multiplier-coupled power-law model. Color specifies log_10_ of the sum of lengths of all repeats in the DRL. Color range is truncated at 10^50^. Explosive genome growth occurs for heavily expansion-biased parameter combinations. **b)** Equilibration time for computational model. **Top left)** Each line represents one parameter combination from the 95% HDR in the three-parameter multiplier-coupled model. Computational model initializes the DRL with the geometric distribution expected under substitutions alone. X-axis displays time in number of generations (under constant time rescaling). Y-axis displays KL divergence from the DRL at the final time point for each parameter combination (such that the KL divergence approaches 0 in finite time). Color scale indicates values of Δτ=τ_*ε*_ -τ_*κ*_ . **Top right)** Same as in top left panel, with log-scaled y-axis. For all parameter combinations, a two-stage equilibration process is evident, indicating an initial rapid phase in which the bulk of the extended tail of long repeats is established, followed by a slower phase of finer-scale changes. Blue dots indicate the timepoint at which the KL divergence first drops below 5% of the initial value for each parameter combination, marking the approximate end of the rapid phase. **Bottom left)** The 5% KL divergence timepoint is displayed as a heatmap, showing that higher values of (*m*, τ_*ε*_, τ_*κ*_ ) and Δτ result in faster establishment of the long repeat tail. While not a direct measure of time to achieve steady state, this indicates that, within the timescale of primate evolution, a relatively rapid change in the shape of the DRL is possible. **Bottom right)** Evolution of the DRL over time. Example shown for parameter combination with maximum posterior under uniform prior (*m*, τ_*ε*_, τ_*κ*_ ) = (2.5, 1.6, 2). Colored lines indicate state of the DRL after indicated number of generations (colored in log time). The shape of the DRL is largely established prior to 1e8 generations.

**Fig. S13.**
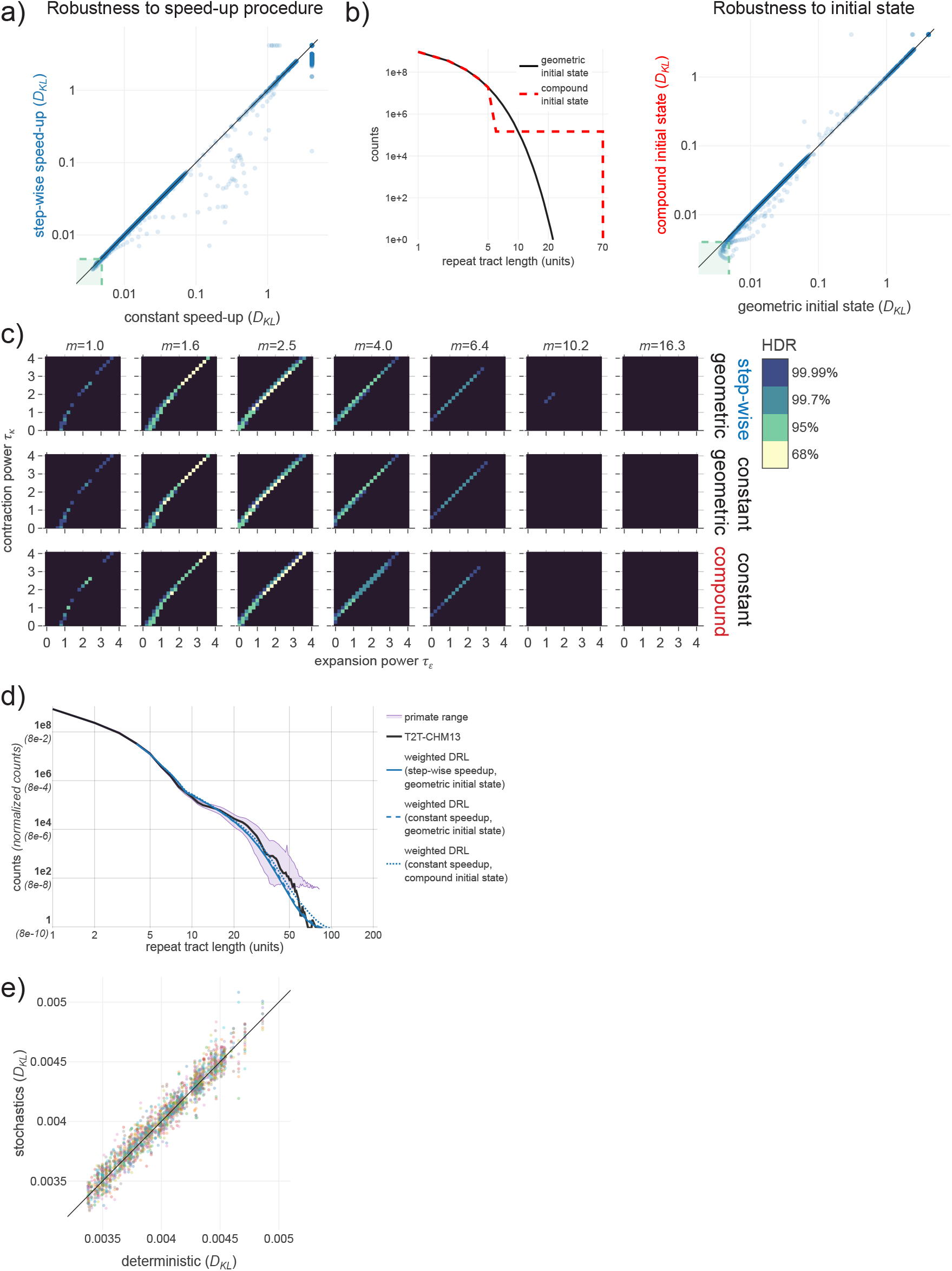
Demonstrations of robustness of inference procedure to simplifying approximations. All comparisons below produced for the three-parameter multiplier-coupled power-law model. *D*_*KL*_ denotes KL divergence relative to the human T2T genome. **a)** To confirm the validity of the computational speed-up procedure, we compared two different run conditions over a sparse grid of parameter combinations (step size = 0.2 for τ_*ε*_ and τ_*κ*_ ). Plot shows correlation between *D*_*KL*_ for final timepoint DRLs, comparing runs with constant speed-up (x-axis) and step-wise speed-up (y-axis) procedures to rescale time. Dashed lines denote the boundary of the 95% HDR. **b)** To confirm that the late-time DRL is independent of the initial condition, we compared runs initialized with two distinct DRLs with the same total weight. Left panel shows two initial DRLs: geometric DRL (used for all other runs; black) and a compound distribution (red) that is geometric with a uniform tail of long repeats. Right panel shows correlation between *D*_*KL*_ for final timepoint DRLs resulting from these initial states (over the same sparse grid of parameter combinations). The vast majority of parameter combinations in panels a and b are highly similar. **c)** Bayesian posterior distributions for the three run conditions included in panels a and b. Color bar indicates highest density range (HDR) of the posterior for specified total probabilities; black region sums to 0.01% of the probability. Correspondence between all three posterior distributions (as well as posterior-weighted DRLs shown in panel **d**) demonstrates robustness of the inference to the computational speed-up procedure and initial condition. **e)** Effect of stochastics on computational model results. We introduced Poisson noise around the number of mutational transitions in each length bin per generation (see **Methods**). 10 stochastic runs are shown for each parameter combination in the 95% HDR (see **Fig. 5a**). Plot shows the limited impact of stochastics on the *D*_*KL*_ summary statistic relative to the 95% HDR (and the empirical range seen in primates).

**Fig. S14.**
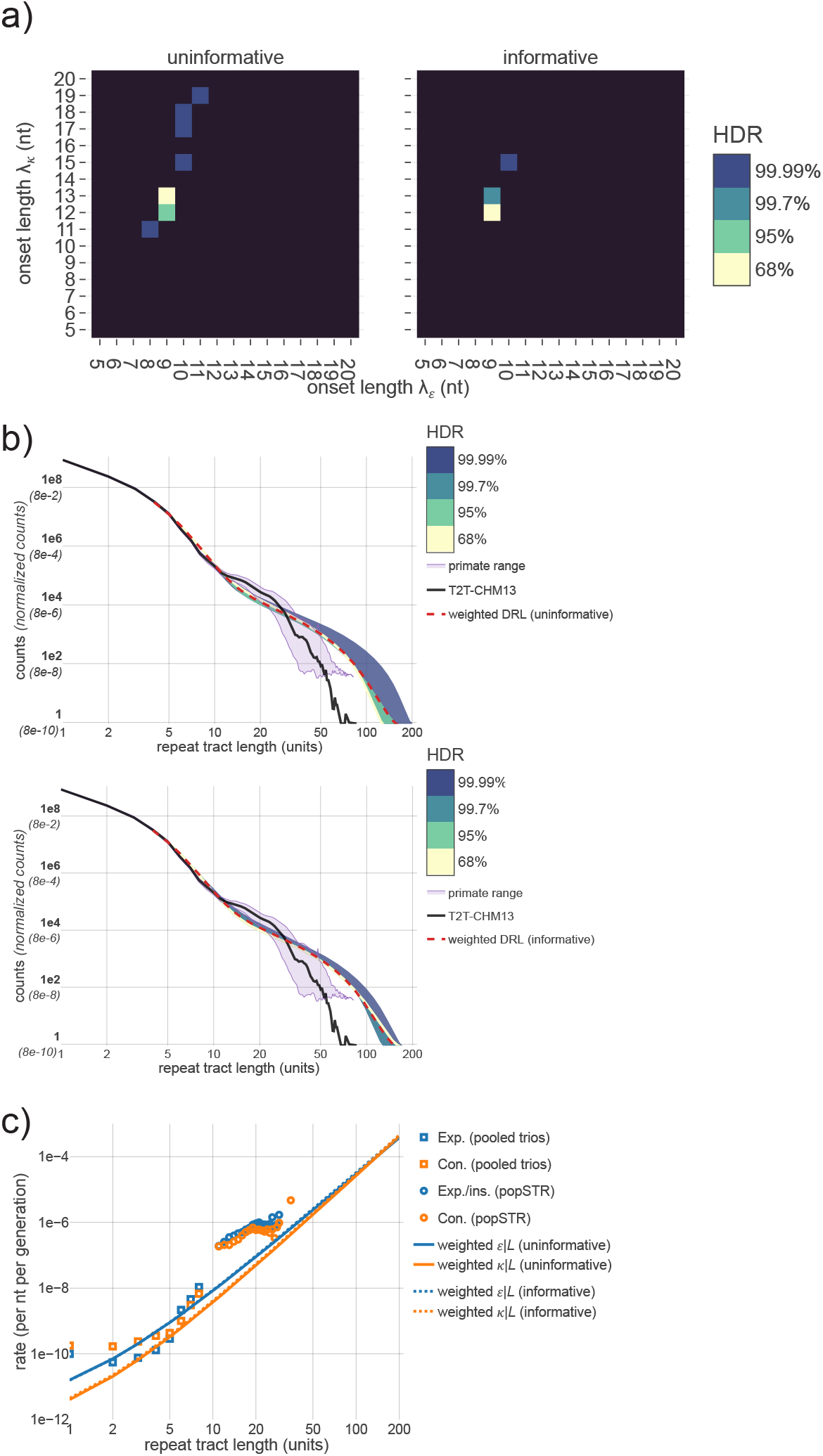
Pure power-law parameterization and the onset length of repeat instability. Bayesian inference results for the pure power-law parameterization (specified in **Table 1**). This model parameterizes expansion and contraction rates at all tract lengths with no reliance on empirical mutation rate estimates; inference results represent information contained within the DRL alone (rate estimates only incorporated in the informative prior). To infer the onset length of repeat instability, the expansion rate was parameterized in terms of λ_*ε*_, the length at which the rate first exceeds the average substitution rate **μ** (lengthening; B>A). Analogously, λ_*κ*_ represents the length at which the contraction rate exceeds the average substitution rate ***ν*** (shortening; A>B). Exponents τ_*ε*_ and τ_*κ*_ parameterize the power-law length dependencies for expansion and contraction, respectively. **a)** To study the onset lengths, the inferred posterior was marginalized over τ_*ε*_ and τ_*κ*_ to produce marginal posterior probabilities for (λ_*ε*_, λ_*κ*_ ). Color indicates highest density range (HDR) of the marginal posterior for specified total probabilities; black region sums to 0.01% of the probability. Marginal posteriors are highly localized under both an uninformative (left) and informative prior (right). **b)** Comparison of empirical DRLs to inference results under uninformative and informative priors. Counts for all DRLs are necessarily normalized for comparison (see **Methods)**; y-axis indicates normalized fractions (parentheses) and counts rescaled to match the number of repeats in the T2T genome (bold labels, black curve). Dashed line represents posterior-weighted DRL. Colored ranges represent the minimum and maximum counts at each length bin across all parameters in the specified HDR (color corresponds to panel a). Purple region shows the min-max range generated from 36 primate genomes (after removing the two most-diverged DRLs and appropriately normalizing; see **Fig. S2, Methods**). This toy model recapitulates the important qualitative features of the empirical DRL (i.e., deviation from geometric distribution at ∼10 nt corresponding to λ values, followed by an extended tail of repeats that truncates at finite length). **c)** Comparison of expansion (blue) and contraction rates (orange) to posterior-weighted length dependence for specified priors. Length dependence shows initial expansion bias and asymptotic contraction bias (the latter observable in all parameterizations).

**Fig. S15.**
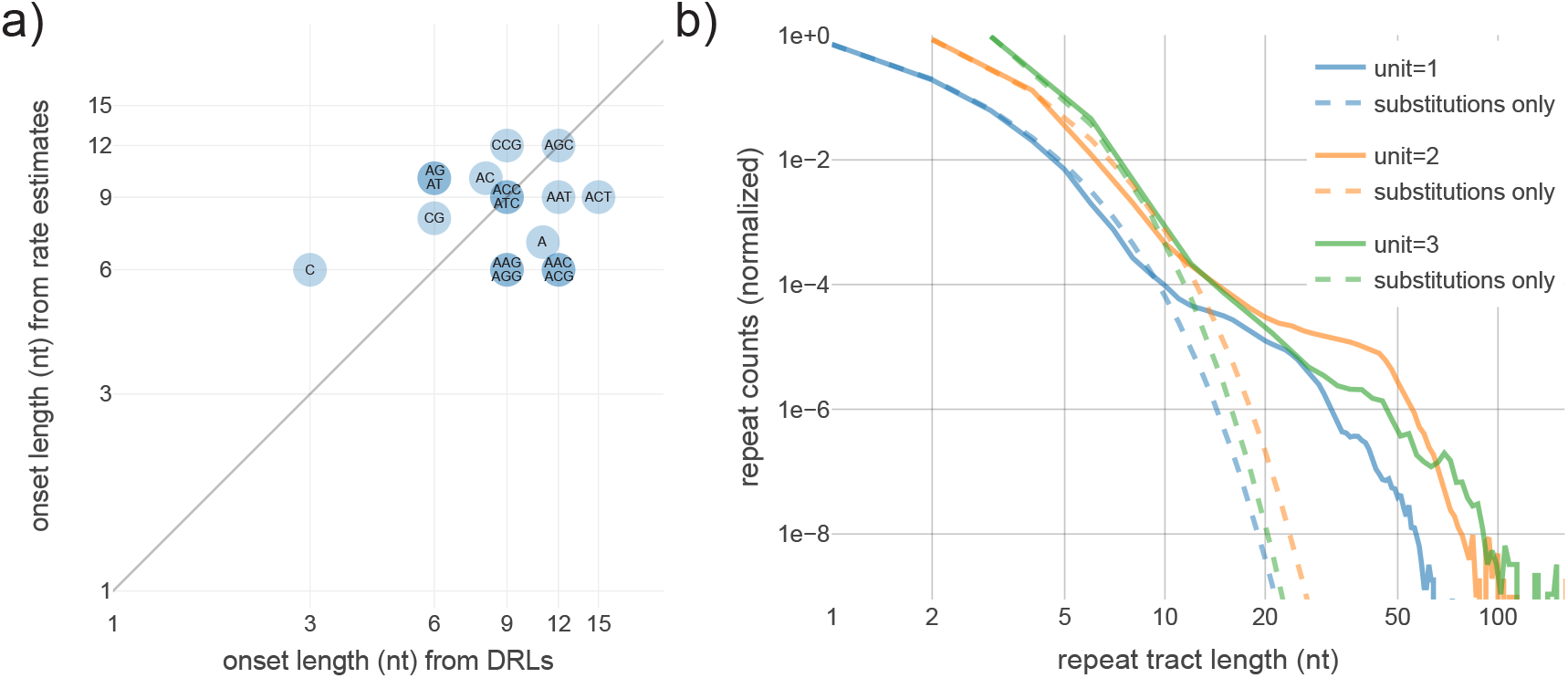
Assessing the onset length of repeat instability for longer motifs. **a)** Onset length of repeat instability as calculated from the deviation of the empirical DRL from geometric decay (see panel b) on the x-axis, and from the comparison of per-repeat expansion and contraction rates to ***μ*** substitution rates on the y-axis. For both measurements, the length is found where a two-fold deviation first occurs. Motifs are indicated in text overlays. Apart from two outliers (C and ACT), all lengths are within a range of 6-12 nt. **b)** Solid lines display normalized DRLs in CHM13-T2T summed across mono-, di-, and trinucleotide motifs. Dashed lines represent a computational model of the substitution process alone (omitting all indels). Computational models were separately run for each motif; resulting DRLs were summed according to motif length. This results in a sum of geometric distributions, which describes the low-length (i.e., <10 nt) portion of each empirical distribution. Empirical deviation from this distribution results from the transition to repeat instability at longer lengths.

**Fig. S16.**
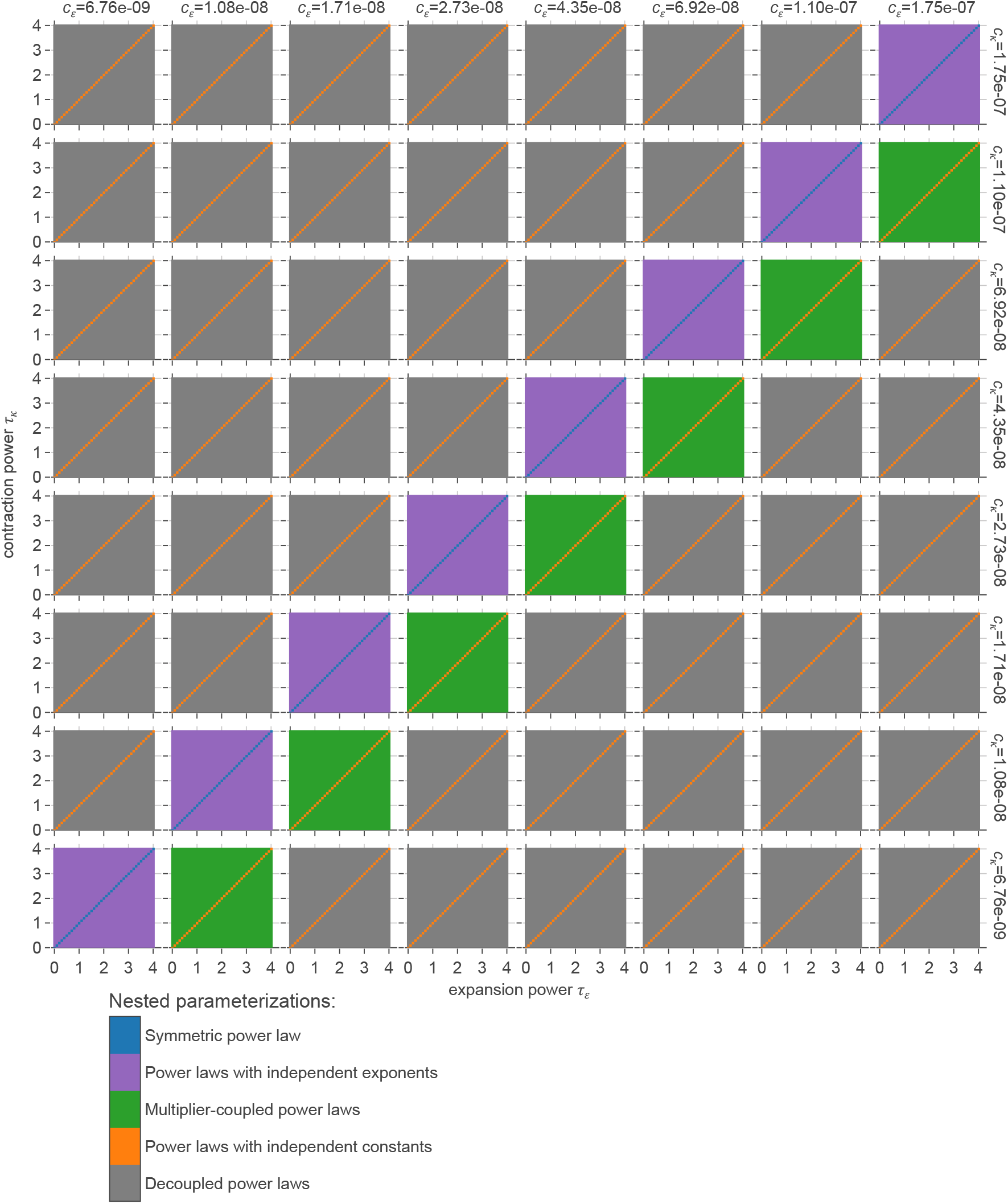
Parameterization nesting procedure. For computational efficiency and straightforward interpretation of Bayes factor ratios, all parameterizations were nested in a four-dimensional grid of parameters associated with the four-parameter decoupled power law model (all points). Each inset plane depicts a grid of (τ_*ε*_, τ_*κ*_ ) values (x-axis, y-axis) denoting the exponents of power-laws for expansion and contraction rates, respectively. Constants c_*ε*_ and c_*κ*_ that determine the expansion and contraction rates at length *L*=9 units are represented as columns and rows, respectively. Parameterizations with fewer degrees of freedom (defined in **Table 1**) are embedded as lower-dimensional slices of the four-dimensional grid and shown in specified colors. Note that the power-law model with independent constants (orange) appears disconnected in this representation but is simply connected in the discrete four-dimensional space.

## Supplementary Note SN1

### 1 Analytic modeling of the repeat length distribution in steady state

To better understand our observations, we sought to describe the equilibrium that emerges once the distribution of repeat tract lengths (DRL in main text; herein, denoted P_*L*_ in discrete form and *ρ*(*L*) in the continuum) has reached steady state at asymptotically late times (i.e., as *t*→ ∞, *P* (*L*; *t*) → *P*_*ss*_ (*L*), where *ss* denotes steady state). Importantly, for any given parameterization, this does not occur for all parameter combinations, as a subset remain inherently unstable. For the parameter values that stabilize, the shape of the distribution is described by a dynamic balance that emerges between mutational processes, with distinct mutational effects dominating the dynamics in different length regimes. For simplicity, the analysis provided here is restricted to the case of mononucleotide repeats consisting of various numbers of *A* bases. Alternative bases that terminate each sequence, labeled *B*, represent any *T, C*, or *G* base (i.e., any non-motif base for A repeats) at either end of an *A* repeat. The distribution of tract lengths for *B* strings will be ignored throughout. Further, our analytic treatment focuses on the three-parameter multiplier-coupled parameterization (see **Methods, Table 1**, Equation SN1; henceforth, ‘multiplier model’) describing the length dependence of the set of repeat instability rates; this parameterization was determined to have the strongest statistical support amongst models tested (see **Table 1, Results**). Where possible, dynamical equations are provided in general form in terms of an arbitrary parameterization of the rates.

#### 1.1 Mutational processes affecting repeats

As described in the manuscript, the mutational processes that are included in our model are the following, enumerated here with corresponding variables (*ϵ*_*L*_, *κ*_*L*_, etc.) that represent the (parameterized) length-dependent rates of each mutation type. For the reader’s convenience, a schematic representation of these mutational processes is provided in **Fig. 2a** along with corresponding effects on repeat length shown in **Fig. 2b**.

1. *Lengthening substitutions µ*: Point mutations *B* →*A* that increase the length of a repeat while preserving the total length of the genome. Such mutations only increase repeat length when they occur on bases adjacent to an existing repeat. Note that two additional transitions result from such mutations: fusion of nearby repeats (merging of two shorter repeats to form a longer repeat) and generation of new ‘repeats’ of length L=1 (included in our description for completeness, despite the lack of repetition). The distinction between these processes can be categorized by the number of *B* bases adjacent to the mutated site, one, zero, and two, for lengthening, fusion, and generation, respectively. The per-target rate *mu* is assumed to be the same constant for any repeat length *L* such that the per-repeat rate of these substitutions is necessarily linear in length, scaling as *µ* × *L* for all repeat lengths.
2. *Shortening substitutions ν*: Point mutations *A* → *B* that decrease repeat length while preserving genome length. Such mutations also result in repeat fission (interruption of a repeat that forms two smaller repeats) and repeat destruction (mutation of L=1 repeats, removing them from the distribution). Again, these transitions can be categorized by adjacency of the mutated base to one, zero, and two *B* bases for repeat shortening, fission, and destruction, respectively. The per-repeat rate of shortening substitutions is again linear in repeat length: *ν*× *L*.
3. *Contractions κ*_*L*_: Deletions of a single *A* base that decrease repeat length and genome length by one unit (i.e., one nucleotide for mononucleotide repeats). Extended deletions of two or more bases are assumed to be subdominant for simplicity (consistent with estimates provided in **Figs. 3a, S4**). In the multiplier model (see **Table 1**, Equation SN1), repeat contractions occur at a length-dependent rate constrained by empirical estimates and further parameterized by two variables; all rates for lengths *L* ≤ 8 are fixed by trio-based rate estimates (see **Methods** and **Fig. 3b**), the rate at *L* = 9 is determined by a constant multiple *m* relative to the value at *L* = 8 (i.e., *κ*_*L*=9_ = *m* × *κ*_*L*=8_), and rates for *L* > 9 follow a power law dependence of the form 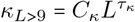. Here, the length-independent constant *C*_*κ*_ sets the initial value of the power law that parameterizes the contraction rate for all *L* ≥ 9 (for intuition, if the power law 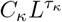 was artificially extended below *L* = 9, *C*_κ_ would be the rate associated with 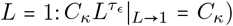 the subscript *c* refers to the crossover in behavior between substitution-dominated mutation rates at lengths *L* < 9 and the onset of repeat instability-dominated rates for *L* > 9. Under this parameterization, solving for 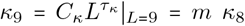 *κ*_8_ yields the definition 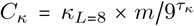, which is dependent on both *τ*_*κ*_ and *m*, but constant for a fixed parameter combination (*m, τ*_*ϵ*_, *τ*_*κ*_ ). The per-repeat target size for contractions in a repeat of length *L* is simply *L*, as the deletion of any base in an *A* repeat has an equivalent effect. Thus, the per-repeat rates are given by multiplying *κ*_*L*_ by an additional factor of *L* (i.e., the per-repeat contraction rate becomes 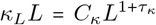 for long repeats *L* ≥ 9). For all length-dependent rates (e.g., *κ*_*L*_), we use the subscript *L* to denote the length dependence for both a discrete list of rates (e.g, *κ*_*L*_ defined in the discrete range of positive definite integers *L* ∈ ℤ^+^) and in the continuum where these rates are interpreted as continuous functions (e.g., *κ*_*L*_ →+ *κ*(*L*) becomes a continuous function describing the contraction rate in the continuous space *L* ∈ ℝ, but we preserve the notation *κ*_*L*_ ≡ *κ*(*L*) in this case for consistency).
4. *Expansions ϵ*_*L*_: Insertions of a single *A* base that increase repeat length and genome length by one unit. Again, extended insertions are assumed to occur at subdominant rates (see **Figs. 3a, S4**). Expansions are defined to be dependent on length in the same way as contractions, defining low lengths empirically and including the same value for the parameter *m*, but with a distinct power law for rates at large lengths: 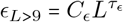. We have again expressed this dependence using the length-independent constant 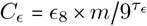 for notational convenience. We envision insertion as the replacement of a single *A* base with a pair of bases *AA* such that the per-target lengths are again multiplied by the target size *L* (e.g., asymptotically, 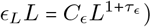.
5. *Non-motif insertions ι*_*L*_: Insertions of a single *B* base that interrupts a repeat, resulting in repeat fission. For conciseness, non-motif insertions will simply be referred to as *insertions* herein (with the implicit distinction between insertions leading to expansions and non-motif insertions). Extended insertions of more than a single base are again ignored (see **Fig. 3a, S4**). The per-target rate of insertions is again length dependent in the same way, with *ι*_*L*≤8_ derived from empirical data, with the parameterization-dependent *ι*_9_ = *mι*_8_ defined using the same multiple *m*, and rates for lengths *L* > 9 defined by the power law 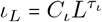. In this case, the per-repeat rate is dependent on a target size of *L* − 1 (*B* must be inserted between two *A* bases in the pre-mutated repeat and there are *L* − 1 pairs of adjacent motifs providing targets for *AA* →*ABA*). Herein, we will assume that the rate *ι*_*L*_ was estimated by collecting the average per-repeat rate of insertions and dividing by the length of the repeat *L* (i.e., as an effective per-target rate), as in **Methods**; however, if the per-target rate *ι*_*L*_ is estimated directly (i.e., by separately counting each *AA* → *ABA* event), one may resort to the approximation *L*− 1 ≈ *L* in the asymptotic large *L* regime, which is the only relevant length range for insertion-based processes due to their highly subdominant rates (relative to, e.g., the expansion or contraction rates in the same regime). We make one additional assumption here to limit the number of free parameters in the multiplier model: with empirical motivation (see manuscript text), we impose the constraint *τ*_*ι*_ = *τ*_*ϵ*_ under the assumption that they are both consequences of the same biological mechanisms such that the rates scale in parallel as 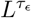, albeit with non-motif insertions occurring at significantly lower rates. Thus, the per-repeat rate of insertions in this model is given by 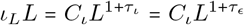 (i.e., multiplying *ι*_*L*_ by *L*, rather than *L* − 1, to recover the estimated per-repeat rate). As before, this is expressed in terms of the length-independent constant 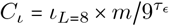, where we have substituted in *τ*_*ι*_ = *τ*_*ϵ*_.

All additional mutational processes and their impact on the repeat length distribution are assumed to be subdominant or ignored for analytic simplicity. For example, deletions of *B* bases are ignored throughout our analysis despite potential relevance to fusion rates, as the direct estimates of the deletion rate for length one *B* strings showed an orders of magnitude suppression relative to the substitution rate *µ* (a directly competing process resulting in repeat fusion; see **Methods**). We make no attempt to model the correlated dynamics of the *B* string length distribution such that insertions of *B* bases that lengthen *B* strings are also ignored, along with the insertion or deletion of extended sequences (i.e., *L* →*L* ± *k*, where *k* > 1; see above). With the exception of extended insertions and deletions, these effects were included in the results of our computational model at appropriate length-independent rates (e.g., the rate of *B* → *BB* was assumed to be the same as the rate of *BB* → *BBB*, ignoring potential repeat instability due to the indistinguishability between *B* ∈ {*C, T, G*}), but do not change the qualitative behavior of the repeat length distribution. For the multiplier-coupled model, we assume the multiplier parameter *m* is positive definite and the exponent parameters *τ*_*κ*_ and *τ*_*ϵ*_ are positive semidefinite, which together imply that *C*_*κ*_, *C*_*ϵ*_, *C*_*ι*_ > 0.

#### 1.2 Summary of the three-parameter multiplier-coupled model (for long repeats)

For the reader’s reference, we summarize the mutation rate parameterization under the three-parameter multiplier-coupled model specified in **Table 1** for each mutation type. Repeat lengths below *L* = 9 units are taken directly from empirical point estimates at each length (see **Fig. 3b**) calculated for expansion, contraction, and insertions. Substitution rates *µ* and *ν* are taken from motif-dependent empirical estimates and assumed to be length independent. For repeat length *L* ≥ 9, the per-target rates for expansion (*ϵ*_*L*_), contraction (*κ*_*L*_), and insertion (*ι*_*L*_) are parameterized as follows.

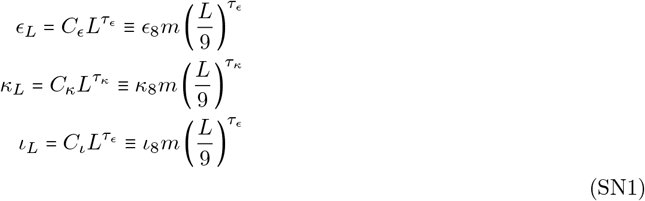

The parameter-dependent constants *C*_*ϵ*_, *C*_*κ*_, and *C*_*ι*_ are computed from *ϵ*_8_, *κ*_8_, and *ι*_8_, respectively, the empirical estimates at *L* = 8. The dependence on the number nine is a result of the largest length with reliable empirical estimates; for the purposes of the following analysis it is only important that this number is of order 10. Per-repeat rates are computed by multiplying each per-target rate by repeat length *L* (e.g., *ϵ*_*L*_*L* for expansions, etc.). Noting that we have made the simplifying assumption that insertions and expansions obey the same power law (i.e., *τ*_*ι*_= *τ*_*ϵ*_), this provides a three-parameter description of the unobserved length dependencies of the mutation rates in terms of (*m, τ*_*ϵ*_, *τ*_*κ*_). This parameterization was used in our computational model (along with other models specified in **Table 1**) to propagate the mutational process over time and substituted into more general analytic expressions below for direct comparison.

Note that it may also be of interest to consider the more general four-parameter decoupled power-law model (see **Table 1**) by using the left-side definitions in Equation SN1 (i.e., independent constants *C*_*ϵ*_ and *C*_*κ*_, with *C*_*ι*_ = *C*_*ϵ*_ /100); where possible, results that require parameterization have been expressed in terms of these more generic constants for this purpose.

#### 1.3 Mutational processes as transitions in length space (local vs. nonlocal)

Having defined each mutational process, we note that the location of the same type of mutation along the repeat sequence can result in dramatically different effects, as alluded to above. The catalogue of changes to repeat length can be separated into ***local*** transitions and ***nonlocal*** transitions (see schematic in **Fig. 2b**), which provides an important distinction in the continuum approximation described below. Local transitions result from mutations that increase or decrease length by one motif unit, while nonlocal transitions change length by more than one unit.

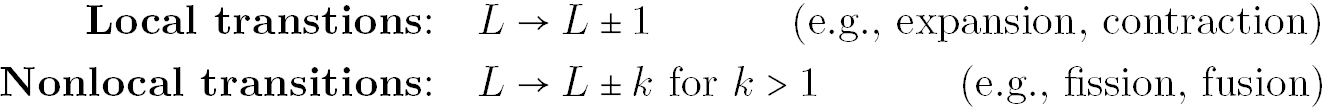

While expansion is a strictly local processes in repeat length (*L* → *L* + 1) with probabilities associated with *P*_*L*_ → *P*_*L*+1_ transitions, *µ* substitutions are local *only* when they occur at sites adjacent to exactly one repeat boundary (e.g., *BBAB* → *BAAB*, rather than *ABAB* → *AAAB*). In contrast, *µ* substitution of a base initially adjacent to two distinct repeats results in an inherently nonlocal process: two repeats (e.g., length *L*_1_ and *L*_2_) are fused into a single longer repeat of length *L*_1_ + *L*_2_ + 1. This transition is both nonlocal in length space and non-conservative with respect to the mass of the probability distribution P(L). Substitutions at bases not adjacent to repeats generate new *L* = 1 ‘repeats’ and do not conserve mass (effectively a ‘source’ at the *L* = 1 boundary). This is counteracted by *ν* substitutions (with a distinct rate) that remove *L* = 1 repeats (effectively, a ‘sink’ at the same boundary). Analogously, repeat fusion is counteracted by repeat fission due to *ν* substitutions that occur in the non-boundary bases (i.e., in the ‘body’) of the repeat; these are again nonlocal in length space and non-conservative in distribution mass.

Fission has a target size linear in the length of the initial repeat (strictly, *L* − 2, ignoring the boundaries). In contrast, fusion has a target dependent on the number of *B* strings of length one. Local decreases in length can occur due to contraction (at a per-repeat rate *at least* linear in the length) or via *ν* substitutions (at a rate 2*ν* corresponding to the finite target size of two bases for repeats *L* ≥ 2). Additional fission events can occur due to insertions. As noted above, additional fusions due to deletions of *B* bases are ignored due to negligible rates.

The nonlocal, non-conservative, and non-linear (e.g., power law dependencies and fusion rates dependent on two distinct length classes) nature of this ensemble of dynamics makes the difficulty of simultaneously modeling these effects immediately clear. Instead, motivated by the questions surrounding the maintenance and prevalence of long (and potentially disease-causing) repeats in the genome, we proceed with an analysis of the asymptotic dynamics. This approach provides a more straightforward understanding of the dominant processes appropriate in long repeat regime (i.e., *L* ≫ 1) and the contrast to forces shaping the short repeat regime (roughly defined here as *L* < 10). We first focus on an analytic description of the distribution at short lengths due to its recognizable shape and relative simplicity.

### 2 Finite difference equation for repeat length changes

We first modeled the dynamics of repeat length changes by writing a discretized finite difference equation that describes the combined effects of all five mutational processes on the distribution P_*L*_(*t*) as it evolves over time.

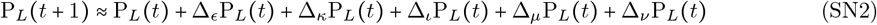

Here, we have assumed that all mutation rates are sufficiently small that the mutational processes can be linearized (i.e., the rate of multiple mutations on the same repeat in a single generation is negligibly small). This equation describes the discrete distribution at time *t* + 1 as it evolves from the distribution at the previous time step due to changes induced by expansion, contraction, insertion, and substitutions (from both *µ* and *ν* mutations), respectively. This can be rearranged to express the change to the distribution for each length bin *L* in a single generation.

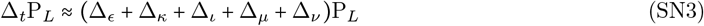

Each term in this expression is defined by the single-generation effect of a given mutational type on the number of length *L* repeats P_*L*_.

#### 2.1 Changes in repeat length due to expansion, contraction, and insertion

Expansions introduce single-unit changes in length (*L* → *L* + 1), independent of their location within the repeat (target size *L*, as defined above). The change in the number of repeats in the *L*^th^ class can be written as a combination of an influx due to insertion from below (*L* − 1 → *L*) and an outflux due to expansion to the class above (*L* → *L* + 1), as follows.

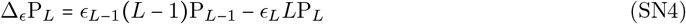

As defined above, *ϵ*_*L*_ represents the length-dependent expansion rate, which dramatically and monotonically increases above lengths of order ten. Contractions have the opposite effects: an influx from above (*L*+1 → *L*) and an outflux to below (*L* → *L* − 1), with a length-dependent mutation rate *κ*_*L*_ and target size *L*.

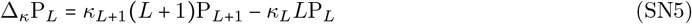

Importantly, we have omitted the effect of contraction-based repeat fusion, under the assumption that this process is subdominant to substitution-based fusion at all repeat lengths (see description of substitution-based fusion, below). (Non-motif) insertions are described by more complicated transitions, due to the position-dependent effects on length. Additionally, insertions are non-conservative, replacing one repeat with two repeats of shorter length (*L* →*k, L* −*k*, where *k* ≥1). The outflux due to insertions occurs at a length-dependent rate *ι*_*L*_ with target size *L*, which is a sum over the *L* possible locations for the insertion that result in distinguishable transitions (up to the symmetry *k* ↔ *L* − *k*; e.g., transitions 5 → 4, 2 are equivalent to 5 → 2, 4, despite distinguishable mutational targets). Note that, for the purposes of writing the finite difference equation, one need not keep track of the eventual state(s) resulting from an outflux. A focal class *L* can gain counts due to insertions in any repeat longer than *L*: insertions that occur *L* units away from either repeat boundary result in an increase in P_*L*_, such that each length class *l* > *L* has 2 potential targets for transitions *l* → *L, l* −*L* (in the case where *l* = 2*L*, there is a single target, but two length *L* repeats are added to P_*L*_). Together, the effects of insertion on a focal *L* class can be written as follows.

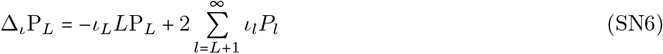

Here, we assume that P_*L*_ decays sufficiently rapidly as *L* increases such that ∑_*l*_ *ι*_*l*_*P*_*l*_ remains finite and P_*L*_ is normalizable. This sum characterizes repeat fission due to insertions alone, which are inherently nonlocal and non-conservative transitions. However, note that contributions from transitions *L* + 1 → *L* are local (though non-conservative) effects included in the same contribution. In this sense, this sum can be considered a combination of local transitions *L* + 1 → *L* (contributing 2*ι*_*L* + 1_*P*_*L* + 1_) and nonlocal transitions *l* ≥ *L* + 2 → *L* (contributing 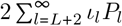); this decomposition is relevant only when insertion rates are appreciable at long repeat length, which we approximate in the continuum limit (detailed below). Similarly, the target size *L* in the outflux in Equation SN6 is a sum of targets for local and nonlocal transitions (with targets 2 and *L* − 2, respectively).

#### 2.2 Changes in repeat length due to substitutions

Substitutions behave distinctly from the above transitions, though some aspects of the transitions they induce are analogous. For example, the outflux due to shortening substitutions is analogous to insertions (with target size *L*), but with a length-independent rate *ν*; this combines substitutions at the boundary that result in local transitions *L* → *L* − 1 (with target 2) and substitutions in the repeat body (target *L* − 2) that result in repeat fission. The influx due to *ν* substitutions is again analogous to insertions and can be represented by a sum over contributions from local transitions *L* + 1 → *L* and from all fissions of repeats of length *l* ≥ *L* + 2 → *L*.

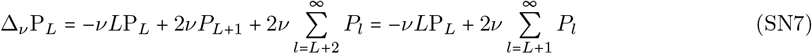

Again, we assume ∑*P*_*l*_ rapidly converges to a finite value such that P_*L*_ is normalizable, though this condition is weaker than for insertions due to the (naively) monotonic increase in the per-target rate *ι*_*L*_.

Length-increasing substitutions with length-independent rate *µ* per target result in an outflux restricted to sites adjacent to the repeat boundary (i.e., *µ* mutations *B*→ *A* at either end of *BA*…*AB*). This includes local transitions when the mutated *B* is initially adjacent to less than two *A* bases (e.g., *BBA* → *BAA* for transitions *L* → *L* + 1 or *BBB* → *BAB* for transitions *L* = 0 → 1) and nonlocal transitions when the mutated base results in repeat fusion (*ABA* → *AAA*). Here, the total target size is 2 and the net outflux is independent of the resulting repeat length(s).

The influx due to *µ* substitutions has distinct contributions from local transitions (*L*−1 → *L*) and nonlocal fusion of shorter repeats (*L*_1_, *L*_2_ → *L*, where *L*_1_ + *L*_2_ = *L* − 1). For the purposes of describing the discrete dynamics, we can introduce a time-dependent constant *p*_*F*_ (*t*) that represents the probability that mutation of a *B* base terminating a repeat results in repeat fusion, rather than a local transition; *p*_*F*_ (*t*) (henceforth, the time-dependence of *p*_*F*_ will be left implicit for brevity) is dependent on the genomic distribution of *B* string lengths at time *t* (i.e., the fraction of *B* strings of length one) or, alternatively, the probability that the three-unit context of the mutated *B* results in a substitution *ABA* → *AAA* (i.e., the probability that the mutated *B* is adjacent to two *A* repeats). The rate of substitution-based repeat fusions is therefore proportional to *µp*_*F*_ and the rate of local length-increasing substitutions is proportional to *µ* (1 −*p*_*F*_). The quantity *p*_*F*_ approaches a time-independent constant in steady state and can be measured empirically from the *B* string length distribution, computed from other genome-wide quantities like the length of the genome and total number of repeats, or computed theoretically in simple cases (see discussion below), but is ultimately unimportant to our analysis and results. The combined effects of *µ* mutations on repeats of length *L* can be characterized as follows.

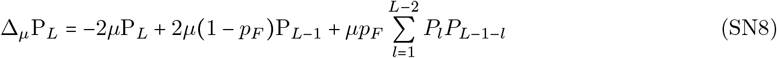

The sum in the rightmost term is the discrete form of a convolution of the distribution P_*L*_ with itself, which has an intuitive interpretation. For integers distributed according to the discrete probability distribution P_*L*_ (assuming P_*L*_ is properly normalized), the probability of randomly sampling two integers *L*_1_, *L*_2_ (<*L* −1) that sum to a fixed value *L* − 1 is given by the convolution 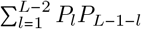; a substitution adds one unit to form a repeat of length *L*. Here, we have assumed that distinct classes of P_*L*_ remain uncorrelated in steady state for simplicity. This may be an oversimplification of the true steady state because, for example, fusion events may preferentially reverse repeat fission (i.e., fusion can re-form the original length of an interrupted repeat generated by fission; the length-dependent per-target rate of insertion-based interruptions can result in correlation between length classes, further complicating the dynamics).

We note that there can be a similar contribution that originates from deletions of a *B* adjacent to two repeats (i.e., deletion-based repeat fusion). However, based on the estimated mutation rates (see Methods), background deletions are orders of magnitude less frequent than *µ* substitutions such that deletion-induced fusion transitions occur at negligible rates.

#### 2.3 Finite difference equation

Summing the above contributions to ΔP_*L*_ yields the full finite difference equation describing the change in each length class P_*L*_ subject to two-way substitutions, expansions, contractions, and insertions.

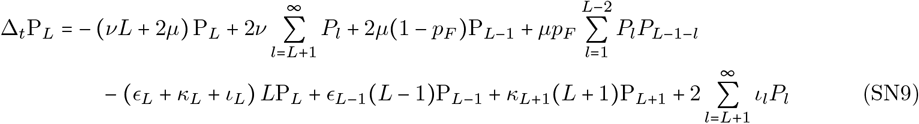

For clarity, the first line of terms summarizes changes due to substitution and the second summarizes insertion- and deletion-based changes due to repeat instability. Inclusion of deletion-based fusion and/or correlations between length classes, both of which are treated as negligible, would add an additional convolution and/or a dependence on the covariance between length classes, respectively.

Guided by striking differences in the length-dependencies of substitutions and repeat instability rates (**Fig. 3b**), we further analyze the steady state dynamics by separating into distinct length regimes in which a subset of mutational processes dictate the vast majority of changes in length. Under this separation of length scales, the dynamics of short repeats (roughly, *L* ≤ 8 for mononucleotide-A repeats) and long repeats (roughly *L* > 10) are largely dominated by distinct mutational forces: the distribution of short repeats is maintained in a dominant balance between the opposing effects of *µ* and *ν* substitutions (including both fission and fusion), while length changes to long repeats are dominated by a (parameter-dependent) balance between expansions, contractions, and fission (potentially including fission resulting from *ν* substitutions). The assumption of dominant balance allows for a dramatic simplification of the full set of contributions to Equation SN9; due to their low rates, all neglected terms provide minor corrections to the resulting approximation to the steady state distribution, which we demonstrate post-hoc.

Equation SN9 represents the deterministic change of the repeat length distribution in a reference sequence (i.e., a single individual) due to mutations alone. Here, we have assumed that natural selection is absent such that the accumulation of mutations in the reference results from mutations aggregating along the lineage ancestral to this individual. This process occurs over sufficiently long times that a steady state is eventually reached. Once in steady state, the time-averaged distribution of repeat lengths (i.e., averaging over stochastic behavior) can be equated to the steady state distribution obtained from Equation SN9.

### 3 Steady-state dynamics in the short repeat length regime

In contrast to previous studies, the empirical distribution we constructed (see **Fig. 1b**) includes very short contiguous sequences and, notably, single base counts for comparison. The relative rates of expansions, contractions, and insertions to both types of substitutions makes it clear that repeat instability is are largely irrelevant to the maintenance of such short sequences (see **Fig. 3b**). From a biological standpoint, this implies that repeat instability is technically irrelevant (i.e., occurs at highly suppressed rates) until repeats exceed roughly *L* = 8 (this differs slightly between motifs of distinct lengths and is more accurately described as roughly 6-12 nucleotides, rather than repeat units, for motifs of unit length *l*_*m*_ < 4 nucleotides; see **Fig. S15**,); this corresponds to the length range where expansion and contraction rates are comparable to substitution rates such that repeat instability becomes relevant to the dynamics.

The shape the distribution of short repeats is well-approximated as a balance between substitutions alone (see, e.g., black vs. dashed blue lines in **Fig. 6**). The distribution is generated by the random process of sequence evolution under two-way substitution with distinct rates *µ* and *ν*. Given indefinite time, this process equilibrates to a steady state genome in which the probability of randomly sampling an *A* base is given by the fraction *p*_*A*_ = *µ* /(*µ*+ *ν*) and the complementary probability of sampling a *B* base is given by *p*_*B*_ = 1 − *p*_*A*_ = *ν*/(*µ* + *ν*). Here, the relevant substitution rates *µ* and *ν* are the single-unit context rates (i.e., averaged over longer contexts) *µ*_*B*_→_*A*_ and *µ*_*B*_→_*A*_, respectively, as this distribution is not characterized by distinctions between local transitions, fission, and fusion. Under this model, a repeat is simply a contiguous sequence of *A* bases that is *L* bases (or repeated units) long and adjacent to a *B* base on either side. Conditioning on an initial *B* base, a length *L* string of *A* bases occurs at a frequency approximately given by the probability of randomly sampling an *A* base *L* successive times, followed by a terminating *B* base. The frequency of a length *L* repeat is therefore given by geometrically distributed distribution proportional to *p*_*A*_^*L*^*p*_*B*_.

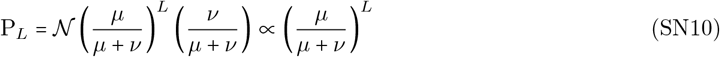

Here, 𝒩 is a normalization constant defined as 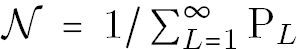, where Σ _*L*_ P_*L*_ is the total mass of the distribution. As the steady state distribution is no longer geometrically distributed when repeat instability becomes relevant (roughly above *L* = 8), the value of 𝒩 cannot be determined by the short repeat dynamics alone; the normalization can only be evaluated after identifying the steady state distribution over all length classes. Given that the constant 𝒩 is unknown, the constant probability associated with sampling repeat-terminating *B* bases (i.e., the factor of *p*_*B*_) can be absorbed into the definition of 𝒩.

#### 3.1 Geometric solution to the substitution-only difference equation

To demonstrate the utility of the finite difference equation in a simpler setting (without repeat instability), one can show that the geometric distribution, when normalized, provides the solution to the short length approximation to Equation SN9 in steady state. Under substitutions alone, the short length regime is well-approximated by the following difference equation.

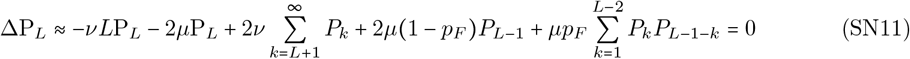

The finite difference equation becomes a steady state condition by imposing time independence (i.e., ΔP_*L*_ = 0). For consistency with Equations SN7 and SN8, the local influx from *ν* substitutions is included in the sum representing fission and the parameter *p*_*F*_ =(−1 *q*_*F*_) will be defined below to represent the appropriate rate of fusion in the present context. The normalized geometric distribution is given by the following expression in terms of *p*_*A*_ and *p*_*B*_ =1− *p*_*A*_, defined above.

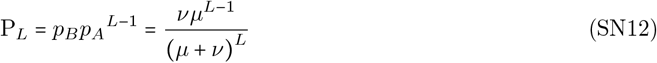

Though fusion and fission are inherently non-conservative transitions that change the number of repeats in the distribution, we make the approximation that individual fission and fusion events do not significantly alter the normalization constant in steady state. The term representing fission is now the incomplete sum over a geometric series, which can be explicitly evaluated as follows.

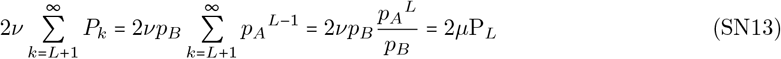

Thus, the influx due to both repeat fission and local transitions, both *ν* substitution-based effects, exactly cancel the outlux due to *µ* mutations.

To evaluate the convolution term, note that the probability distribution for the length of *B* strings under two-way substitution 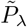 is geometric by the same arguments as for the *A* distribution, but with reversed probabilities *p*_*A*_ ↔ *p*_*B*_. Normalizing string lengths *λ* = [1,∞), (i.e., conditional on a *B* string of length *λ* ≥ 0), the normalized probability distribution is as follows.

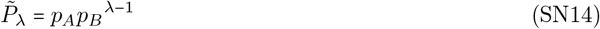

The probability that any given *A* string is terminated by a *B* string of length *λ* = 1 is simply 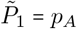. This determines the steady state probability of fusion *p*_*F*_, for which a *µ* substitution in a length one *B* string results in the transition *L*_1_, *L*_2_, → *L*_1_ + *L*_2_ + 1. The complement is the probability of locally increasing the length by a single unit (as opposed to fusion) due to an adjacent *µ* substitution, 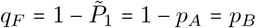 After cancelling the *ν* substitution influx with the *µ* substitution outflux, we can rewrite the remaining terms in Equation SN11 by substituting in *p*_*F*_ =*p*_*A*_, *q*_*F*_ = *p*_*B*_ and evaluating the partial sum in the fusion term.

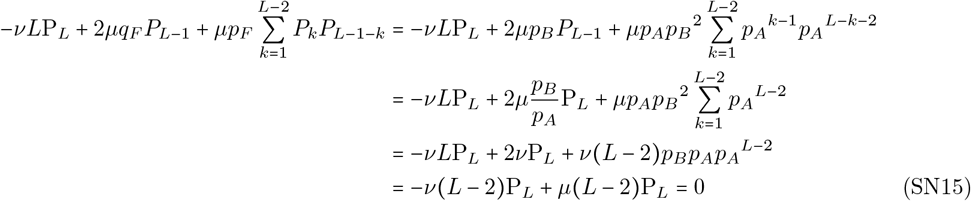

We find that a geometric distribution of both *A* and *B* repeats satisfies the steady state equation under two-way substitution, which justifies the aforementioned approximation for the substitution-dominated short length regime of the distribution.

#### 3.2 Interactions between short and long repeats are largely restricted to boundary effects

Here, we are only concerned with the geometric falloff in Equation SN10 that defines the shape of the distribution in this length regime, which is given by the proportionality on the right hand side of the above expression. Importantly, this length dependence remains largely independent of the dynamics of the long length regime, which only affects the normalization constant. This separation of the dynamics for short and long length repeats is a reasonable approximation because the rate of transitions between these regimes is low and primarily limited to the intermediate lengths at *L* ∼ 8 − 10 (i.e., the boundary between the short and long repeat regimes). As discussed below, we allow for an influx from the long length regime but note that it is inherently negligible because the relative mass of short length repeats vastly outweighs the mass in the long repeat tail of the distribution. In this sense, the short repeat regime can be treated as a probability sink for the long repeat distribution for length transitions that exit the long length regime. Similarly, the mass in the long length regime is sourced by short repeats that approach the length regime boundary and are quickly subject to expansion-biased instability; this can be considered a local conditions on the boundary between the regimes without altering the dynamics elsewhere. If the steady state is in detailed balance (i.e., the flux in and out of each length class vanishes independently), fluxes at the length regime boundary must cancel such that both ends of the distribution maintain their relative weights.

While it is immediately clear that the distribution of *A* repeats deviates from a simple geometric decay due to the combined action of expansion, contraction, and insertion in the long length regime, the *B* string length distribution is unaltered by expansion and contraction, which solely manipulate *A* repeat length. While likely unrealistic, we proceeded under the assumption that *B* strings do not constitute repeats and are therefore not directly subject to repeat instability. Therefore, any deviation from a simple geometric distribution can only be due to the effects of insertion, which generates new length one *B* strings during each transition. Although the additional source of *B* strings will contribute to the total number of *B* strings in the eventual steady state balance between the *A* and *B* distributions, localization of this influx to the lowest length class (*B* strings of length *L* =1) implies that the normalized distribution of *B* lengths remains approximately geometric.

### 4 Steady-state dynamics for asymptotically long repeat lengths

The steady-state distribution at long lengths has a qualitatively distinct shape from the short length regime, exhibiting a much heavier tail and slower decay than the geometric distribution. Although our de novo rate estimates do not cover the entirety of this range, we observed an initially rapid, monotonic increase in the per-target expansion and contraction rates with length; this must eventually reach the well-documented extreme rates observed in repeat expansion disorders. Assuming sustained, naively monotonic rate increases with length, per-target repeat instability rates must eventually exceed substitution rates due to the stronger dependence on length (this is already observable at lengths of order 10 in **Fig. 3b**). Above some sufficiently long length, the instability-based length changes comprise the majority of mutations, dominating the mutational dynamics; this suggests that the long repeat regime of the steady-state DRL is shaped by the mutational processes with the strongest length dependencies. However, the point at which substitution-based fission (which scales linearly with repeat length) can be ignored is entirely parameter dependent. The forces relevant to the longest length repeats in the DRL are thus some subset of expansion, contraction, substitution-based fission, and insertion-based fission (or the combination of all four).

To disentangle the relative importance of each mutational process, we employed a large *L* continuum approximation to the finite difference equation to analyze the long-length dynamics. For long repeats, *L* ≫ 1 such that the distance between length bins can be treated as infinitesimal; this allows for a continuous approximation to the discrete Equation SN9 appropriate only for repeat lengths of at least order 10. The distinction between local and nonlocal terms becomes important in the continuum, as local differences are approximated by continuous derivatives, while the nonlocal sums in Equation SN9 (i.e., mutational transitions from across the rest of the distribution) are approximated as integrals.

#### 4.1 Local contributions to the change in P_*L*_

We first focus on local changes to the length distribution due to each of the forces. Starting with expansion in Equation SN4, the discrete change to a focal length class P_*L*_ is the difference between an influx due to *L* − 1 → *L* length-increasing transitions and an outflux due to *L* → *L* + 1 length-increasing transitions. The strictly local nature of expansion allows for a local approximation to Δ_*ϵ*_P_*L*_ using a Taylor expansion in the continuum. To elucidate the continuum limit, we treat expansions in detail, below, but note that the approximation for all local contributions can be obtained analogously. We first define the distance between discrete length classes Δ*L* = 1, which suggests that we must change units before formally taking the continuum limit and assessing the accuracy of the resulting approximation.

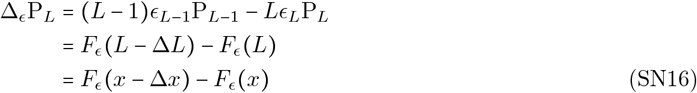

Here, *ϵ*_*L*_ is the length-dependence of the expansion rate (e.g., 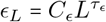 for *L* ≥ 9 under the power-law model in Equation SN1). The function *F*_*ϵ*_ (*L*) = *ϵ*_*L*_ *LP*_*L*_ was defined solely for notational convenience. In the third line, repeat length *L* was rescaled under a change of variables defined by *x* = *L*/*c* (with Δ*x* = Δ*L*/*c* = 1/*c*); *c* is an arbitrary dimensionful constant with the same units as *L* (i.e., measured in a number of motif units) such that *x* is dimensionless. If *x* is kept fixed while *L* grows large, Δ*x* gets increasingly smaller. Thus, choosing an appropriate *c* allows the backward finite difference Δ*F*_*ϵ*_ (*x*) = *F*_*ϵ*_ (*x* − Δ*x*) − *F*_*ϵ*_ (*x*) to be approximated by a Taylor expansion in small Δ*x* truncated at a some finite order (for all practical purposes, this is equivalent to a Taylor expansion around ‘small’ Δ*L* after setting Δ*L* = 1 when *L* ≫ 1, but side steps the concern that Δ*L* = 1 motif unit is definitionally not infinitesimal). Taking the continuum limit *L* → ∞ and Δ*x* → 0, *F*_*ϵ*_ becomes the continuous function *f*_*ϵ*_ and the discrete derivative approaches the continuous derivative Δ*F*_*ϵ*_ → ∂_*x*_*f*_*ϵ*_. For finite *L* (and thus finite *c*), the first-order continuum approximation is accurate to order Δ*x*^2^.

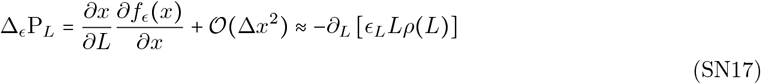

This is expressed in terms of a general, length-dependent expansion rate *ϵ*_*L*_ (at sufficiently long lengths) to allow for generic parameterizations, including Equation SN1. To arrive at this approximation, the variable change was inverted (i.e., imposing *L* = *cx*, where ∂*L*/∂*x* = 1 /*c*; the chain rule is explicit in the middle expression for clarity) after taking the continuum approximation. Here, the notation 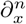 represents the *n*^th^ derivative with respect to *x* and will be used henceforth for brevity. In the length continuum, we assume that the discrete distribution P_*L*_ can be well-approximated by the continuous, differentiable function *ρ*(*L*) (defined in the rescaled length units *ρ*(*x*) ≡ lim_Δ*x*→0_ *P* (*x*)) such that the derivative ∂_*L*_*ρ*(*L*) represents the flux through length *L*. The continuum approximation is only applicable for *L* ≫ 1 and all expressions dependent on *ρ*(*L*) should be considered implicitly conditioned on *L* ≳ 10 such that the long-length scaling behavior of the mutation rates can be used directly (i.e., empirically-derived rates for *L* < 9 can be ignored such that length scaling for each rate is dictated by the parameterization, e.g., Equation SN1). Importantly, at finite *L*, corrections to the lowest-order approximation to the finite difference Δ*F* (*L*) for a discrete function *F* (*L*) cannot be treated as negligible when ∂_*L*_*f* (*L*) vanishes (see discussion of second-order corrections, below). Equation SN17 describes a strictly local transition that drives a length-dependent flux through the focal length *L*, which represents the collective impact of expansion rates on the distribution; if desired, the chain rule can be applied to show a separate flux and length-dependent loss of mass due to expansion (i.e., Δ_*ϵ*_P_*L*_ ≈ −*ρ* (*L*) ∂_*L*_ (*ϵ*_*L*_*L*) − *ϵ*_*L*_*L*∂_*L*_*ρ*(*L*)). Noting that the derivative is negative throughout the roughly monotonic empirical distribution, the flux due to expansion may describe an overall increase or decrease in length, depending on the relative magnitude of the non-derivative contribution.

The effects of contraction Δ_*κ*_P_*L*_ can be summarized analogously in terms of contributions to and from the adjacent shorter and longer length bins, respectively, which becomes a local length-dependent flux in the opposite direction in the continuum limit. This sign difference emerges because the continuum approximation is to a forward finite difference for contractions and to a backward finite difference for expansions (i.e., the difference between continuum approximations for *L* + 1 → *L* and *L* − 1 → *L* transitions).

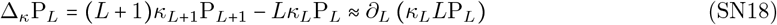

Like expansions, contractions amount only to local changes to the distribution, but the length dependence of the per-repeat rate remains important to our understanding of the dynamics.

In contrast, substitutions induce both local and nonlocal transitions. We first isolate the local transitions from in Equation SN7 and SN8, treating the nonlocal contributions as fissions and fusions (see below). In discrete form, the local influx and outflux towards increasing length due to *µ* substitutions and towards decreasing length due to *ν* substitutions can be represented as follows.

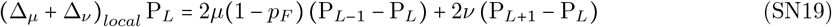

Unlike expansion and contraction, the target size for local changes in length due to substitution is a length-independent factor of 2 associated with mutations at either boundary (e.g., *AAA* → *AAB* and → *BAA* for *ν* substitutions; *µ* substitutions reverse these transitions), which have a distinct effect from mutations at non-boundary loci. The target size for *µ* substitutions is reduced further because a fraction *p*_*F*_ transitions on the boundary result in fusion, rather than local increases in length (i.e., leaving a target of 2 (1 −*p*_*F*_) = 2*q*_*F*_ per repeat). The first-order continuum approximation to Equation SN19 is the following.

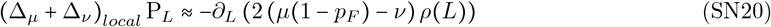

Insertions include a similar local term, but this comes with an additional complication. If an insertion occurs adjacent to either boundary (with target size 2: *AA*…*AA* → *ABA*…*AA* and → *AA*…*ABA*), the resulting repeat fission desribes the replacement of one length *L* repeat with one length *L* −1 repeat and one length 1 ‘repeat’ (*L* → *L* − 1, 1). This is the only example of repeat fission that also contains a local transition, but will not be explicitly accounted for in our treatment of insertion-based fission below. The local component of this contribution can be written as follows.

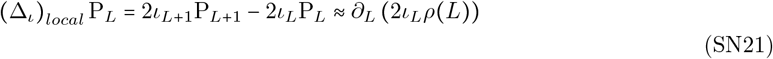

Due to the limited target of two possible insertions resulting in this local transition, the appropriate length dependence inside the derivative is twice the per-target rate *ι*_*L*_, rather than a dependence on the per-repeat insertion rate. The correlated nonlocal contribution to the *L* = 1 bin can be ignored because the relatively low rate of insertions results in an exceedingly small influx into *L* = 1 relative to the geometrically distributed mass maintained by substitutions.

##### 4.1.1 Competition between first-order local effects

The length-independent (and modest) target size, matching power law exponent (assumed for all parameterizations), and globally suppressed rate of insertions relative to expansions (based on empirical estimation, where possible; see **Fig. 3b**) together ensure that local insertions remain subdominant to expansions (i.e., (Δ_*ι*_)_*local*_ P_*L*_ ≪ Δ_*ϵ*_P_*L*_ for any *L*); as a result, local insertions may be neglected entirely. By the same argument, and given that expansion and contraction rates far exceed the substitution rates at long lengths, local changes in length due to substitutions are likely negligible in the *L* ≫1 regime of interest. Additionally, we will at this point treat time as continuous (i.e., assuming infinitesimal generation time after appropriately rescaling the units of all rates) such that the per-generation change in occupancy of P_*L*_ can be approximated by the time-derivative ∂_*t*_*ρ* (*L*) (note that we revert back to the notation ΔP_*L*_ when referencing the discrete equations). Under these approximations, the first-order approximation to the local dynamics can be summarized by the following.

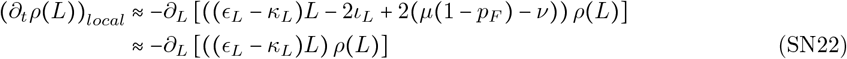

The quantity *ϵ*_*L*_ − *κ*_*L*_ represents the bias between expansion and contraction at a given length; the lowest-order approximation to the local dynamics thus describes a flux through each length class due to the length-dependent expansion-contraction bias.

The sign difference between the expansion and contraction terms suggests that, if their magnitudes are nearly identical at a given length (i.e., (Δ_*ϵ*_ +Δ_*κ*_) *P*_*l*_ ≈ 0 such that length changes are nearly symmetric at length *l*), contributions from insertion and/or substitution, which subdominant otherwise, could become relevant (in principle). Under the somewhat simplistic multiplier-coupled parameterization in Equation SN1, this can occur only at a single, specific (but parameter-dependent) length *L*\*.

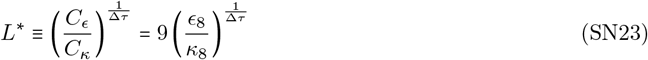

We have expressed this in terms of the collapsed parameter Δ*τ*, the difference between the contraction and expansion exponents (referred to extensively below).

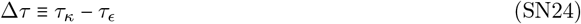

*L*\* is independent of the multiplier *m* and dependent only on Δ*τ*, rather than on *τ*_*ϵ*_ and/or *τ*_*κ*_ individually. Note that the factor of 9 in Equation SN23 is simply a consequence of defining the multiplier model in terms of the estimated rates *ϵ*_8_ and *κ*_8_ (and qualitatively unimportant). While this quantity is notable because of the potential for subdominant local contributions (from insertion and substitution), second-order corrections to the finite difference equation (due to expansion and contraction) dramatically exceed these effects due to their scaling with target size. However, *L* also aids in qualitative interpretation because it describes the transition from expansion- to contraction-biased dynamics in this parameterization.

##### 4.1.2 Second-order corrections to the local behavior and diffusive dynamics

When the first-order approximation to the local dynamics vanishes (due to nearly symmetric expansion an contraction rates across some range of lengths), subsequent corrections to the continuum approximation at finite *L* can dominate the leading-order local behavior. The computationally-explored parameter space (i.e., values of (*m, τ*_*ϵ*_, *τ*_*κ*_) in the multiplier model) included *L*\* values well-within the populated range of the long length tail of the DRL. In such cases, further corrections to the local behavior are necessary to characterize the dynamics in the continuum approximation.

With this in mind, we obtained the next-order approximation (i.e., the first strictly non-vanishing contribution) to Equation SN9 by Taylor expanding the expressions for expansion, contraction, insertion, and substitution to second order in the continuum limit. For expansion, we approximate Equation SN17 to order Δ*x*^2^ in the dimensionless variable *x* (i.e., the appropriately rescaled *L*) to produce the following approximation.

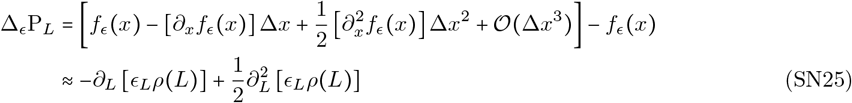

All other mutational effects can be expanded to the same order analogously. Importantly, the second-order contribution is positive-definite for each mutational process such that their sum is non-vanishing (in contrast to the sign difference that allows the first-order term to vanish at *L*\*). This results in the following approximation to the continuous-time local dynamics in the large length regime.

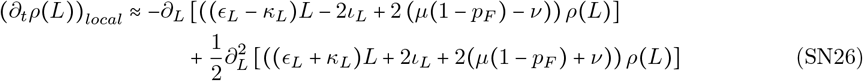

This expression provides a continuum approximation for all *local* length transitions in terms of arbitrary, length-dependent mutation rates (estimated directly, or parameterized). Higher order corrections would only become necessary in the neighborhood of a length at which the first and second derivative terms both vanish (for example, in the unrealistic scenario where extrema in the length dependencies *Lϵ*_*L*_ *ρ*(*L*), *Lκ*_*L*_*ρ*(*L*), *ι*_*L*_*ρ* (*L*), and *ρ*(*L*) occur at the same length). Using the multiplier model in Equation SN1, the relative importance of each mutational process to local transitions can be assessed by contrasting their relative magnitudes and length dependencies.

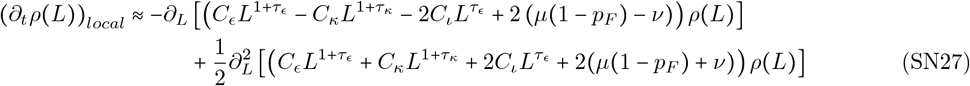

The second derivative terms in this approximation can be collectively viewed as diffusion-like changes to repeat length. Unlike bias-driven fluxes from the first derivative, the second-order term represents symmetric, bidirectional transitions (i.e., equiprobable local increases and decreases in length). Due to the monotonic increase in instability rates, these diffusion-like effects grow stronger at longer lengths. Relative to expansions and contractions, substitutions and insertions contribute negligibly to the second derivative; both contributions have a finite target sizes and, despite increasing per-target rates, insertion occurs far less frequently than expansion at all lengths (i.e., *C*_*ι*_ ≪ *C*_*ϵ*_ such that *ι*_*L*_ ≪ *ϵ*_*L*_, independent of parameter values). The local dynamics are therefore well approximated by the following expression, dependent only on expansion and contraction.

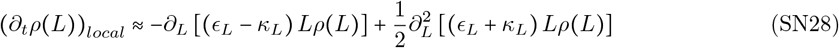

This general form can again be evaluated under any parameterization (e.g., Equation SN1) to approximate the ensemble of local transitions for long repeats, provided the assumption that expansion exceeds insertion remains valid. These local changes in length are a competition between bias-driven directional flux and diffusive changes in length, both generated primarily by expansion and contraction. Diffusive effects necessarily dominate when bias vanishes (e.g., in the neighborhood of *L*\*); at all other lengths, the relative importance of these effects is a parameter dependent competition between the net expansion-contraction bias (i.e., asymmetric component of instability) and the average magnitude of expansion and contraction (symmetric component).

#### 4.2 Repeat fission as a nonlocal contribution to changes in length

In contrast to the local effects described above, repeat fission and fusion are inherently nonlocal processes. Fission is a consequence of interruptions due to insertions or *ν* substitutions that generates both a flux out of the focal *L* class (producing two shorter repeats) and a corresponding flux into the same class from fissions of longer repeats; the total number of repeats is altered by the fission of one contiguous repeat into two. Conceptual distinctions between substitution and insertion in this context are minor in the continuum: insertions allow transitions to and from adjacent length classes (e.g., *L*→ *L* − 1, 1), while substitutions require transitions to and from at least two length bins away (e.g., *L* → *L* − 2, 1). Additionally, insertion within a repeat necessarily results in fission and does not conserve total genomic length (though, this does not *a priori* alter the mass of the *A* repeat distribution), unlike substitutions. The most pertinent difference owes to the length-dependent per-target rate of insertions, which can dramatically increase the number of such events as repeat length increases, despite initially much lower per-site rates at lower repeat lengths (see **Fig. 3b**). In both cases, the nonlocal nature of fission-generated length transitions complicates the modeling of the dynamics, as the associated influx from all higher length bins amounts to a sum of distinct contributions to the change in P_*L*_ at each length (see Equations SN6 and SN7).

##### 4.2.1 Substitution-based fission

Fission occurs only as a consequence of *ν* substitutions, with no dependence *µ*. After removing the target for local transitions at the repeat boundaries (see Equation SN19), the remaining target for nonlocal transitions is *L* − 2, corresponding to the body of the repeat. While the local transitions generate derivatives in the continuum (Equation SN26), the remaining terms in Equation SN7 result in an integral over the distribution of repeats longer than the focal class P_*L*_.

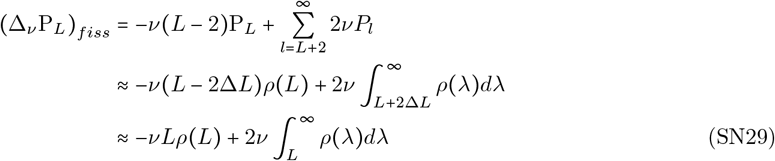

Here, the lower limit of the sum differs from Equation SN7 after removing local transitions *L*+1 → *L*. However, the continuum approximation applies only to the asymptotically long length regime where *L* ≫ Δ*L* = 1; in this limit, the approximations *L* + 2Δ*L* ≈ *L* (integration limit) and *L* − 2Δ*L* ≈ *L* (target size for outflux) were used to obtain the final line, above. The linear length scaling in the outflux represents the fact that any substitution in the repeat body alters the repeat length. In contrast, there are exactly two fission targets that generate transitions from each longer length class: specific substitutions *L* units away from either boundary of a length *λ* repeat result in a shorter repeat of length *L* (along with one of length *λ* − *L* −1, which does not add to the length *L* bin). In the special case where *λ* = 2*L* + 1, there is only one relevant target for substitution (the middle base), but this event generates two length *L* repeats, rather than one. All longer repeats therefore contribute identically to the influx, independent of length, resulting in an integral over the distribution of longer length bins. The net direction of fission (i.e., influx minus outflux) depends on the focal length and the number of longer-length repeats in the distribution tail (i.e., how rapidly it decays and truncates).

##### 4.2.2 Insertion-based fission

Fission due to insertions can be described analogously by taking the continuum limit of the nonlocal component of Equation SN6 after removing the local transitions included in Equations SN21 and SN26. Note that, in this case, the length dependence of the per-target rate *ι*_*L*_ appears under the sum (and integral).

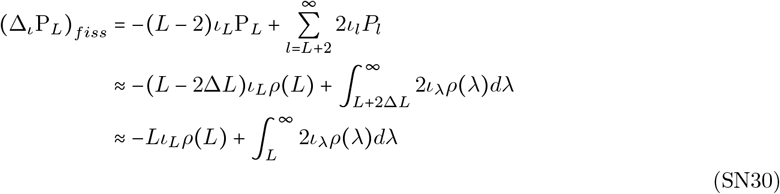

Analogous to substitutions, the target size *L −* 2 and summation limit *l = L* + 2 refer only to nonlocal transitions away from the repeat boundary, both of which are approximately *L* in the *L* ≫ 1 continuum. The resulting contributions resemble Equation SN29, but the length dependence of *ι*_*L*_ alters the length scalings (e.g., under the multiplier model, the outflux scales as 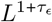, rather than *L*). The influx is instead a weighted integral over the longer-length distribution tail; the net direction of fission therefore depends on the length dependence of *ι*_*L*_ (e.g., net fission in the multiplier model depends on the parameter *τ*_*ϵ*_). This difference in overall length scalings can result in a competition between substitution and insertion that can transition from substitution-dominated to insertion-dominated fissions as the focal length is increased.

#### 4.3 Repeat fusion under random sampling of the length distribution

Repeat fusion, the process by which either *µ* substitutions or, at a lower rate, deletions of length one *B* strings (henceforth *B-dels*) result in the merging of two shorter repeats into a longer repeat, substantially complicate our model of repeat length dynamics. For simplicity, we only describe *µ* substitution-based fusion (as shown in Equation SN8), but note that differences between substitution- and B-del-generated fusion are entirely analogous to those between substitution- and insertion-generated fission. The only notable exception is that we found no evidence that the rate of deletions between repeats (namely, deletions of single-nucleotide or single-unit interruptions) harbor any dependence on the length of the repeat. Additionally, empirical estimates of *de novo* rates from trio data indicate that the per-target rate of B-dels (roughly 2 × 10^− 10^ per generation) is suppressed relative to the per-target rate for *µ* substitutions (roughly 4 × 10 ^−9^ per generation; see **Methods**) by more than an order of magnitude, suggesting B-dels provide an entirely negligible correction to substitution-based fusion rates. Henceforth, all discussion of fusion is focused on substitution-generated events.

Fusion can only occur due to mutations at *B* sites immediately adjacent to two *A* sites and thus occur at a rate proportional to the fraction *B* strings of single-unit length *p*_*F*_ (see Equation SN8 and subsequent discussion below Equation SN14). In the absence of insertions, this probability is given by *p*_*F*_ = *µ*/ (*µ*+ *ν*) (expansions and contractions of *A* repeats do not alter this rate). Although insertions could alter this rate, their inclusion results in a (slightly) greater influx into the *L* = 1 class of *B* strings. This amounts only to an additional source at the low length boundary; while this alters the total number of *B* strings, the normalized probability distribution remains geometric (i.e., Equation SN14) with a slightly modified rate constant. We confirmed via our computational model that this geometric distribution is nearly identical to that under two-way substitution alone, due to the overwhelming mass in the length one class and negligible influx due to the low overall insertion rates. However, the computational model is an abstraction that assumes *B*-strings are not susceptible to repeat instability. In practice, the value of *p*_*F*_ could be better estimated from the empirical distribution of *B* strings, but this was unnecessary in the current setting, as any such estimate does not impact our analysis or results.

We proceed by taking the continuum limit of Equation SN8 after removing local contributions (i.e., those proportional to (1 −*p*_*F*_)).

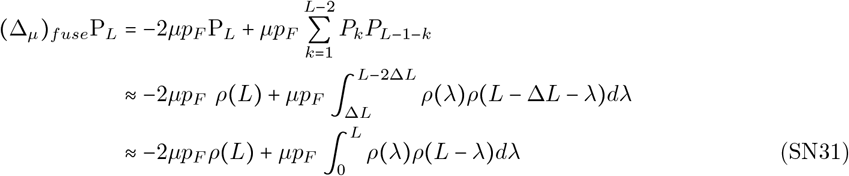

As written, the sum contains an implicit factor of two associated with swapping the subscripts *k* ↔ *L* −1 − *k* (i.e., double counting when summing up to *k* = *L* − 2), which remains in integral form (i.e., *λ* ↔ *L* − *λ* when integrating to *L*). The expression for fusion is explicitly nonlocal due to the quadratic, integral dependence on the distribution, which describes randomly sampling two shorter repeats of appropriate lengths. In the second line, we have again taken the large *L* asymptotic limit *L* ≫ Δ*L* = 1 to suppress subdominant terms. In this form, it becomes clear that the integral is simply a convolution of the distribution *ρ* (*L*) with itself over a finite length window; this can be readily interpreted as the distribution of the sum of two random lengths drawn from the same probability distribution *ρ* (*L*) (albeit not necessarily normalized appropriately due to non-conservative transitions). Noting that *p*_*F*_ < 1 and *µ* < *ν*(from empirical estimates), the fusion outflux is strictly less than the outflux from substitution-based fission alone; this contribution becomes negligible in the long length regime due to the finite target size for fusions (i.e., 2*µp*_*F*_ ≪ *νL*). However, the integral terms cannot be analogously compared due to their distinct functional forms and non-overlapping limits.

Due to the substantial analytic complications introduced by repeat fusion, we turned to our computational model to quantify its effects across the long repeat regime. Using the three-parameter multiplier-coupled model (see Equation SN1), we produced final-time point flux plots (e.g., those shown in **Fig. 6c**) across the parameter space to assess the relative effects of each class of transitions at each length. While fusion generated prominent transitions in the short repeat regime (maintaining the geometric decay detailed above), it remained negligible at longer lengths, regardless of parameter combination. We confirmed this observation by exploring parameters within the 95% HDR of our alternative parameterizations (see **Table 1**), which reinforced this observation. This suggested that, at least in steady state, the dominant dynamics in the long repeat regime remain largely unaffected by infrequent transitions due to repeat fusion.

#### 4.4 Steady-state condition for long repeat dynamics

Collecting the continuum approximations to each term of Equation SN9, the continuous-time dynamics of long repeats in the asymptotic large *L* regime can be described by the following partial differential equation (PDE).

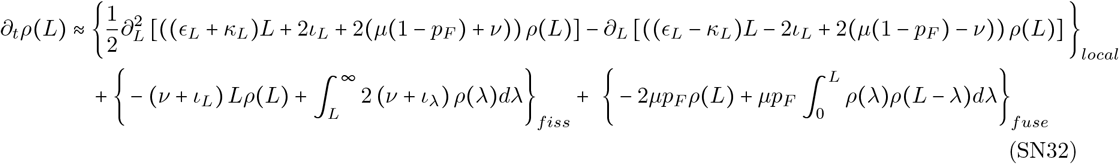

The above expression is again left in terms of general length-dependent rates of expansion, contraction, and insertion and appropriately labeled to differentiate local, fission-, and fusion-based contributions. For completeness, we have reintroduced local contributions from insertion and substitution (these subdominant effects will be dropped again shortly). Setting the time derivative to zero, we find an integral ordinary differential equation (ODE) for the steady state distribution.

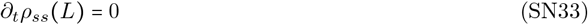

Henceforth, the subscript *ss* (indicating steady state) will be dropped and assumed throughout. We now approximate this condition to focus on the dominant terms driving the distribution at large lengths. Again, substitutions and insertions are subdominant in the local terms due to a length-independent target size. Additionally, *νL* ≫ 2*µp*_*F*_ such that the fusion outflux remains subdominant at long lengths. These approximations yield a slightly simpler steady state condition in the asymptotic *L* ≫ 1 regime.

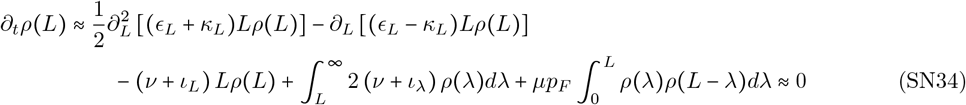

Expressing this in terms of the power-law parameterization in Equation SN1, we find the following.

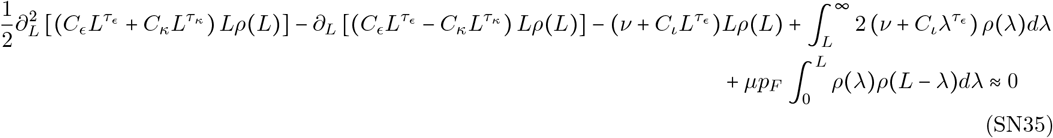

Importantly, no generic closed-form solution to this ODE can be found. First, the complications introduced by fusion are significant, as integration must be performed over the short repeat length regime, limiting our ability to decouple the long length asymptotic dynamics. Second, even when omitting fusion entirely, the remaining terms describe a second-order integral ODE, which can be recast as a third order ODE to remove explicit nonlocal transitions; unfortunately, few third order ODEs are exactly soluble.

The functional form of the fusion term fundamentally limited our ability to further disentangle the dynamics. Motivated by our computational results, we proceeded under the *ansatz* that, for long lengths *L* ≫ 1, the fusion term is everywhere negligible relative to fission and local transitions, the latter providing the largest contributions to the asymptotic dynamics due to length scaling. The validity of this assumption, which we confirmed by exploring the allowed parameter space (for various parameterizations) with our computational model, is likely a consequence of both the low rate associated with *µp*_*F*_ ≪ *νL* and the rapid geometric decay in the short length regime followed by the further (though sub-geometric) monotonic decay at longer lengths. In contrast, Equation SN15 demonstrates the importance of fusion to the short length dynamics, which exactly balance length decreases due to *ν* substitutions. We found non-negligible contributions from fusion only in the substitution-dominated short length regime, consistent with our analytic results for short repeats, and in cases where the distribution is far from steady state. A more principled argument for the relative suppression of fusion at long lengths likely exists, but is unnecessary for the present purposes. For further analysis, we proceed under the following approximation of Equation SN35 for *L* ≫ 1, which retains nonlocal transitions only in the form of repeat fission.

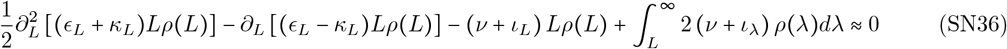

Again using the parameterization in Equation SN1, steady state is maintained under the following approximation.

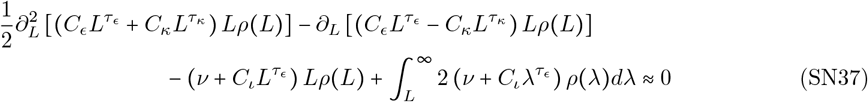

The above expression describes four distinct effects that collectively lead to steady state, parameters permitting: bidirectional diffusion due to net repeat instability from the combined effects of expansion and contraction, expansion-contraction bias generating a directional flux, net (substitution- and/or insertion-based) outflux due to repeat fission, and net nonlocal (substitution- and/or insertion-based) influx due to fission of any longer repeats. Strictly speaking, the outflux due to fission is technically local in the equation (despite representing non-local transitions, appropriately accounted for by the nonlocal influx), as it corresponds to the rate of interruptions of repeats in the focal length class without reference to the subsequent repeat lengths.

#### 4.5 Decomposition of parameter space into dynamical regimes

To better understand the dynamics, we characterized the behavior of the multiplier-coupled power-law model of instability rates in qualitatively distinct parameter regimes as primarily controlled by a sum of two, three, or four of the terms in the more complete steady state approximation shown in Equation SN37. These regimes can be identified by studying the length scaling associated with each term, which indicates the primary difference between local and fission-based contributions to changes in length. In contrast to the generality of Equation SN34 (and, assuming subdominant fusion; Equation SN36), the following decomposition is dependent on the details of both our empirically estimated mutation rates and the parameterization in Equation SN1; differing estimates or parameterizations may result in differences in the quantities of importance to the dynamics, but are unlikely to fundamentally alter the subsequent qualitative conclusions about the steady state distribution.

##### 4.5.1 Asymptotic length dependence of local transition rates and Δ*τ*

First focusing on local length changes, the quantity Δ*τ* (see Equation SN24) provides an indicator variable for the sign of the bias term at asymptotically large lengths. This can be seen by manipulating the length dependence inside the first derivative term representing the directional (i.e., signed) per repeat rates.

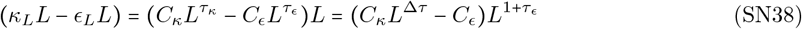

The sign of this term determines the asymptotic dominance of either the expansion or contraction rate (i.e., the bias at long lengths) and is dependent only on Δ*τ* (i.e., the constants *C*_*ϵ*_ and *C*_*κ*_ do not scale with length). For convenience, Δ*τ* ≡ *τ*_*κ*_ − *τ*_*ϵ*_ was defined to correspond to the sign of *κ*_*L*_ − *ϵ*_*L*_ such that positive Δ*τ* indicates asymptotic contraction bias (and stability, explained below).

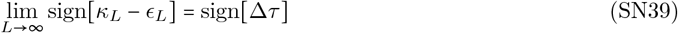

As described in Equation SN23, Δ*τ* (provided Δ*τ* ≠ 0) specifies a length *L*^*^ at which the sign of *κ*_*L*_ − *ϵ*_*L*_ can reverse if the directional flux changes above *L* = 9 (relative to the initially expansion-biased rates at low lengths, *ϵ*_*L*=9_ > *κ*_*L*=9_). The overly simplistic power law parameterization limits the behavior to, at most, one such sign change in the long length regime (i.e., there is no more than one intersection between two monotonically growing power law functions): asymptotic contraction-bias (Δ*τ*≥ 0) requires one sign reversal, while asymptotic expansion bias (Δ*τ* ≤ 0) depicts a consistent directional flux throughout the long length regime (due to the empirically estimated expansion bias at *L* = 8). The asymptotic length dependence of the directional flux coefficient is determined by the scaling of the faster of the two rates (i.e., the larger exponent, *τ*_*ϵ*_ or *τ*_*κ*_ ).

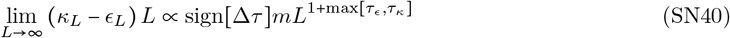

Here, the dependence on *m* is common to the definitions of *C*_*ϵ*_ and *C*_*κ*_ (see discussion below). As can be seen in Equation SN28, the larger of the two exponents also dictates the asymptotic length dependence of the diffusion coefficient in the second derivative term.

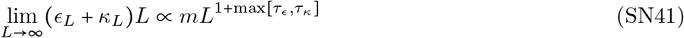

Monotonicity of both instability rates requires *τ*_*ϵ*_, *τ*_*κ*_ ≥ 0. As a result, the sign of Δ*τ* dictates the larger exponent (i.e., when Δ*τ* > 0, max[*τ*_*ϵ*_, *τ*_*κ*_ ] = *τ*_*κ*_ ; when Δ*τ* < 0, max[*τ*_*ϵ*_, *τ*_*κ*_ ] = *τ*_*ϵ*_ ) and thus the asymptotic length scaling for the coefficients of both directional and diffusive local changes in length. Because Δ*τ* adequately characterizes the local dynamics (which, in turn, determine the shape of the DRL at steady state), the parameter space roughly decomposes along lines of constant Δ*τ* . Parameter combinations with the same Δ*τ* result in similar steady state DRLs (see **Fig. S11**), which, in our inference, form a ridge of similar posterior probabilities based on their divergence from the empirical DRL (see **Figs. 5a** and **S8a**).

The linear dependence on *m* is common to the dominant local terms in Equation SN37 (*C*_*ϵ*_, *C*_*κ*_ ∝ *m* in Equation SN1), which describe the collective effects of repeat instability due to expansion and contraction. Unlike Δ*τ, m* contains no information about the relative contributions of expansion, contraction, and insertion (again linear in *m*; *C*_*ι*_ ∝ *m*). However, because substitution-driven transitions do not depend on *m*, this parameter captures a competition between instability-related effects (including fission due to insertion) and fission due to substitution. This generates deviations from lines of constant Δ*τ* that are visible at low *m* values in **Figs. 5a** and **S8a**. At low *m* and slower growth rates (*τ*_*ϵ*_, *τ*_*κ*_ ≲ 1), substitutions play a more substantial role (further discussed below).

##### 4.5.2 Relative strength of substitution- and insertion-driven fission and *L*_fis_

In contrast to the local dynamics, repeat fission is asymptotically dominated by insertion, which outcompetes the substitution rate at asymptotically long lengths. A characteristic transition length *L*_fis_ emerges from the multiplier-coupled model, above which the fission rate is dominated by insertion-based interruptions. The fission-transition length can be found by comparing the per-target rates of substitution *ν* and insertion 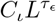.

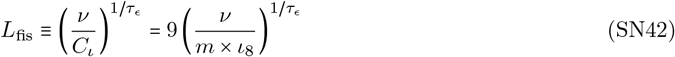

Here, definition of *C*_*ι*_ was used to show the explicit dependence on *m*. From our empirical estimates, *ν*/ *ι*_8_ ≈ 40 for mononucleotide *A*-repeats (see **Fig. 3b**); thus, for values of *m* ≲ 40 (which includes all computationally explored parameters), the ratio being exponentiated is greater than one. Consequently, as *τ*_*ϵ*_ increases, *L*_fis_ decreases, eventually exiting the long length regime (i.e., *L*_fis_ ∼ 9 for *τ*_*ϵ*_ ≫ 1) such that, for large *τ*_*ϵ*_, all fission events in long repeats are insertion-dominated. Substitutions dominate more of the long length regime when both *τ*_*ϵ*_ and *m* are small (i.e., for *τ*_*ϵ*_ < 1 and *m* ≪ 50), while substitutions are subdominant at more long lengths when *τ*_*ϵ*_ and *m* are large.

The quantities *τ*_*ϵ*_ and *m* together characterize the asymptotic length dependence of repeat fission by determining the value of *L*_fis_, the length at which a switch occurs from substitution- to insertion-dominated fission (at lengths *L* < *L*_fis_ and *L* > *L*_fis_, respectively). In the parameter regime where fission is entirely insertion-dominated at long length (i.e., *L*_fis_ ∼ 10 when *m, τ*_*ϵ*_, or both parameters are sufficiently large), all relevant terms in Equation SN37 are linear in *m* such that the dynamics are independent of this parameter; in this case, the relative importance of local transitions and fission can be characterized by the values of *τ*_*ϵ*_ and Δ*τ* (or, equivalently, *τ*_*ϵ*_ and *τ*_*κ*_). In this extreme, in addition to the decoupling between short and long repeat dynamics, long repeats are predominantly subject to insertion- and deletion-based mutations (i.e., repeat instability-generated expansions, contractions, and insertions) and evolve *mechanistically* independently (i.e., change length predominantly due to distinct mutational processes) from short repeats; importantly, this claim implicitly requires that long repeat dynamics are independent of *µ* (i.e., fusion remains negligible). In the opposite extreme, where insertions remain irrelevant at all populated lengths, the dynamics are independent of *τ*_*ϵ*_ and can be represented in the space of (Δ*τ, m*).

In Section 6 this supplemental note, we assess the range of plausible values of *L*_fis_ suggested by the results of our Bayesian inference procedure.

##### 4.5.3 Distinguishable dynamical regimes

The sign of Δ*τ* decomposes the space into two primary regions and the boundary between them, only one of which evolves towards a recognizable steady-state equilibrium on reasonable timescales. The repeat length distribution only stabilizes for the subset of the parameter space with Δ*τ* ≥ 0, which we refer to as the (asymptotically) *contraction-biased* regime. Expansion-biased directional flux (Δ*τ* < 0) leads to indefinite repeat expansion, exceeding the net shortening effects of contraction and repeat fission (the boundary at Δ*τ* = 0 is discussed below). In contrast, asymptotic contraction bias, which is directionally consistent with fission, truncates the distribution at finite lengths, leading to a stable distribution (consistent with our empirical observations). The collection of stable parameter combinations can be further decomposed based on the extent to which contraction dominates over expansion (i.e., the value of Δ*τ* ); this is the primary subject of our subsequent analysis. Within each subregime of Δ*τ* > 0, the full set of changes in repeat length in Equation SN32 that lead to a stable distribution can be reduced to a subset of effects that approximate the dominant contributions that shape the distribution. Parameter combinations with Δ*τ* ≫ 1 are controlled largely by a local balance between diffusion and directional flux (contraction-biased at most or all lengths in the *L* ≫ 1 regime), both of which dominate over fission. Intermediate values 1 ≳ Δ*τ* > 0.6 show a balance between local transitions and an outflux due to fission (i.e., the non-integral terms representing fission in Equation SN37). Weak contraction bias (very roughly Δ*τ* ≲ 0.5) requires a full accounting of local effects and both the influx and outflux due to fission (i.e., the full set of terms in Equation SN37).

At all points in the stable regime, the net increase in repeat length due to expansion (ignoring subdominant *µ* contributions) is counteracted by the combined effects of contraction (which exceeds expansion above some length *L*^*^) and fission due to insertions and/or substitutions. Based on our computational analyses, fusion remains far too infrequent to stabilize long repeats (justifying the approximation in Equation SN37). This regime is bounded by Δ*τ* = 0, but we note that very low positive values of Δ*τ* have expansion-dominated rates across an extended range of lengths (i.e., *L*^*^ → ∞ as Δ*τ* → 0; see Equation SN23), truncating the distribution at unrealistically high *L*. These parameters (i.e., small, positive Δ*τ* ) are inconsistent with the empirical distribution.

The remaining set of parameter combinations fall into two categories, both of which are implausible with respect to the collection of empirical observations presented in this manuscript: those with very low instability rates (i.e., very slowly evolving distributions that do not equilibrate on evolutionarily-relevant timescales) and those with unstable dynamics (i.e., rapid aggregation of repeats and explosive growth in genome size). Exceedingly slow evolution occurs if *m* is order one and both exponents approach zero (i.e., *τ*_*ϵ*_, *τ*_*κ*_ < 1 such that *τ*_*ϵ*_ and ∥Δ*τ*∥ both remain small). These parameter combinations depict repeat instability rates that grow much more slowly with length than our empirical rate estimates (see **Fig. 3b**); as a result, this regime is entirely ruled out under the informative prior (indicating inconsistency with rate estimates based on popSTR [1]; see **Fig. 5a**). We refer to this region of parameter space as the *slowly evolving* regime, as the repeat instability rates depicted remain scantly above the estimated substitution rates, even for the longest well-populated lengths in the human genome.

Parameter combinations that are dynamically unstable (referred to as the *unstable* regime) correspond to universally expansion-biased dynamics (Δ*τ <* 0, excluding slowly evolving parameters that may or may not equilibrate). This regime comprises the majority of the parameter space explored in our computational model, which can be dynamically disallowed simply by requiring steady state based on our empirical findings. These parameters represent a non-equilibrium dynamical regime subject to strong nonlinear effects that result in increasingly rapid changes in the repeat length distribution. In the multiplier-coupled model, parameters with Δ*τ =* 0 are similarly unstable due to the universal expansion-bias at all lengths, which is inherited from the empirically observed bias at *L* = 8 (i.e., *κ*_9_ − *ϵ*_9_ = *m*(*κ*_8_ − *ϵ*_8_) > 0) and subsequent parallel growth of expansion and contraction.

Last, we note that the decomposition of the dynamics is dramatically simplified for mid-to-large multipliers *m* ≳ 4. We focus our discussion of the analytics on this case, where the dynamics can be characterized largely by Δ*τ* alone (see, e.g., **Fig. 5a**). However, the same dynamics apply to smaller *m*, values with the additional complication that the balance between substitutions and insertions breaks the dynamical similarity along lines of constant Δ*τ* (this can be seen as curvature along posterior ridges in **Fig. 5a** due to more explicit dependence on *τ*_*ϵ*_ near the origin).

#### 4.6 Unstable dynamics in the asymptotically expansion-biased regime Δ*τ* ≤ 0

We briefly discuss the dynamics of the unstable regime, as it informs our considerations for more realistic parameter combinations. There are two distinguishable cases where steady state cannot be assumed, contrary to the observation of long term maintenance of the DRL across the primate phylogeny. First, the expansion rate, which is initially dominant at *L* = 9 (i.e., *ϵ*_9_ > *κ*_9_), may have a length dependence that rapidly outcompetes that of contraction, resulting in increasingly larger expansion-bias with increasing length. This corresponds to Δ*τ <* 0 with large magnitudes ∥Δ*τ*∥ ≫ 1. In the second case, which occurs for relatively small values of ∥Δ*τ* ∥ approaching Δ*τ* = 0 (i.e., when *τ*_*ϵ*_ = *τ*_*κ*_) the length dependence of expansion increases with length comparably to, or slightly in excess of, the length-dependence of the contraction rate. In this case, the directional flux is minimized such that the relative importance of repeat fission becomes inflated. In the former case with highly dominant expansion, the bias generates a large directional flux that rapidly increases repeat lengths. In this regime, the flux is well approximated by the rate 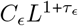 (i.e., *τ*_*κ*_ is negligible at all lengths *L* > 10) and the rate of changes in length rapidly accelerates with increasing repeat length. This nonlinearity results in an indefinitely extending tail, some of which feeds back into lower length classes due to fission, as large *τ*_*ϵ*_ also generates large rates of insertion-based repeat fission 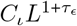; however, the coefficient *C*_*ι*_ is roughly two orders of magnitude smaller than *C*_*ϵ*_ such that fission alone is unable to counteract expansion alone at any length. Repeat fission additionally increases the mass of the distribution, as it does not conserve repeat number (one repeat is replaced with two shorter length repeats). These shorter repeats are then subject to the large directional push due to expansion, which further increases the weight in the distribution tail. This feedback loop generates a rapid, indefinitely growing genome, which reshapes the distribution. Rapid fission of any given repeat in the long tail equiprobably adds mass to the distribution in all shorter length classes, an integrated effect that is increased with increasing mass above a given length. This eventually leads to an extreme influx into the shortest length classes, inherently coupling the dynamics in the short and long length regimes (i.e., violating our assumption of separability and distorting the substitution-based geometric distribution). In addition to the unstable dynamics, the indefinite extension of the distribution tail quickly results in highly relevant expansion probabilities approaching one (because power-law rates do not saturate), which simultaneously makes computational modeling impractical across this regime and will invariably result in characteristic changes to the shape of the distribution when the power law parameterization is corrected to saturate at this probabilistic bound.

In the second case, for Δ*τ* near zero (still assuming *τ*_*ϵ*_, *τ*_*κ*_ > 1 to avoid the slowly evolving regime), the directional flux is again expansion dominated and non-vanishing. Here, the relative importance of fission is inflated, which limits the rate at which the tail extends. However, the difference between the expansion and contraction rates, even at *L =* 8, is substantially in excess of the insertion rate. This leads to the inability of repeat fission to independently counteract the directional flux at all lengths.

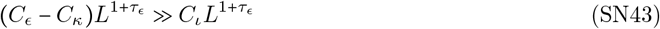

In this sense, contraction must be sufficiently large to mitigate expansion in order for the distribution to truncate at finite length. This only occurs when Δ*τ* > 0 such that the contraction rate approaches, and eventually exceeds, the expansion rate (though this may occur at an extremely large length for the smallest values of Δ*τ >* 0). This defines a bound for the contraction-biased regime that leads to steady state. A small, non-negligible contribution from fission is relevant above this bound, but is insufficient to control the directional push from expansion for very low ∥Δ*τ*∥. We note that this effort may be aided by substitution-based fission, but the associated length scaling (i.e., the fission rate *νL*) only becomes relevant for small multipliers *m* when *τ*_*ϵ*_ ≲ 1 (i.e., when 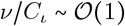 and 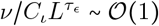 across relevant lengths); parameters in this range correspond to sufficiently small rates that evolution proceeds exceedingly slowly, as discussed above. Relative to the directional flux, diffusive changes in length are also magnified when ∣Δ*τ*∣ is small, but do not stabilize the distribution.

#### 4.7 Stable dynamics in the asymptotically contraction-biased regime Δ*τ >* 0

Given the similarity between primate DRLs (**Fig. 1b**), realistic parameter combinations in the multiplier-coupled model sit within the subset that evolve towards stable, steady-state distributions. In contrast to the unstable regime, parameter combinations with asymptotic contraction bias Δ*τ >* 0 result in a distribution that approaches a stable steady state. Perhaps more intuitively, this regime can be equivalently characterized by *L*^*^, the length at which expansion and contraction rates are equal and the directional flux vanishes. For all values Δ*τ* > 0, evaluation of Equation SN23 shows that *L*^*^ ≥ 9 (and *L*^*^ ≥ 10 for realistic values of Δ*τ* ) such that this intersection sits within, or near the boundary of, the long length regime. As the combination of our empirical estimates at *L* = 8 and the multiplier-coupled model (namely, *m* > 0) dictate that the expansion rate exceeds the contraction rate at length *L* = 9, an intersection between the rates must occur prior to the asymptotic dominance of contraction. Repeat lengths below *L*^*^ are necessarily expansion-biased and those above *L*^∗^ are contraction-biased, with no directional flux at *L*^∗^. For large values of Δ*τ*, the value of *L*^*^ approaches a length close to *L* = 10. For very small Δ*τ*, this can sit at very large lengths, approaching *L*^*^ → ∞ as Δ*τ* → 0. At many values of Δ*τ* > 0, *L*^*^ occurs well within the tail of well-populated lengths in the whole-genome human DRL.

The transition between expansion- and contraction-biased lengths controls and complicates the dynamics. The rates of expansion and contraction can remain within the same order of magnitude over an extended range when Δ*τ* is small, as the approach to *L*^*^ is slow from either side. This can result in a first derivative term (i.e., the directional flux) that remains relatively small over lengths spanning part or all of the distribution tail. In this case, the diffusion term, which arises as a subdominant correction to the discrete local behavior (see Section 4.1.2) plays an important role in the extended neighborhood of *L*^*^ . For the smallest values of Δ*τ* ≪ 1, the intersection at *L* occurs at extreme lengths above those populated in the empirical distribution. This results in a dramatically extended tail due to a wide range of expansion-biased lengths before contraction bias becomes appreciable. Additionally, this generates a dramatically larger genome (with an excess of very long repeats) inconsistent with the observed range of mammalian genome sizes. This is somewhat similar to the unstable dynamics described above, but is eventually counteracted by sufficiently large contraction rates that truncate the distribution and stabilize the (presumably unrealistic) shape of the distribution.

As mentioned above, the dynamics of long repeats across the space of parameters with Δ*τ >* 0 can be approximated by Equation SN37 under the assumption that repeat fusion is sufficiently infrequent. However, the dynamics reduce further in parts of this regime where fission (influx and, to a lesser extent, outflux) can be treated as negligible. While only appropriate for some parameter combinations, such approximations can be useful because they eliminate the need to explicitly treat the nonlocal effects of fission influx, which reduces the second-order integro-differential equation to a second-order ODE.

##### 4.7.1 Strong asymptotic contraction bias

We first attempted to describe the dynamics in the regime with sufficiently large Δ*τ* such that the majority of long repeats have contraction-biased rates (i.e., *L*^*^ ∼ 10 occurs immediately above the short repeat regime). In this case, the diffusion term remains relevant because *L*^*^ lies in the long length regime but the directional flux becomes increasingly relevant with increasing length, an effect magnified by necessarily larger Δ*τ* . For most contraction-biased parameter combinations, the dynamics of Equation SN37 are well approximated by the following.

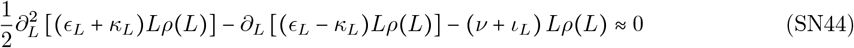

In terms of the parameterization in Equation SN1, this becomes the following.

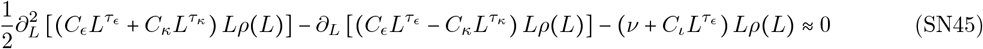

Here, we have treated the integral terms describing the influx due to fission as subdominant. These contributions are outcompeted by each of the remaining rates, which scale asymptotically in length with larger exponents (i.e., as 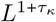 or 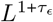).

##### 4.7.2 Strictly local approximation for strong asymptotic contraction bias Δ*τ* ≫ 1

In the regime of largest Δ*τ*, the outflux due to fission is negligible, as well; the asymptotic scalings of the directional flux and diffusion coefficients 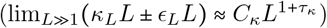 outcompetes the fission outflux at very large lengths 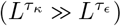. This defines a dynamical sub-regime within the space of contraction-biased parameter values, with dynamics well approximated by the following balance.

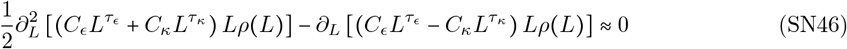

In this regime, sufficiently rapid decay in *ρ L* results in a net negative directional flux (the correct sign associated with contraction bias) that counteracts the strictly positive diffusion term. For example, when *m* = 4, this expression provides a good approximation to the dynamics primarily for Δ*τ* > 1.5 (see **Fig. SN1**) and breaks down as fission becomes increasingly relevant for weaker contraction bias (for larger multipliers, this regime breaks down at lower Δ*τ* ; e.g., for *m* = 16, Equation SN46 remains a good approximation for Δ*τ* > 1; see **Fig. SN6**). In this regime, *L* occurs at or adjacent to *L* = 10 such that nearly all long lengths are contraction-biased. For example, the parameter combination (*m, τ*_*ϵ*_, *τ*_*κ*_) = (4, 0, 2) results in *L*^*^ ≈ 11 and a contraction rate of roughly 2.5 times the expansion rate at *L* = 18, while for (*m, τ*_*ϵ*_, *τ*_*κ*_) = (4, 0, 4), *L*^*^ ≈ 12 and the contraction rate is roughly tenfold the expansion rate at *L =* 18. In the more extreme case of Δ*τ =* 4, we can better understand the truncation of the distribution by further approximating Equation SN46 under the very rough assumption that the asymptotic dependence is immediately relevant for lengths *L > L*^*^ .

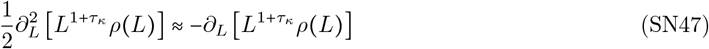

Noting that the contraction rate constant *C*_*κ*_ cancels such that this equation is dependent only on the exponent *τ*_*κ*_, an approximation for the asymptotic shape of the steady state distribution can be obtained in closed form (valid only for lengths *L* ≫*L*^*^ ).

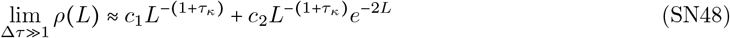

To evaluate the accuracy of this rough approximation, the arbitrary constants *c*_1_ and *c*_2_ can be found using values from our computational model at two lengths *L*_1_ and *L*_12_ (where *L*_1_, *L*_2_ ≫ *L*^*^ ) to constrain the distribution at *ρ*(*L*_1_) and *ρ*(*L*_2_). Comparing this expression to numerical solutions, we found reasonable agreement for the most extreme values of Δ*τ* > 0 (e.g., Δ*τ* = 3−4 for *m* > 4, the largest Δ*τ* with computational modeled DRLs) at the longest lengths *L* > *L*^*^ in the distribution and an expected departure as *L* → *L*^*^ .

#### 4.8 Intermediate asymptotic contraction bias

Intermediate values of Δ*τ* > 0 (e.g., roughly 1.5 > Δ*τ* ≳ 0.7 for *m* = 4; see **Fig. SN1**) require a description of the outflux due to fission that appears in Equation SN45. At the larger end of this range (e.g., Δ*τ* ∼ 2), the numerical solution to Equation SN46, which omits all effects of fission, approximates the asymptotic shape of the distribution for lengths *L L*^*^ . This indicates that the impact of outflux due to fission is primarily localized to intermediate lengths *L*^*^ > *L* > 10 and therefore most relevant when the rate of expansion exceeds contraction. At the same time, this suggests that the truncation of the distribution is driven by the contraction-biased directional flux, rather than fission alone or due to the combined effects of fission and contraction.

#### 4.9 Weak asymptotic contraction bias

For smaller values of Δ*τ* that approach *τ*_*ϵ*_ = *τ*_*κ*_ (e.g., roughly Δ*τ* < 0.7 for *m* = 4; see **Fig. SN1**), the dynamics revert to Equation SN37, which omits only fusion. The resulting distributions stabilize, in part, due to the nonlocal influx from fission of longer repeats. For these parameter combinations, *L*^∗^ sits in the middle of the long length tail; for lengths *L* > *L*^*^, the distribution is well-approximated by solutions to Equation SN45. This is consistent with a nonlocal net flow from lengths *L* > *L*^*^ to lengths *L*^*^ > *L* > 10 and indicates that the dynamically relevant effects of fission influx are primarily localized to the latter. The net effects of fission alone (i.e., fission influx minus outflux) result in a net loss of long repeats, with little gain from any existing longer length repeats, and a compensatory net gain of intermediate length repeats within the distribution tail (exceeding those lost to the short length regime). With decreasing values of Δ*τ* → 0, *L*^*^ approaches very large values, extending the range of lengths that receive a net influx. All effects represented in SN37 are thus required to adequately approximate the dynamics that lead to a steady state distribution when Δ*τ <* 1 (and a potentially broader range, depending on *m*).

The nonlocal integral dependence in Equation SN37 that describes the influx due to fission complicates the steady state condition in this regime. To find solutions, we re-expressed the second-order integro-differential equation as a third order differential equation by applying an overall length derivative to each term.

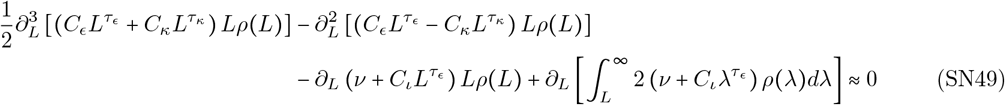

Taking this derivative allows us to apply the fundamental rule of calculus to replace the derivative the integral with the integrand evaluated at the integration limits. Under the assumption that the distribution decays sufficiently rapidly such that 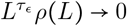 as *L*→ ∞, the integral term becomes the following.

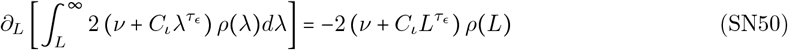

This allows us to re-expresses the second-order integro-differential equation as the following third order ODE.

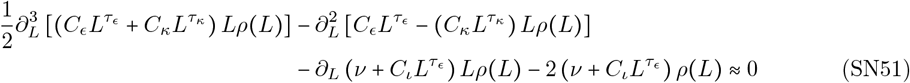

After applying a length derivative to the steady state condition *∂*_*L*_ (*∂*_*t*_*ρ* (*L*) = 0, this now corresponds to a constraint on *∂*_*L*_*ρ*(*L*), the flux through each length *L*. This can be seen by swapping the order of the length and time derivatives (i.e., *∂*_*t*_ (*∂*_*L*_*ρ(L*)) = 0), dictating that the various fluxes through each length must sum to a time-independent constant *ϕ*_*L*_. In the special case where *ϕ*_*L*_ = 0, this corresponds to a steady state condition that maintains the shape of the distribution in equilibrium. We confirmed via our computational model that, once steady state was reached, the net flux through each individual bin independently vanished (see **Figs. SN2-SN4**), indicating that the equilibrated state is maintained in a detailed balance. Insofar as our approximations remain valid, Equation SN51 provides a local expression for the steady-state flux through the repeat length distribution in the large length regime; this includes the nonlocal contributions of repeat fission to the flux, represented by boundary effects at length *L* (more accurately, at length *L* + 2 ≈ *L*). While this equation cannot be solved analytically, numerical solutions can be readily obtained.

Assuming the effects of fusion remain subdominant in the long length regime, Equation SN51 captures the full set of relevant dynamics associated with repeat length transitions. Solutions to this equation, obtained after applying the additional constraint that the fluxes vanish, are applicable across the full range of parameters that evolve towards steady state distributions Δ*τ >* 0. In contrast, Equations SN45 and SN46 are approximations to these dynamics appropriate in a subset of parameter space (very roughly, when 2.5 > Δ*τ >* 1 and when Δ*τ <* 1, respectively). However, in addition to aiding in our intuitive understanding of the dynamics, the absence of the third derivative in the latter equations makes numerical solutions more reliable, as they are less susceptible to instabilities in numerical techniques; this facilitates a slightly more reliable comparison between the numerical solutions and results of our computational model.

#### 4.10 Obtaining numerical solutions to the steady state dynamics for Δ*τ <* 0

To compare our analytic understanding of the dynamics to the results of our computational model for generic parameters, we resorted to solving Equations SN45, SN46, and SN51 numerically. All numerical solutions were obtained using the NDSolve function in Mathematica 14.0 [2]. Solutions to second-order differential equations require the specification of two additional constraints that together fix the normalization constant and the linear coefficient that determines the relative weight of the two real solutions to the equation, if both exist. The third order equation for constant flux requires a third constraint that ensures vanishing flux.

For the second-order equations, we chose to constrain the values of *ρ*(*L*) at two lengths, *L*_1_ and *L*_2_, using the results of our computational model (i.e., *ρ* (*L*_1_) = *ρ*_*sim*_ (*L*_1_) and *ρ* (*L*_2_) = *ρ*_*sim*_ (*L*_2_), where *ρ*_*sim*_ (*L*) is the value of the computationally propagated (i.e., ‘simulated’) distribution at length *L* once it has reached steady state). The choice of *L*_1_ and *L*_2_ is somewhat arbitrary, provided they are both in the long length regime where the continuum approximation is valid and that any additional constraints on the regime of validity are respected (e.g., Δ*τ >* 0 and sufficiently far from Δ*τ* = 0, *τ*_*ϵ*_, *τ*_*κ*_ ≥ 0, etc.). For the majority of comparisons, we chose two lengths that are well defined for any parameter combination: *L*_1_ = *L*\* to the nearest integer value) and *L*_2_ = *L*_trunc_, where *L*_trunc_ is the length bin for which the occupancy of (rounded the non-normalized distribution first drops below a single count (i.e., *L*_trunc_ = min [*L* for *ρ*(*L*) < 1]) and represents the truncation point of the distribution. *L*_trunc_ is uniquely defined because of two properties: *1*. all computationally modeled distributions decayed monotonically, and *2*. we propagated the expected DRL under a mean field approximation, omitting stochastic effects. All comparisons were made using non-normalized DRLs to identify the truncation point of the distribution and obtain *L*_trunc_.

For solutions of Equation SN51, we chose to apply the third constraint by fixing the value of the DRL at *L*_3_ = *L*_2_ − 1 for convenience. We note that inappropriate choice of *L*_3_ outside of the regime of validity of the approximation can result in numerical instability in solutions to the third order ODE (e.g., constraining the solution using a length class *ρ*_*sim*_ *L*_3_ that has not yet equilibrated). We chose not to use the intuitive lower bound of the long length regime at *L* = 10 to identify any potential effects associated with the breakdown of the continuum approximation.

For any parameter combinations with *L*^∗^ < 10 (i.e., non-equilibrium cases with Δ*τ <* 0) we chose a lower bound at *L*_1_ = 10 to avoid values in the short length regime; however, unstable dynamics were compared to numerical solutions only to confirm significant departure from any steady state characterized by Equation SN51 and to identify any common features that emerged.

Finally, for all comparisons in the long length regime, the value of *ν* was replaced with *ν*_*fission*_, the appropriate rate estimated from the three-unit context *AAA* → *ABA* in which substitutions *A* → *B* result in repeat fission. Mutation rates *µ* and *ν* that appear in Equation SN10 were replaced with distinctly estimated rates relevant for the three-unit contexts associated with local transitions due to substitutions defined by *µ*_*local*_ (i.e., the summed rates of *ABB* → *AAB* and *BBA* → *BAA* substitutions) and *ν*_*local*_ (the summed rates of *AAB* → *ABB* and *BAA* → *BBA* substitutions), respectively.

### 5 Comparison between numerical solutions and computationally modeled distributions

As described above, numerical solutions to Equations SN45, SN46, and SN51 were compared across the space of parameter combinations that led to steady state distributions. Because our computational model obtained results over a large, but finite number of iterations, parameters corresponding to insufficiently high mutation rates failed to equilibrate in the allotted time (i.e, in a number of iterations corresponding to at least 10^9^ generations of evolution, after accounting for a factor that progressively rescales time to increase computational speed). These slowly evolving computational results were localized to the lowest values of (*τ*_*ϵ*_, *τ*_*κ*_) for the lowest multipliers *m* ≤ 2.5 (i.e., points closest to the origin of the (*τ*_*ϵ*_, *τ*_*κ*_) plane for *m* ≤ 2.5; for larger values of *m* ≥ 4, all parameters equilibrated sufficiently quickly) and spanned a larger range of parameter values for smaller *m*. For *m* = 2.5, this roughly corresponds to parameter values *τ*_*ϵ*_,*τ*_*κ*_ ≲ 1. In this parameter range, the ridge of roughly equivalent posterior values (calculated by comparing computationally modeled distributions at the final time point to the empirical distribution) deviates from lines of constant Δ*τ* . This corresponds to the point at which substitution becomes non-negligible and substitution-based fission occurs at a rate comparable to or greater than insertion-based fission. Given indefinite time to evolve, such parameter combinations likely equilibrate, provided the combined action of contraction, substitution, and insertion is sufficient to truncate the distribution at finite length. As no equilibrium was reached, these points were excluded from our comparison to numerically produced steady-state distributions. Additionally, all such parameters were excluded from the 95% HDR of the posterior when using an informative prior (see **Fig. 5a**).

For all points outside of this slowly evolving region, equilibrated steady state distributions are quantitatively similar along lines of constant Δ*τ* . The following plots show comparisons at points along an anti-diagonal line perpendicular to Δ*τ* = 0 defined by *τ*_*ϵ*_ + *τ*_*κ*_ = 3.5; for comparison, we included parameter combinations with exponents *τ*_*ϵ*_ = *τ*_*κ*_ = 1.7 (our discretization scheme did not allow for equal exponents that sum to 3.5). Parameter combinations on this line are representative of the full set of computationally modeled Δ*τ* outside of the slowly evolving region and could be obtained for all considered multipliers. Along the line of *τ*_*ϵ*_ + *τ*_*κ*_ = 3.5 (after adding a point with Δ*τ* = 0 and *τ*_*ϵ*_ + *τ*_*κ*_ = 3.4), we selected examples that span the qualitatively distinct behaviors across the space of Δ*τ >* 0 for values Δ*τ* = {0, 0.1, 0.3, 0.5, 0.7, 1.1, 1.5, 2.5, 3.5} . The first two points were included for completeness, as they show examples of computationally modeled distributions that are not expected to be well described by numerical solutions. For Δ*τ =* 0.1 the location of *L*^*^ ≈ 942, which is substantially longer than the maximum length included in our computational model at *L*_*boundary*_ = 200; a reflective boundary condition was imposed at *L = L*_*boundary*_ to simultaneously prevent excessively mutation rates that preclude further computational iteration and to identify parameter combinations that result in excessively large genome size when truncation occurs at unrealistically large lengths *L*_trunc_ ≫ 200 (e.g., for Δ*τ =* 0.1). The point at Δ*τ =* 0 was included as an example of unstable dynamics; the computationally modeled distribution equilibrates to the boundary condition at *L*_*boundary*_ (note the balanced fluxes in the largest lengths for computational results with Δ*τ =* 0 in Figs. SN2i and SN3i), resulting in an artifactual shape (i.e., the non-monotonicity in computationally modeled curves shown in Figs. SN1i, SN4i-SN6i).

#### 5.0.1 Comparisons of dynamical approximations and steady-state distributions

**Fig. SN1** compares computationally modeled distributions for these example parameter combinations (for *m =* 4) to the geometric distribution that describes the short length regime (Equation SN10 using values the average single-unit context rates *µ* = *µ*_*B*→*A*_ and *ν* = *µ*_*A*→*B*_) and to the three nested approximations of the long length tail of the distribution obtained by numerically solving Equation SN45, SN46, and SN51. These plots contrast numerical solutions produced under the full set of dynamics (excluding fusion), in the absence of influx due to fission, and in the absence of fission entirely. By observing the lengths at which each successive approximation breaks down, we localized the incoming and outgoing flux due to fission in length space.

For the same parameter combinations, **Fig. SN2** shows the total influx and outflux for each length associated with the individual effects of expansion, contraction, substitution-based fission, and insertion-based fission (as well as fusion and and local transitions due to substitutions). The total flux was separately normalized for each bin (for visualization purposes, as the true magnitudes differ dramatically); bins with equal incoming and outgoing net flux have reached equilibrium (i.e., are maintained in a detailed balance). Slight deviation from this equilibrium occurred in every run, indicating a true steady state distribution was not yet reached (as expected in finite time). However, for all results of interest, the magnitude of deviation is very small. Some parameters with modest values of Δ*τ* showed slight deviation from equilibrium at L=1 characteristic of disequilibrium between the *A* and *B* distributions (i.e., a source of new *A* counts at the *L* = 1 boundary) but showed detailed balance across all remaining classes. This is in contrast to unstable parameter combinations, which harbor large deviations from equilibrium in many or most length classes; at the same time, non-equilibrium simulations rapidly populated the largest length classes, subsequently equilibrating to the artificially imposed boundary condition at the large *L* boundary of the computationally modeled grid.

**Fig. SN3** provides an alternative characterization of the flux for each bin: the incoming and outgoing fluxes for each mutational effect were plotted separately, rather than computing their net (substitutions were separated into local and nonlocal contributions to demonstrate subdominance of the former relative to expansion and contraction). When separating into directional fluxes, the dominance of expansion and contraction over all other fluxes in both directions is made clear; the much higher rates of repeat instability dominate local fluxes throughout the long length regime. This also leads to large-scale diffusion, which captures the significant bidirectional flux that occurs for both expansions and contractions.

**Fig. SN4** shows a direct comparison between the computationally modeled sum of all fluxes (including fusion) at each length for comparison to three approximations of the full dynamics: local dynamics alone, local dynamics and fission outflux (treating influx as negligible), and local dynamics along with a full model of fission (all transitions other than fusion). The accuracy of each approximation can be seen at each length bin via the overlap with the net flux under the full model. In contrast to **Fig. SN1**, which compares numerical solutions that approximate the steady-state distributions, these plots directly compare components of the finite difference equation (and, in a continuum approximation, the differential equations in steady state) for each model specified by Equations SN37, SN45, and SN46. In particular, the nonlocal interactions in Equation SN37 are accounted for directly, without requiring the intermediate step that leads to Equation SN51. This provides a complementary set of comparisons that lead to the same qualitative observations about the regime of validity and accuracy of each approximation across the parameter space, while retaining length-dependent information about the role of each effect across the long repeat regime. Additionally, this provides further justification for the assumption that repeat fusion remains negligible for long repeat dynamics, despite it’s qualitative importance to short repeat dynamics.

Figs. SN5 and SN6 show comparisons between numerical and computational results for the same values of *τ*_*ϵ*_, *τ*_*κ*_, and Δ*τ* shown in **Fig. SN1**, but with multipliers of *m* = 1 and *m* = 16, respectively. For intuition about the effect of the dynamics on the genome size, all comparisons below are shown for non-normalized distributions. The total genome-wide target for *A* bases, 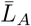, corresponds to the weighted mean of the non-normalized distribution: 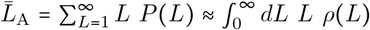. The total genome size is the sum of 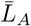 and the corresponding target for *B* bases 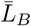. Comparing the same values of Δ*τ* across multipliers *m*, we found qualitative consistency of the decomposition into three dynamical regimes but with boundaries that quantitatively depend on *m*. When *m* is low, the approximations represented by numerical solutions to Equation SN45 (along with the rough analytic solution) and Equation SN46 break down at larger values of Δ*τ* (relative to the same approximations at larger *m*) because substitutions become more relevant, increasing the relative importance of fissions. For example, in **Fig. SN5** for *m* = 1, the purely local approximation in Equation SN45 breaks down at or above Δ*τ* ∼ 2, rather than around Δ*τ* ∼ 1. for *m* = 4 (**Fig. SN1**) and around Δ*τ* ∼0.5 for *m* = 16 (**Fig. SN6**). Here, the relative rate of substitution is closer to the insertion rate and the total rate of fission is closer to the expansion and contraction rates. The relative increase in the strength of fission implies that fission plays a more substantial role in the maintenance of the steady state distribution. In contrast to the more aggressive approximations, numerical solutions to Equation SN51 (i.e., the continuum model with all effects except fusion) remain accurate across the *τ*_*ϵ*_ + *τ*_*κ*_ = 3 line, even at low values of *m*.

**Fig. SN7** shows comparisons between the computationally modeled distribution, numerical solutions, and the closed form approximation in Equation SN48 for parameter combination within and outside of the regime of validity of the latter (i.e., only appropriate when Δ*τ* ≫ 1). Parameter combinations are shown for Δ*τ* = {2, 3, 4} and *m* = {1, 4, 16}. This rough approximation captures the asymptotic falloff of the distribution only when Δ*τ* and *m* are sufficiently large (very roughly Δ*τ* ≳ 3 − 4 when *m* > 4), failing to characterize the shape at lengths closest to the lower boundary of the long length regime around *L* = 10.

## 6 Estimation of *L*\*and *L*_fis_ from the Bayesian posterior distribution

Using the results of our Bayesian inference, we estimated *L* and *L*_fis_, parameters that emerged from our analysis of the repeat length dynamics at steady state. Under the informative prior, the subset of parameter values within the 95% HDR of the posterior lies along a narrow ridge of Δ*τ* values (henceforth, *max posterior ridge*) with appreciable exponents *τ*_*ϵ*_, *τ*_*κ*_> 2 and spanning multipliers *m* = {2.5, 4, 6.4}. This subset of realistic parameter combinations is localized at Δ*τ* = 0.5, rather than at specific values of the parameters *τ*_*ϵ*_ and *τ* _*κ*_ (at least within grid with step size of 0.1 for *τ*_*ϵ*_, *τ*_*κ*_). Using Equation SN23, these parameter values suggest a transition from expansion- to contraction-biased dynamics occurs at *L*^*^ = 22.8 (this is extended to a range of *L* ≈ 20 − 30 for the 99.7% HDR). This occurs at intermediate lengths within the long length regime, well below the truncation point of the DRL (for the T2T genome assembly, *L*_trunc_ = 65). Posterior values under the uninformed (uniform) prior suggest a similar values for Δ*τ* and thus *L*^*^, despite a larger spread in the 95% HDR; this suggests that the rough estimation of *L*\* does not require instability rate estimates at intermediate lengths (e.g., those estimated from popSTR data [1]) but can instead be informed by properties of the DRL (e.g., onset length for repeat instability, truncation length, etc.).

In addition to constraining *L*, our inference results can be used to estimate the range of probable values of *L*_fis_, representing the transition from substitution-dominated fission to insertion-dominated fission. Under the informative prior, the maximum posterior of (*m, τ*_*ϵ*_, *τ*_*κ*_) = (4, 3.1, 3.6) has a corresponding fission-transition at *L*_fis_ = 18.9; the 95% HDR of the posterior spans a range of *m* = 2.5 − 6.4 and *τ*_*ϵ*_ = 2 − 3.5, which corresponds to a tight range of *L*_fis_ = 17.2 − 20.3 (under the uninformed prior, the posterior lacks power to estimate this parameter, with *L*_fis_ = 17.2 − ∞). These estimates suggest that a transition in the mutational process driving repeat interruption occurs somewhere in the neighborhood of *L* ∼ 20, below which fission is largely a consequence of substitutions, while repeats above this range of lengths are primarily broken up by non-motif insertions.

The intermediate values of *L*^*^ and *L*_fis_ (which sit between the long repeat regime boundary and the truncation length) suggest that, in addition to the separability of long repeat dynamics from the substitution-driven short repeat regime, most long repeats are also *mechanistically independent* from short repeats; the primary mutational mechanisms that alter repeat length may be categorically different from replication and repair pathways that generate substitutions in shorter sequences. In this sense, two distinct boundaries can be placed on the lengths of repetitive sequences, corresponding to up to three distinct regimes: repeats below roughly 10 nucleotides are primarily subject to random substitutions; repeats below *L*= *L*_fis_ ∼ 20, which may experience expansion and contraction, are subject to substitution-based interruptions; repeats longer than *L*_fis_ primarily exhibit repeat instability-based length changes, evolving both dynamically and mechanistically independently from shorter-length repeats subject to substitution-based effects.

---

**Figure SN1:**
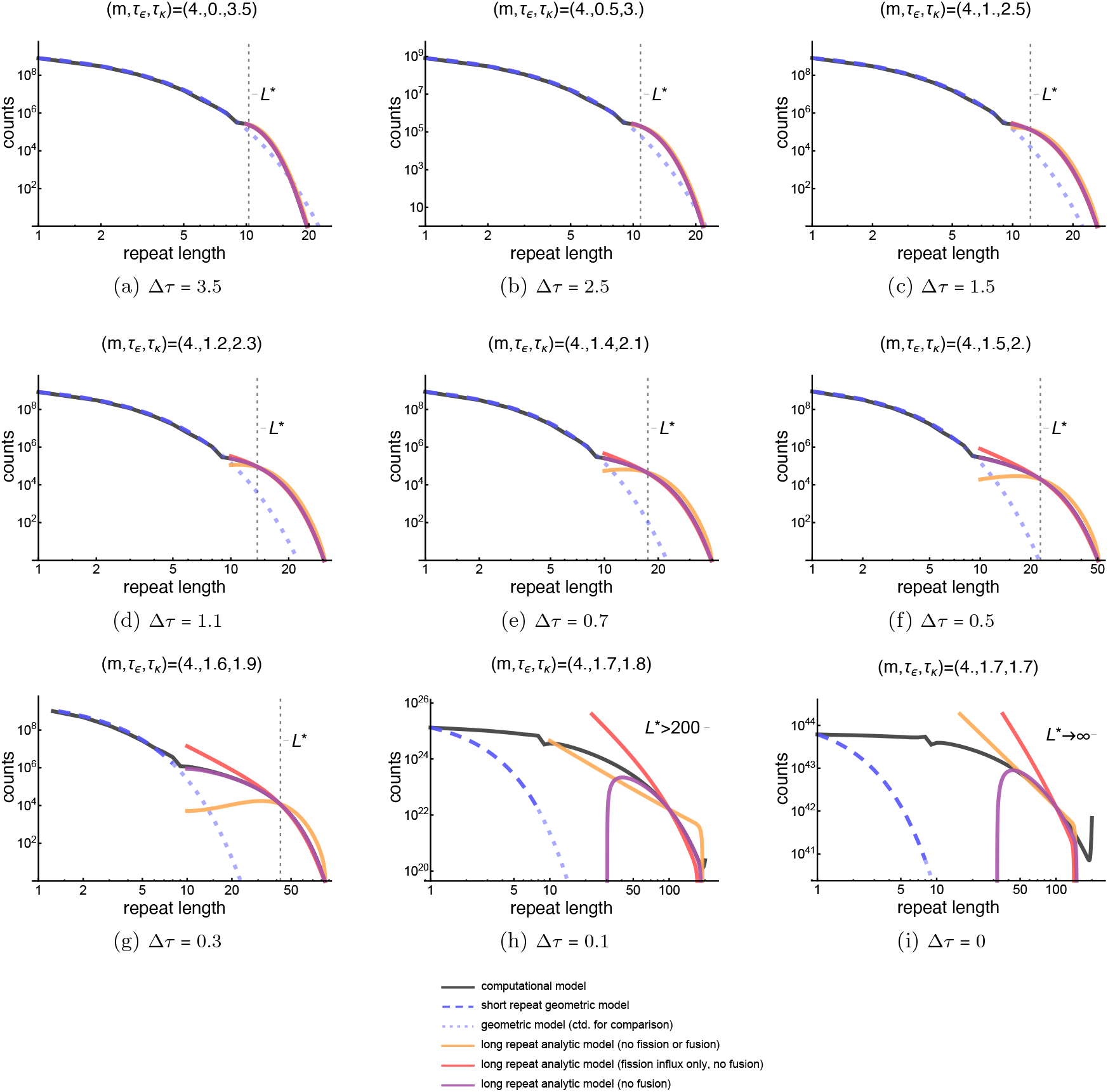
Comparison between computational model results and numerical solutions to steady state equations for m = 4 for parameter values with of constant τ_*ϵ*_ + τ_*κ*_ = 3.5. For contrast, (*τ*_*ϵ*_, *τ*_*κ*_) = (1.7, 1.7) (Δ*τ* = 0; *τ*_*ϵ*_ + *τ*_*κ*_ = 3.4) provides an example of *τ* = *τ* . Each inset shows plots of the computationally modeled distribution at the final time point (black), geometric approximation for shorter repeats of length *L* < 10 (blue, continued as blue dashed line for comparison to distribution tail shape), numerical solutions to Equation SN46 with no fission (orange), numerical solutions to Equation SN45 with fission out but no fission in (red), and numerical solutions to Equation SN51 with fission (purple). (a, b) Comparisons for parameter combinations with Δ*τ* > 1.5 (referred to as Δ*τ* ≫ 1) show good agreement for all numerical solutions; fission is negligible. (c) Boundary between large Δ*τ* and intermediate values where fission out first becomes relevant. (d, e) Fission outflux becomes relevant when Δ*τ* < 1.5. Local approximation (orange) underestimates DRL for *L* < *L*^*^ and overestimates *L* > *L*^*^ (though, approx. remains close near *L*_trunc_). (f, g) Fission influx is relevant for *L* < *L*, while the DRL for *L* > *L*^*^ is well approximated when considering only fission outflux and local dynamics (i.e., treating fission influx as negligible). Under informative prior, Δ*τ* = 0.5 for all parameters in 95% HDR (i.e., panel (f) represents realistic regime for humans; uninformative prior suggests similar values Δ*τ* ∼ 0.4 − 0.6). (h) *L*^*^ lies far above computational grid boundary (*L*_*boundary*_ = 200). DRL would truncate at a length *L* ≫ 200 and stabilize if the grid was extended far beyond realistic lengths. (i) Unstable dynamical regime subject to nonlinear growth. The distribution shows clear interaction with the reflecting boundary.

**Figure SN2:**
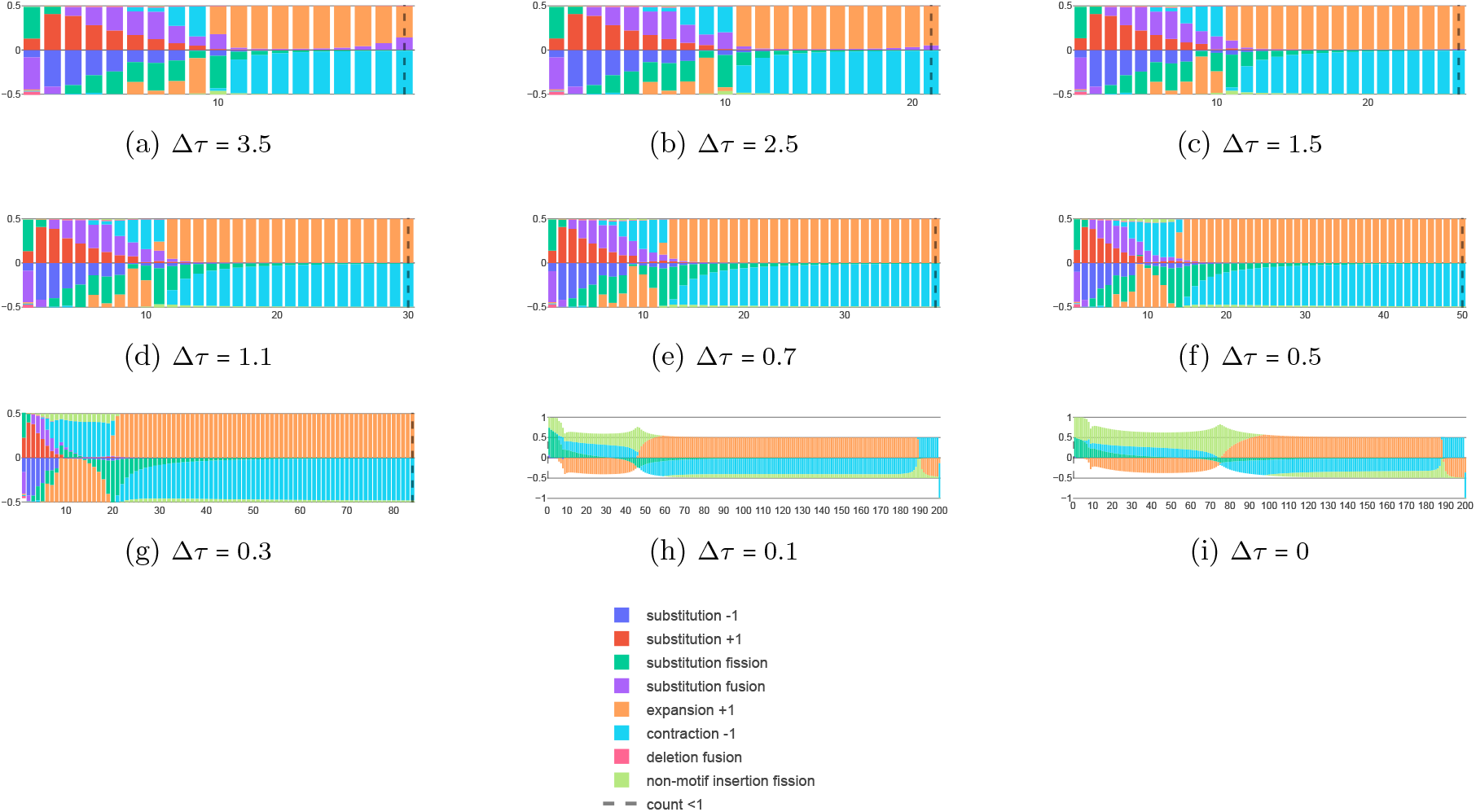
Computational model results for the net flux per mutation type. Subplots (a)-(i) correspond to the same parameters shown in Figures SN1. After the final time point of each run, flux in and out of each bin was calculated and attributed to the following transitions: local length increase due to *µ* substitutions (darker blue); local length decrease due to *ν* substitutions (darker red); nonlocal fission generated by *ν* substitutions (darker green); nonlocal fusion due to *µ* substitutions (purple); local length increase due to expansion (orange); local length decrease due to contraction (lighter blue); nonlocal fusion resulting from deletion of a *B* base (lighter red); nonlocal fission due to non-motif insertions (lighter green). Dashed black line shows longest populated length *L*_trunc_ where *ρ* (*L* > *L*_trunc_) < 1. Net flux per category was computed as flux in minus flux out (i.e., net change in the number of repeats per length class per mutational transition). After computing the net flux for each effect, the sum of magnitudes of all effects was separately normalized at each length (i.e., height of stacked bars sums to one). If a given transition results in a net influx (outflux), associated bar appears above (below) the axis. Bins showing identical heights above and below zero are maintained in detailed balance.

**Figure SN3:**
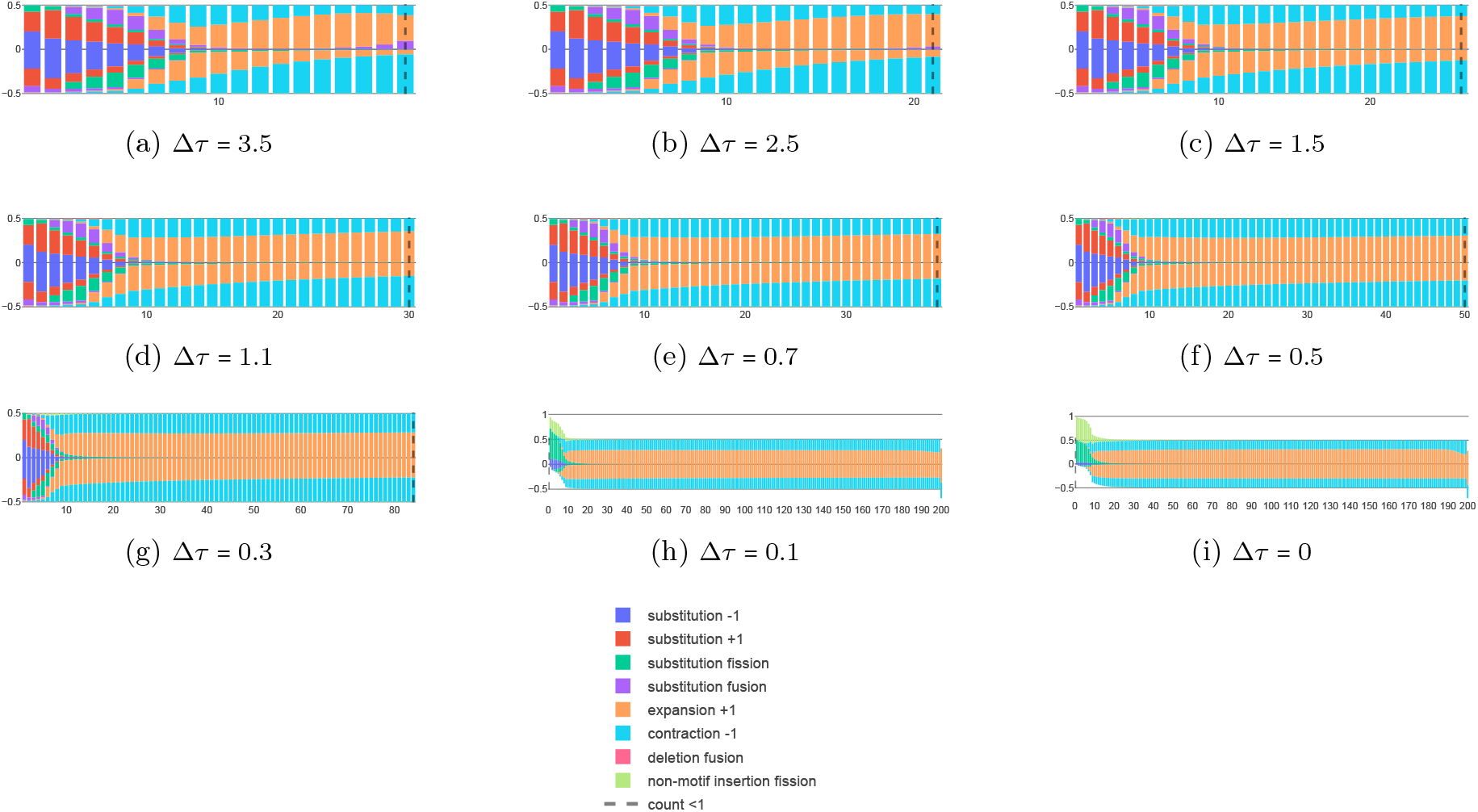
Computational model results for directional flux per mutation type. Subplots (a)-(i) correspond to the same parameters shown in Figures SN1. After the final time point of each run, flux in and out of each bin was calculated and attributed the same mutational processes as in Figure SN2 (shown in corresponding colors). For each category, flux in and flux out are plotted separately at each length (shown above and below zero, respectively). The sum of the magnitudes of all effects (influxes plus outfluxes) was separately normalized to one at each length. Bins showing identical heights above and below zero (i.e., influx equal to outflux) are maintained in detailed balance. In contrast to Figure SN2, each bar height represents the fraction of total number of transitions (in either direction) due to each signed mutational transition (e.g., fraction of number of transitions from expansion influx events, expansion outflux events, contraction influx events, etc.).

**Figure SN4:**
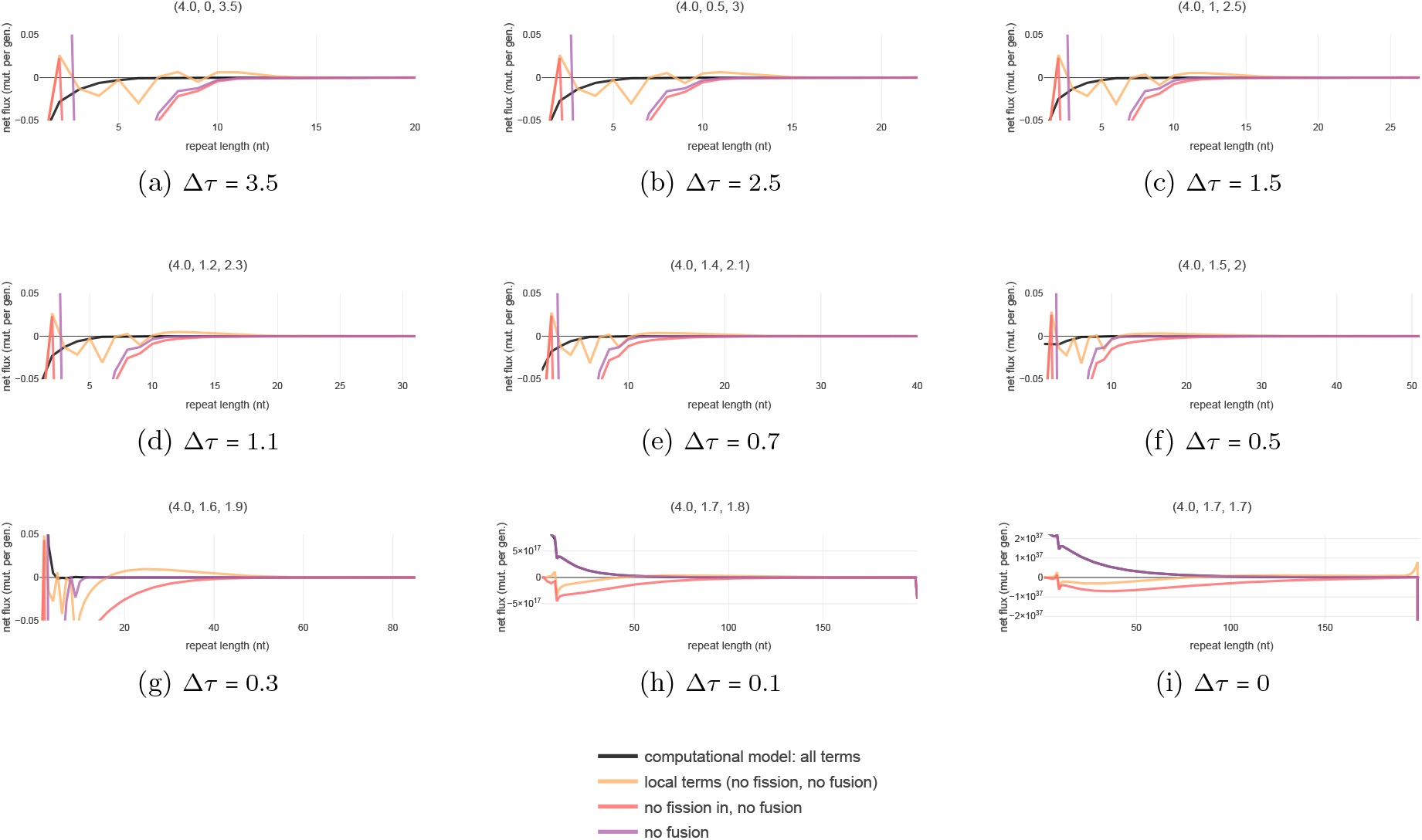
Computational model results for collective non-normalized fluxes showing the relevance of local transitions, fission, and fusion. Subplots (a)-(i) correspond to the same parameters shown in Figures SN1. At the final iterated time point, fluxes were calculated for each length class and summed appropriately to test the accuracy of different analytic models of the long length regime specified in Equations SN37, SN45, and SN46. Each model specifies an approximate steady state equation assembled as a subset of the full collection of terms shown in Equation SN32. Equation SN32, which includes repeat fusion, was summarized by adding the fluxes due to all mutational effects separately at each length (shown in black); detailed balance occurs when all fluxes sum to zero at a given length. Each approximation is deemed appropriate at lengths where they overlap the black curve (restricted to *L* > 10). In contrast to Figure SN1, which tests the accuracy of solutions to the approximated steady state equations, this comparison tests the differential equation more directly by specifying the magnitude of individual terms in the expression (within a given parameter and length regime); in particular, this comparison captures nonlocal effects in Equation SN37 directly, without reference to Equation SN37. The model missing only fusion (Equation SN37; purple) deviates from the full model (Equation SN32; black) only for *L* ≲ 10 indicating fusion is negligible in the long repeat regime. All three approximations overlap for large Δ*τ*, indicating the dominant behavior is local (described by Equation SN46; yellow); the model with fission treated strictly as an outflux (Equation SN45; red) remains a good approximation to the full effects of fission (purple) above roughly Δ*τ* ∼ 0.5. (h, i): The reflective boundary imposed in our computational model generates artefactual spikes in fluxes near *L*_*boundary*_ = 200 due to nonzero counts in these length classes.

**Figure SN5:**
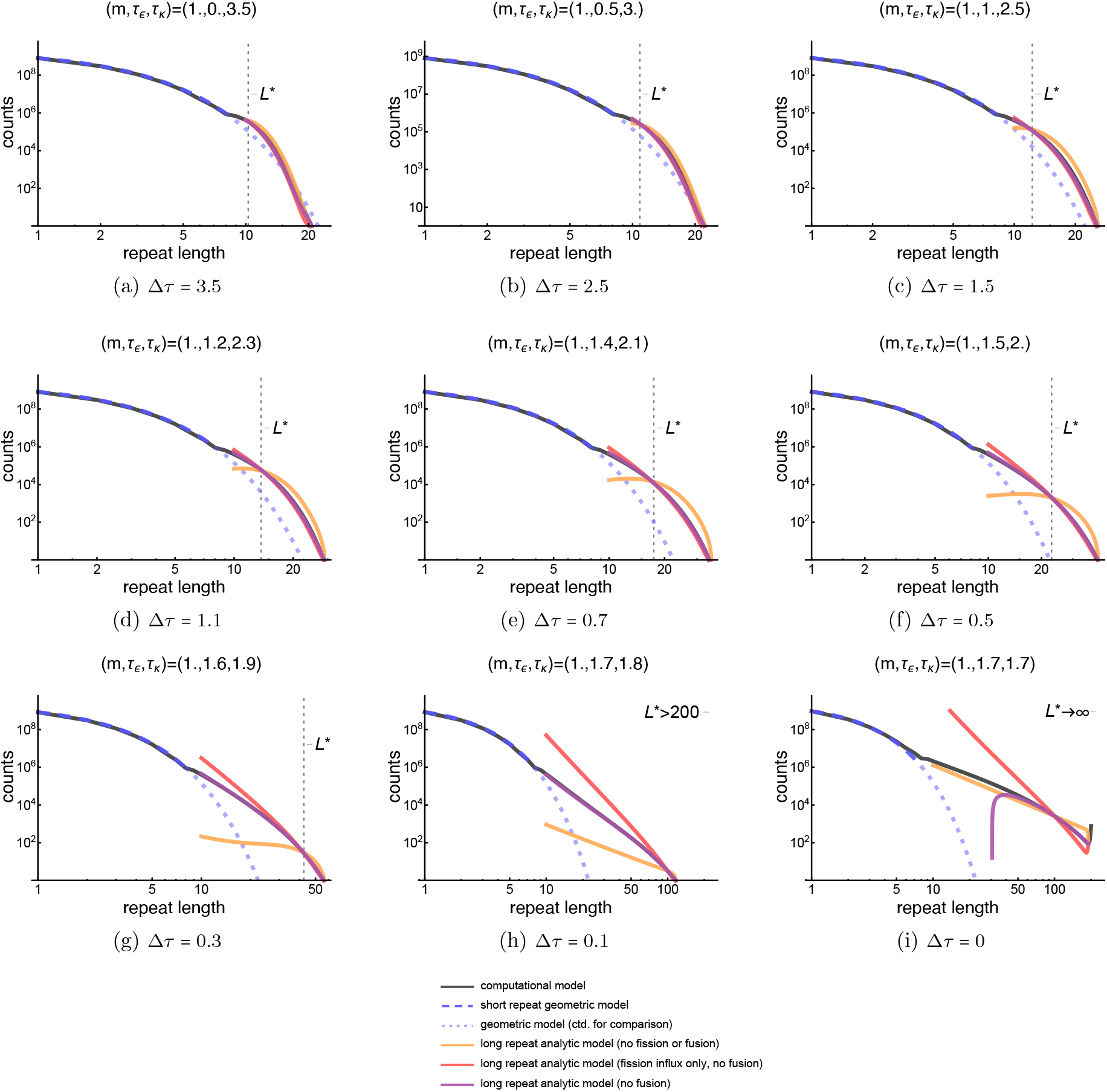
Comparison between computational model results and numerical solutions to steady state equations for m = 1. Computationally modeled distributions are plotted for the same (*τ*_*ϵ*_, *τ*_*κ*_) (and therefore Δ*τ* ) parameters plotted in Figure SN1 (shown as the same inset panel), but for *m* = 1. Each inset shows plots of the computationally modeled distribution at the final time point (black), geometric analytic approximation for shorter repeats of length *L* < 10 (blue, continued as blue dashed line for comparison to distribution tail shape), numerical solutions to Equation SN46 with no repeat fission (orange), numerical solutions to Equation SN45 with fission out but without fission in (red), and numerical solutions to Equation SN51 with fission out and fission in (purple). As expected, the local approximation (orange) deviates at larger Δ*τ* for smaller *m*; substantial deviation can be seen when Δ*τ* ≤ 1.5. The qualitative properties of each approximation are unchanged.

**Figure SN6:**
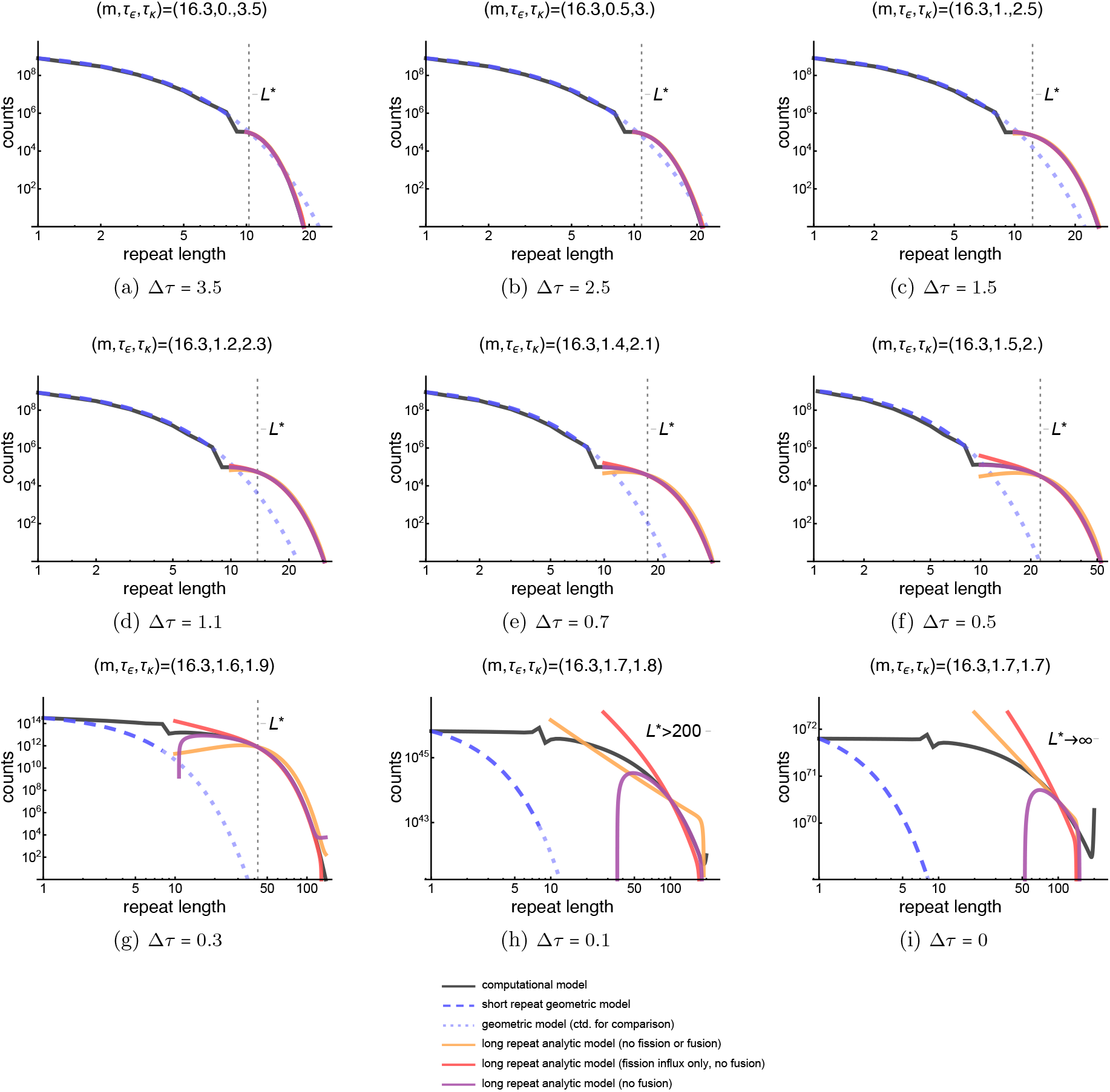
Comparison between computational model results and numerical solutions to steady state equations for m = 16.3. Computationally modeled distributions are plotted for the same (*τ*_*ϵ*_, *τ*_*κ*_ ) (and therefore Δ*τ* ) parameter combinations plotted in Figure SN1 (shown in the same location), but for *m* ≈ 16. Each inset shows plots of the computationally modeled distribution at the final time point (black), geometric analytic approximation for shorter repeats of length *L* < 10 (blue, continued as blue dashed line for comparison to distribution tail shape), numerical solutions to Equation SN46 with no repeat fission (orange), numerical solutions to Equation SN45 with fission out but without fission in (red), and numerical solutions to Equation SN51 with fission out and fission in (purple). As expected, the local approximation (orange) deviates at smaller Δ*τ* for larger *m*; in this case, deviation is only visible when Δ*τ* < 1. The qualitative properties of each approximation are unchanged.

**Figure SN7:**
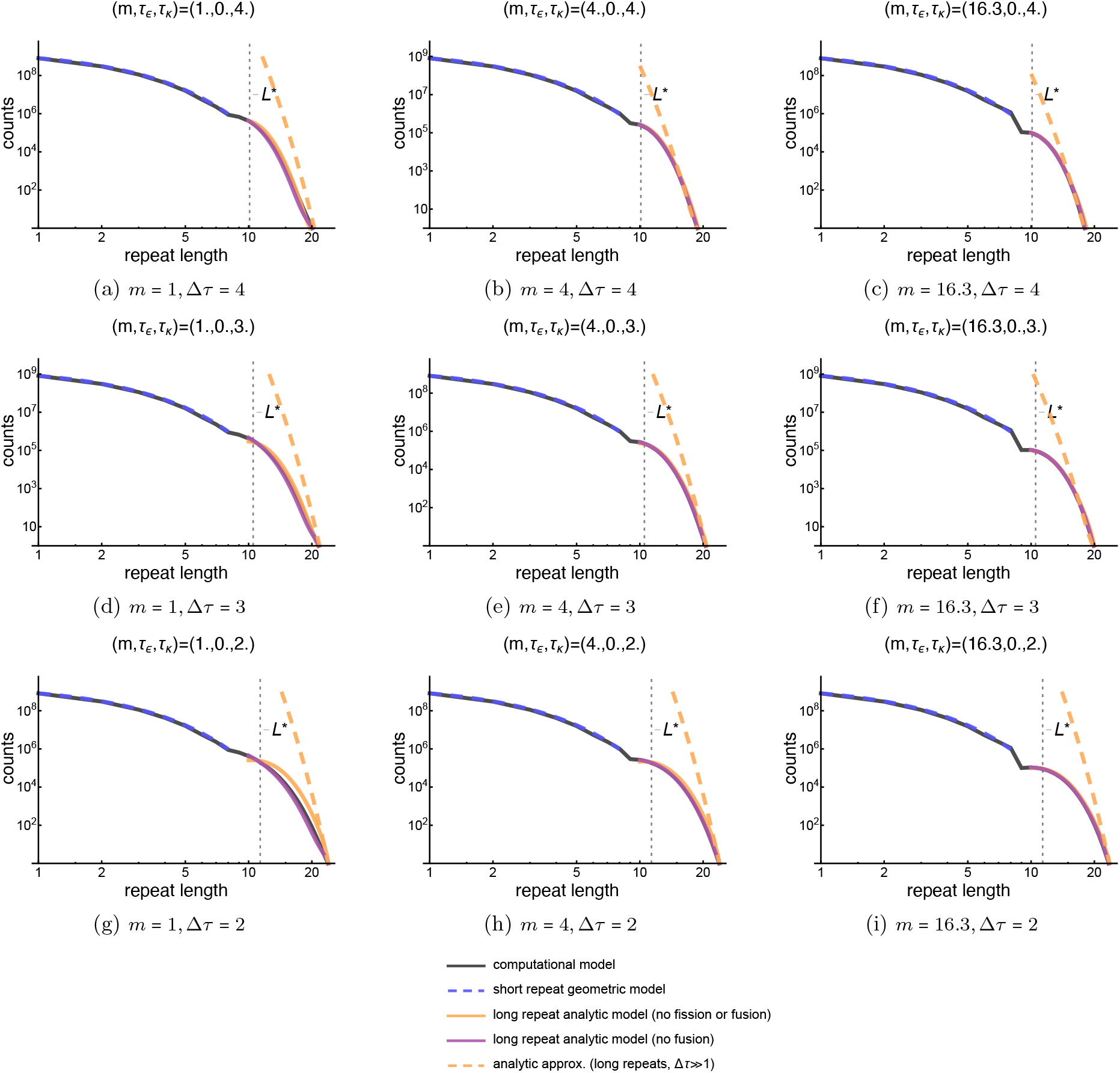
Accuracy of analytic approximation for distribution falloff in the Δτ ≫ 1 regime. The rough approximation in Equation SN48 to the shape of the steady state distribution when Δ*τ* ≫ 1 is shown for a range of parameter combinations with Δ*τ* ≥ 1. Each row plots the same combination of (*τ*_*ϵ*_, *τ*_*κ*_) for multipliers *m* = 1 (left: a, d, g), *m* = 4 (center: b, e, h), and *m* ≈ 16 (right: c, f, i). Each column plots the same value of *m* for Δ*τ* = 4 (top: a-c), Δ*τ* = 3 (middle: d-f), Δ*τ* = 2 (bottom: g-i). For larger Δ*τ* and *m* values, the analytic solution approximates the computationally modeled and numerically generated distributions, except at lengths adjacent to the short repeat regime (roughly 13 > *L* > 10, noting axes are log spaced). At very low *m* and lower Δ*τ*, the local approximation deviates numerically, indicating further analytic approximation used to solve this equation poorly approximates the steady-state DRL.

## References

1 Nurk S, Koren S, Rhie A, et al. The complete sequence of a human genome. Science. 2022;376(6588):44–53. doi:10.1126/science.abj6987

2 Han J, Hsu C, Zhu Z, Longshore JW, Finley WH. Over-representation of the disease associated (CAG) and (CGG) repeats in the human genome. Nucleic Acids Res. 1994;22(9):1735–1740. doi:10.1093/nar/22.9.1735

3 Cox R, Mirkin SM. Characteristic enrichment of DNA repeats in different genomes. Proc Natl Acad Sci U S A. 1997;94(10):5237–5242. doi:10.1073/pnas.94.10.5237

4 Katti MV, Ranjekar PK, Gupta VS. Differential distribution of simple sequence repeats in eukaryotic genome sequences. Mol Biol Evol. 2001;18(7):1161–1167. doi:10.1093/oxfordjournals.molbev.a003903

5 Mirkin SM. Expandable DNA repeats and human disease. Nature. 2007;447(7147):932–940. doi:10.1038/nature05977

6 Depienne C, Mandel JL. 30 years of repeat expansion disorders: What have we learned and what are the remaining challenges?. Am J Hum Genet. 2021;108(5):764–785. doi:10.1016/j.ajhg.2021.03.011

7 Hujoel MLA, Handsaker RE, Kamitaki N, et al. Insights into the causes and consequences of DNA repeat expansions from 700,000 biobank participants. Preprint. bioRxiv. 2024;2024.11.25.625248. Published 2024 Nov 26. doi:10.1101/2024.11.25.625248

8 Khristich AN, Mirkin SM. On the wrong DNA track: Molecular mechanisms of repeat-mediated genome instability. J Biol Chem. 2020;295(13):4134–4170. doi:10.1074/jbc.REV119.007678

9 Wang G, Vasquez KM. Dynamic alternative DNA structures in biology and disease. Nat Rev Genet. 2023;24(4):211–234. doi:10.1038/s41576-022-00539-9

10 Dolzhenko E, Deshpande V, Schlesinger F, et al. ExpansionHunter: a sequence-graph-based tool to analyze variation in short tandem repeat regions. Bioinformatics. 2019;35(22):4754–4756. doi:10.1093/bioinformatics/btz431

11 van Wietmarschen N, Sridharan S, Nathan WJ, et al. Repeat expansions confer WRN dependence in microsatellite-unstable cancers. Nature. 2020;586(7828):292–298. doi:10.1038/s41586-020-2769-8

12 Matos-Rodrigues G, van Wietmarschen N, Wu W, et al. S1-END-seq reveals DNA secondary structures in human cells. Mol Cell. 2022;82(19):3538–3552.e5. doi:10.1016/j.molcel.2022.08.007

13 Erwin GS, Gürsoy G, Al-Abri R, et al. Recurrent repeat expansions in human cancer genomes. Nature. 2023;613(7942):96–102. doi:10.1038/s41586-022-05515-1

14 Bacolla A, Tainer JA, Vasquez KM, Cooper DN. Translocation and deletion breakpoints in cancer genomes are associated with potential non-B DNA-forming sequences. Nucleic Acids Res. 2016;44(12):5673–5688. doi:10.1093/nar/gkw261

15 Grasberger H, Dumitrescu AM, Liao XH, et al. STR mutations on chromosome 15q cause thyrotropin resistance by activating a primate-specific enhancer of MIR7-2/MIR1179. Nat Genet. 2024;56(5):877–888. doi:10.1038/s41588-024-01717-7

16 Kayhanian H, Cross W, van der Horst SEM, et al. Homopolymer switches mediate adaptive mutability in mismatch repair-deficient colorectal cancer. Nat Genet. 2024;56(7):1420–1433. doi:10.1038/s41588-024-01777-9

17 Wright SE, Todd PK. Native functions of short tandem repeats. Elife. 2023;12:e84043. Published 2023 Mar 20. doi:10.7554/eLife.84043

18 Aksenova AY, Mirkin SM. At the Beginning of the End and in the Middle of the Beginning: Structure and Maintenance of Telomeric DNA Repeats and Interstitial Telomeric Sequences. Genes (Basel). 2019;10(2):118. Published 2019 Feb 5. doi:10.3390/genes10020118

19 Cox R, Mirkin SM. Characteristic enrichment of DNA repeats in different genomes. Proc Natl Acad Sci U S A. 1997;94(10):5237–5242. doi:10.1073/pnas.94.10.5237

20 Bell GI, Jurka J. The length distribution of perfect dimer repetitive DNA is consistent with its evolution by an unbiased single-step mutation process. J Mol Evol. 1997;44(4):414–421. doi:10.1007/pl00006161

21 Dechering KJ, Cuelenaere K, Konings RN, Leunissen JA. Distinct frequency-distributions of homopolymeric DNA tracts in different genomes. Nucleic Acids Res. 1998;26(17):4056–4062. doi:10.1093/nar/26.17.4056

22 Field D, Wills C. Abundant microsatellite polymorphism in Saccharomyces cerevisiae, and the different distributions of microsatellites in eight prokaryotes and S. cerevisiae, result from strong mutation pressures and a variety of selective forces. Proc Natl Acad Sci U S A. 1998;95(4):1647–1652. doi:10.1073/pnas.95.4.1647

23 Kruglyak S, Durrett RT, Schug MD, Aquadro CF. Equilibrium distributions of microsatellite repeat length resulting from a balance between slippage events and point mutations. Proc Natl Acad Sci U S A. 1998;95(18):10774–10778. doi:10.1073/pnas.95.18.10774

24 Dokholyan NV, Buldyrev SV, Havlin S, Stanley HE. Distributions of dimeric tandem repeats in non-coding and coding DNA sequences. J Theor Biol. 2000;202(4):273–282. doi:10.1006/jtbi.1999.1052

25 Ellegren H. Microsatellite mutations in the germline: implications for evolutionary inference. Trends Genet. 2000;16(12):551–558. doi:10.1016/s0168-9525(00)02139-9

26 Kruglyak S, Durrett R, Schug MD, Aquadro CF. Distribution and abundance of microsatellites in the yeast genome can Be explained by a balance between slippage events and point mutations. Mol Biol Evol. 2000;17(8):1210–1219. doi:10.1093/oxfordjournals.molbev.a026404

27 Xu X, Peng M, Fang Z. The direction of microsatellite mutations is dependent upon allele length. Nat Genet. 2000;24(4):396–399. doi:10.1038/74238

28 Calabrese PP, Durrett RT, Aquadro CF. Dynamics of microsatellite divergence under stepwise mutation and proportional slippage/point mutation models. Genetics. 2001;159(2):839–852. doi:10.1093/genetics/159.2.839

29 Sibly RM, Whittaker JC, Talbot M. A maximum-likelihood approach to fitting equilibrium models of microsatellite evolution. Mol Biol Evol. 2001;18(3):413–417. doi:10.1093/oxfordjournals.molbev.a003817

30 Dieringer D, Schlötterer C. Two distinct modes of microsatellite mutation processes: evidence from the complete genomic sequences of nine species. Genome Res. 2003;13(10):2242–2251. doi:10.1101/gr.1416703

31 Lai Y, Sun F. The relationship between microsatellite slippage mutation rate and the number of repeat units. Mol Biol Evol. 2003;20(12):2123–2131. doi:10.1093/molbev/msg228

32 Whittaker JC, Harbord RM, Boxall N, Mackay I, Dawson G, Sibly RM. Likelihood-based estimation of microsatellite mutation rates. Genetics. 2003;164(2):781–787. doi:10.1093/genetics/164.2.781

33 Ellegren H. Microsatellites: simple sequences with complex evolution. Nat Rev Genet. 2004;5(6):435–445. doi:10.1038/nrg1348

34 Messer PW, Arndt PF, Lässig M. Solvable sequence evolution models and genomic correlations. Phys Rev Lett. 2005;94(13):138103. doi:10.1103/PhysRevLett.94.138103

35 Eichler EE, Clark RA, She X. An assessment of the sequence gaps: unfinished business in a finished human genome. Nat Rev Genet. 2004;5(5):345–354. doi:10.1038/nrg1322

36 Treangen TJ, Salzberg SL. Repetitive DNA and next-generation sequencing: computational challenges and solutions. Nat Rev Genet. 2011;13(1):36–46. Published 2011 Nov 29. doi:10.1038/nrg3117

37 Kristmundsdóttir S, Sigurpálsdóttir BD, Kehr B, Halldórsson BV. popSTR: population-scale detection of STR variants. Bioinformatics. 2017;33(24):4041–4048. doi:10.1093/bioinformatics/btw568

38 Jeanjean SI, Shen Y, Hardy LM, et al. A detailed analysis of second and third-generation sequencing approaches for accurate length determination of short tandem repeats and homopolymers. Nucleic Acids Res. 2025;53(5):gkaf131. doi:10.1093/nar/gkaf131

39 Zoonomia Consortium. A comparative genomics multitool for scientific discovery and conservation. Nature. 2020;587(7833):240–245. doi:10.1038/s41586-020-2876-6

40 Kelkar YD, Tyekucheva S, Chiaromonte F, Makova KD. The genome-wide determinants of human and chimpanzee microsatellite evolution. Genome Res. 2008;18(1):30–38. doi:10.1101/gr.7113408

41 Pumpernik D, Oblak B, Borstnik B. Replication slippage versus point mutation rates in short tandem repeats of the human genome. Mol Genet Genomics. 2008;279(1):53–61. doi:10.1007/s00438-007-0294-1

42 Molla M, Delcher A, Sunyaev S, Cantor C, Kasif S. Triplet repeat length bias and variation in the human transcriptome. Proc Natl Acad Sci U S A. 2009;106(40):17095–17100. doi:10.1073/pnas.0907112106

43 Payseur BA, Jing P, Haasl RJ. A genomic portrait of human microsatellite variation. Mol Biol Evol. 2011;28(1):303–312. doi:10.1093/molbev/msq198

44 Ananda G, Walsh E, Jacob KD, et al. Distinct mutational behaviors differentiate short tandem repeats from microsatellites in the human genome. Genome Biol Evol. 2013;5(3):606–620. doi:10.1093/gbe/evs116

45 Ananda G, Hile SE, Breski A, et al. Microsatellite interruptions stabilize primate genomes and exist as population-specific single nucleotide polymorphisms within individual human genomes. PLoS Genet. 2014;10(7):e1004498. Published 2014 Jul 17. doi:10.1371/journal.pgen.1004498

46 Kristmundsdottir S, Jonsson H, Hardarson MT, et al. Sequence variants affecting the genome-wide rate of germline microsatellite mutations. Nat Commun. 2023;14(1):3855. Published 2023 Jun 29. doi:10.1038/s41467-023-39547-6

47 Ellegren H. Heterogeneous mutation processes in human microsatellite DNA sequences. Nat Genet. 2000;24(4):400–402. doi:10.1038/74249

48 Mitra I, Huang B, Mousavi N, et al. Patterns of de novo tandem repeat mutations and their role in autism. Nature. 2021;589(7841):246–250. doi:10.1038/s41586-020-03078-7

49 Kunst CB, Warren ST. Cryptic and polar variation of the fragile X repeat could result in predisposing normal alleles. Cell. 1994;77(6):853–861. doi:10.1016/0092-8674(94)90134-1

50 Eichler EE, Holden JJ, Popovich BW, et al. Length of uninterrupted CGG repeats determines instability in the FMR1 gene. Nat Genet. 1994;8(1):88–94. doi:10.1038/ng0994-88

51 Zoghbi HY, Orr HT. Spinocerebellar ataxia type 1. Semin Cell Biol. 1995;6(1):29–35. doi:10.1016/1043-4682(95)90012-8

52 Latham GJ, Coppinger J, Hadd AG, Nolin SL. The role of AGG interruptions in fragile X repeat expansions: a twenty-year perspective. Front Genet. 2014;5:244. Published 2014 Jul 29. doi:10.3389/fgene.2014.00244

53 Santoro M, Masciullo M, Silvestri G, Novelli G, Botta A. Myotonic dystrophy type 1: role of CCG, CTC and CGG interruptions within DMPK alleles in the pathogenesis and molecular diagnosis. Clin Genet. 2017;92(4):355–364. doi:10.1111/cge.12954

54 Wright GEB, Black HF, Collins JA, et al. Interrupting sequence variants and age of onset in Huntington’s disease: clinical implications and emerging therapies. Lancet Neurol. 2020;19(11):930–939. doi:10.1016/S1474-4422(20)30343-4

55 Völker J, Breslauer KJ. How sequence alterations enhance the stability and delay expansion of DNA triplet repeat domains. QRB Discov. 2023;4:e8. Published 2023 Nov 6. doi:10.1017/qrd.2023.6

56 Wilkinson RD. Approximate Bayesian computation (ABC) gives exact results under the assumption of model error. Stat Appl Genet Mol Biol. 2013;12(2):129–141. Published 2013 May 6. doi:10.1515/sagmb-2013-0010

57 Handsaker RE, Kashin S, Reed NM, et al. Long somatic DNA-repeat expansion drives neurodegeneration in Huntington’s disease. Cell. 2025;188(3):623–639.e19. doi:10.1016/j.cell.2024.11.038

58 Rolfsmeier ML, Lahue RS. Stabilizing effects of interruptions on trinucleotide repeat expansions in Saccharomyces cerevisiae. Mol Cell Biol. 2000;20(1):173–180. doi:10.1128/MCB.20.1.173-180.2000

59 Kelkar YD, Eckert KA, Chiaromonte F, Makova KD. A matter of life or death: how microsatellites emerge in and vanish from the human genome. Genome Res. 2011;21(12):2038–2048. doi:10.1101/gr.122937.111

60 Rajan-Babu IS, Dolzhenko E, Eberle MA, Friedman JM. Sequence composition changes in short tandem repeats: heterogeneity, detection, mechanisms and clinical implications. Nat Rev Genet. 2024;25(7):476–499. doi:10.1038/s41576-024-00696-z

61 Doolittle WF, Sapienza C. Selfish genes, the phenotype paradigm and genome evolution. Nature. 1980;284(5757):601–603. doi:10.1038/284601a0

62 Orgel LE, Crick FH. Selfish DNA: the ultimate parasite. Nature. 1980;284(5757):604–607. doi:10.1038/284604a0

63 Werren JH, Nur U, Wu CI. Selfish genetic elements. Trends Ecol Evol. 1988;3(11):297–302. doi:10.1016/0169-5347(88)90105-X

64 Kelkar YD, Strubczewski N, Hile SE, Chiaromonte F, Eckert KA, Makova KD. What is a microsatellite: a computational and experimental definition based upon repeat mutational behavior at A/T and GT/AC repeats. Genome Biol Evol. 2010;2:620–635. doi:10.1093/gbe/evq046

65 Sinden RR, Wells RD. DNA structure, mutations, and human genetic disease. Curr Opin Biotechnol. 1992;3(6):612–622. doi:10.1016/0958-1669(92)90005-4

66 Strand M, Prolla TA, Liskay RM, Petes TD. Destabilization of tracts of simple repetitive DNA in yeast by mutations affecting DNA mismatch repair. Nature. 1993;365(6443):274–276. doi:10.1038/365274a0

67 McMurray CT. Mechanisms of DNA expansion. Chromosoma. 1995;104(1):2–13. doi:10.1007/BF00352220

68 Pearson CE, Sinden RR. Trinucleotide repeat DNA structures: dynamic mutations from dynamic DNA. Curr Opin Struct Biol. 1998;8(3):321–330. doi:10.1016/s0959-440x(98)80065-1

69 Iyer RR, Pluciennik A. DNA Mismatch Repair and its Role in Huntington’s Disease. J Huntingtons Dis. 2021;10(1):75–94. doi:10.3233/JHD-200438

70 Baptiste BA, Ananda G, Strubczewski N, et al. Mature microsatellites: mechanisms underlying dinucleotide microsatellite mutational biases in human cells. G3 (Bethesda). 2013;3(3):451–463. doi:10.1534/g3.112.005173

71 Putnam CD. Strand discrimination in DNA mismatch repair. DNA Repair (Amst). 2021;105:103161. doi:10.1016/j.dnarep.2021.103161

72 Murat P, Guilbaud G, Sale JE. DNA polymerase stalling at structured DNA constrains the expansion of short tandem repeats. Genome Biol. 2020;21(1):209. Published 2020 Aug 21. doi:10.1186/s13059-020-02124-x

73 Habraken Y, Sung P, Prakash L, Prakash S. Binding of insertion/deletion DNA mismatches by the heterodimer of yeast mismatch repair proteins MSH2 and MSH3. Curr Biol. 1996;6(9):1185–1187. doi:10.1016/s0960-9822(02)70686-6

74 Sia EA, Kokoska RJ, Dominska M, Greenwell P, Petes TD. Microsatellite instability in yeast: dependence on repeat unit size and DNA mismatch repair genes. Mol Cell Biol. 1997;17(5):2851–2858. doi:10.1128/MCB.17.5.2851

75 Gendrel CG, Dutreix M. (CA/TG) microsatellite sequences escape the inhibition of recombination by mismatch repair in Saccharomyces cerevisiae. Genetics. 2001;159(4):1539–1545. doi:10.1093/genetics/159.4.1539

76 Jensen LE, Jauert PA, Kirkpatrick DT. The large loop repair and mismatch repair pathways of Saccharomyces cerevisiae act on distinct substrates during meiosis. Genetics. 2005;170(3):1033–1043. doi:10.1534/genetics.104.033670

77 Surtees JA, Alani E. Mismatch repair factor MSH2-MSH3 binds and alters the conformation of branched DNA structures predicted to form during genetic recombination. J Mol Biol. 2006;360(3):523–536. doi:10.1016/j.jmb.2006.05.032

78 Panigrahi GB, Slean MM, Simard JP, Gileadi O, Pearson CE. Isolated short CTG/CAG DNA slip-outs are repaired efficiently by hMutSbeta, but clustered slip-outs are poorly repaired. Proc Natl Acad Sci U S A. 2010;107(28):12593–12598. doi:10.1073/pnas.0909087107

79 Omer S, Lavi B, Mieczkowski PA, Covo S, Hazkani-Covo E. Whole Genome Sequence Analysis of Mutations Accumulated in rad27Δ Yeast Strains with Defects in the Processing of Okazaki Fragments Indicates Template-Switching Events. G3 (Bethesda). 2017;7(11):3775–3787. Published 2017 Nov 6. doi:10.1534/g3.117.300262

80 Tsutakawa SE, Thompson MJ, Arvai AS, et al. Phosphate steering by Flap Endonuclease 1 promotes 5’-flap specificity and incision to prevent genome instability. Nat Commun. 2017;8:15855. Published 2017 Jun 27. doi:10.1038/ncomms15855

81 Bae SH, Bae KH, Kim JA, Seo YS. RPA governs endonuclease switching during processing of Okazaki fragments in eukaryotes. Nature. 2001;412(6845):456–461. doi:10.1038/35086609

82 Deshmukh AL, Porro A, Mohiuddin M, et al. FAN1, a DNA Repair Nuclease, as a Modifier of Repeat Expansion Disorders. J Huntingtons Dis. 2021;10(1):95–122. doi:10.3233/JHD-200448

83 Goldmann JM, Wong WS, Pinelli M, et al. Parent-of-origin-specific signatures of de novo mutations. Nat Genet. 2016;48(8):935–939. doi:10.1038/ng.3597

84 Yuen RK, Merico D, Cao H, et al. Genome-wide characteristics of de novo mutations in autism. NPJ Genom Med. 2016;1:160271–1602710. doi:10.1038/npjgenmed.2016.27

85 Jónsson H, Sulem P, Kehr B, et al. Parental influence on human germline de novo mutations in 1,548 trios from Iceland. Nature. 2017;549(7673):519–522. doi:10.1038/nature24018

86 An JY, Lin K, Zhu L, et al. Genome-wide de novo risk score implicates promoter variation in autism spectrum disorder. Science. 2018;362(6420):eaat6576. doi:10.1126/science.aat6576

87 Halldorsson BV, Palsson G, Stefansson OA, et al. Characterizing mutagenic effects of recombination through a sequence-level genetic map. Science. 2019;363(6425):eaau1043. doi:10.1126/science.aau1043

88 Sasani TA, Pedersen BS, Gao Z, et al. Large, three-generation human families reveal post-zygotic mosaicism and variability in germline mutation accumulation. Elife. 2019;8:e46922. Published 2019 Sep 24. doi:10.7554/eLife.46922

89 Jonsson H, Magnusdottir E, Eggertsson HP, et al. Differences between germline genomes of monozygotic twins. Nat Genet. 2021;53(1):27–34. doi:10.1038/s41588-020-00755-1

90 Goes FS, Pirooznia M, Tehan M, et al. De novo variation in bipolar disorder. Mol Psychiatry. 2021;26(8):4127–4136. doi:10.1038/s41380-019-0611-1

91 McGinty RJ, Sunyaev SR. Revisiting mutagenesis at non-B DNA motifs in the human genome. Nat Struct Mol Biol. 2023;30(4):417–424. doi:10.1038/s41594-023-00936-6

92 Wolfram Research, Inc., Mathematica, Version 14.0, Champaign, IL (2024).

## Supplementary References

1 Kristmundsdottir, S. et al. Sequence variants affecting the genome-wide rate of germline microsatellite mutations. en. Nat Commun 14, 3855 (2023).

2 Wolfram Research, Inc. Mathematica, Version 14.0, Champaign, IL, (2024).

